# Microbial nitrogen removal versus recycling in the redox transition zone of a meromictic lake and its coupling to sulfur

**DOI:** 10.1101/2025.01.03.627289

**Authors:** Jana Tischer, Moritz F. Lehmann, Guangyi Su, Fabio Lepori, Jakob Zopfi

**Affiliations:** Department of Environmental Sciences, University of Basel, Basel, Switzerland; Department for Environment, Constructions and Design, University of Applied Sciences and Arts of Southern Switzerland, Canobbio, Switzerland

## Abstract

Organotrophic denitrification is an important nitrogen (N) removal process in lakes, but alternative N reduction processes such as lithotrophic sulfur (S)-oxidizing denitrification may be greatly underappreciated. We studied the redox transition zone (RTZ) in the meromictic water column of the North Basin of Lake Lugano (Switzerland) to characterize N transformation pathways coupled to the S and carbon (C) cycles. Incubations with ^15^N-labeled and unlabeled nitrate showed low denitrification rates and a general limitation of organic electron donors. The most accessible fractions of exported primary production biomass may have been largely consumed in the oxic water column during sedimentation, and did not reach the RTZ at ∼100 m. Conversely, sulfide (H_2_S) and methane (CH_4_), major end products of anaerobic degradation of the more recalcitrant organic matter fractions in the sediment, represent a continuous source of energy to the RTZ, fostering the establishment of a community of S- and CH_4_-dependent nitrate reducers, dominated by *Sulfuritalea* and *Candidatus* Methylomirabilis over several years of observation. Anoxic incubation experiments with H_2_S amendments revealed a strong stimulation of dissimilatory nitrate reduction to ammonium (DNRA), but not denitrification. High relative abundances of the archaeal ammonia oxidizer *Candidatus* Nitrosopumilus and bacterial nitrifiers indicate intense nitrate regeneration by nitrification in the upper RTZ. The potential interaction between nitrification and S-driven DNRA is unclear. However, their importance in close proximity suggests that, at least under conditions of carbon limitation, N recycling between the nitrate and ammonium pools, predominates over N removal via complete denitrification in the Lake Lugano North Basin.

## Introduction

Due to the widespread use of fertilizers in agriculture and waste water release, large amounts of reactive (i.e., fixed) nitrogen (N) have entered and altered natural ecosystems (Gruber and Galloway 2008). It is estimated that up to 75% of the anthropogenic N introduced into inland waters are removed along the freshwater/seawater continuum before reaching coastal marine ecosystems. In this regard, lakes serve as efficient N sinks, playing a particularly important role in mitigating anthropogenic nutrient loads (Howarth et al. 1996). Canonical organotrophic denitrification, the microbial reduction of nitrate (NO_3_^-^) to dinitrogen gas (N_2_) with organic compounds as substrate, is considered to be the most important N removal process (Seitzinger 1988). However, there is increasing evidence for alternative substrates such as methane (CH_4_) (Raghoebarsing et al. 2006) and inorganic electron donors like reduced sulfur (S) compounds (Hulth et al. 2005; Burgin and Hamilton 2007). Anaerobic ammonium oxidation to N_2_ (anammox) and the dissimilatory nitrate reduction to ammonium (DNRA) represent additional, possibly underappreciated, N turnover processes in lakes (Schubert et al. 2006; Roland et al. 2018). Yet, while these major N-transformations are well known, their relative contributions to N cycling, their different trophic modes (i.e., organotrophic versus lithotrophic), as well as the responsible microbial actors in lakes are not well understood. This is particularly important in the context of lacustrine N budgets: in contrast to denitrification and anammox, DNRA promotes N recycling rather than fixed-N removal.

Denitrification and DNRA have in common, besides the initial reduction step from NO_3_^-^ to NO_2_^-^, that both processes can be performed by organotrophic as well as lithotrophic microorganisms (Pandey et al. 2020). It is assumed that DNRA dominates under conditions with a high availability of organic carbon (C_org_) relative to NO_3_^-^, as e.g., in lake sediments, while denitrification is favored in high-NO_3_^-^ environments (Kelso et al. 1997; Dong et al. 2011). Furthermore, the type and availability of alternative inorganic substrates may also affect the mode of nitrate reduction (Brunet and Garcia-Gil 1996; Cojean et al. 2020). Organotrophic N reduction is performed by a wide variety of ubiquitous facultative or strict anaerobic microorganisms, including many Proteobacteria, as well as some Firmicutes and Actinobacteria (Shapleigh 2013; Pandey et al. 2020). Moreover, a range of obligate and facultative chemolithotrophic bacteria (e.g., species of the families *Rhodocyclaceae*, *Sulfurimonadaceae*, or *Hydrogenophilaceae*) can couple, for example, the oxidation of sulfide (H_2_S), elemental sulfur (S^0^), or thiosulfate (S_2_O_3_^2-^) with the reduction of NO_3_^-^ to NO_2_^-^, and from there to N_2_ gas or to NH_4_^+^ (Zumft 1997; Shao et al. 2010; Pandey et al. 2020). To date, only few specialized microorganisms are known (e.g., *Candidatus* Methylomirabilis) that oxidize CH_4_ anaerobically via NO_3_^-^ or NO_2_^-^ reduction to N_2_ (Raghoebarsing et al. 2006; Yao et al. 2024)

The absolute and relative importance of these different N-turnover modes in the natural environment can be highly variable. In lacustrine water columns, reported N-transformation rates vary by several orders of magnitude (i.e., between <50 nmol N L^-1^ d^-1^ and >10 µmol N L^-1^ d^-1^; Table 1). Alternative modes of suboxic N reaction were shown to contribute substantially to total N_2_ production (sometimes even exceeding rates of canonical organotrophic denitrification), such as S-dependent denitrification in Wintergreen Lake (Burgin et al. 2012), methane-dependent denitrification in several Indian reservoirs (Naqvi et al. 2018), or anammox in Lake Tanganyika (Schubert et al. 2006). Studies that concurrently investigated the rates of denitrification, DNRA, and anammox within the same lake water column are rare. Lake Kivu stands out as an exception, with all three processes confirmed to co-occur there (Roland et al. 2018). Obviously, the ecosystem functioning of lakes in general, e.g., the mitigation of excessive N loading versus internal recycling, will strongly depend on the relative importance of denitrification and anammox on the one hand versus DNRA on the other.

**Table 1.**
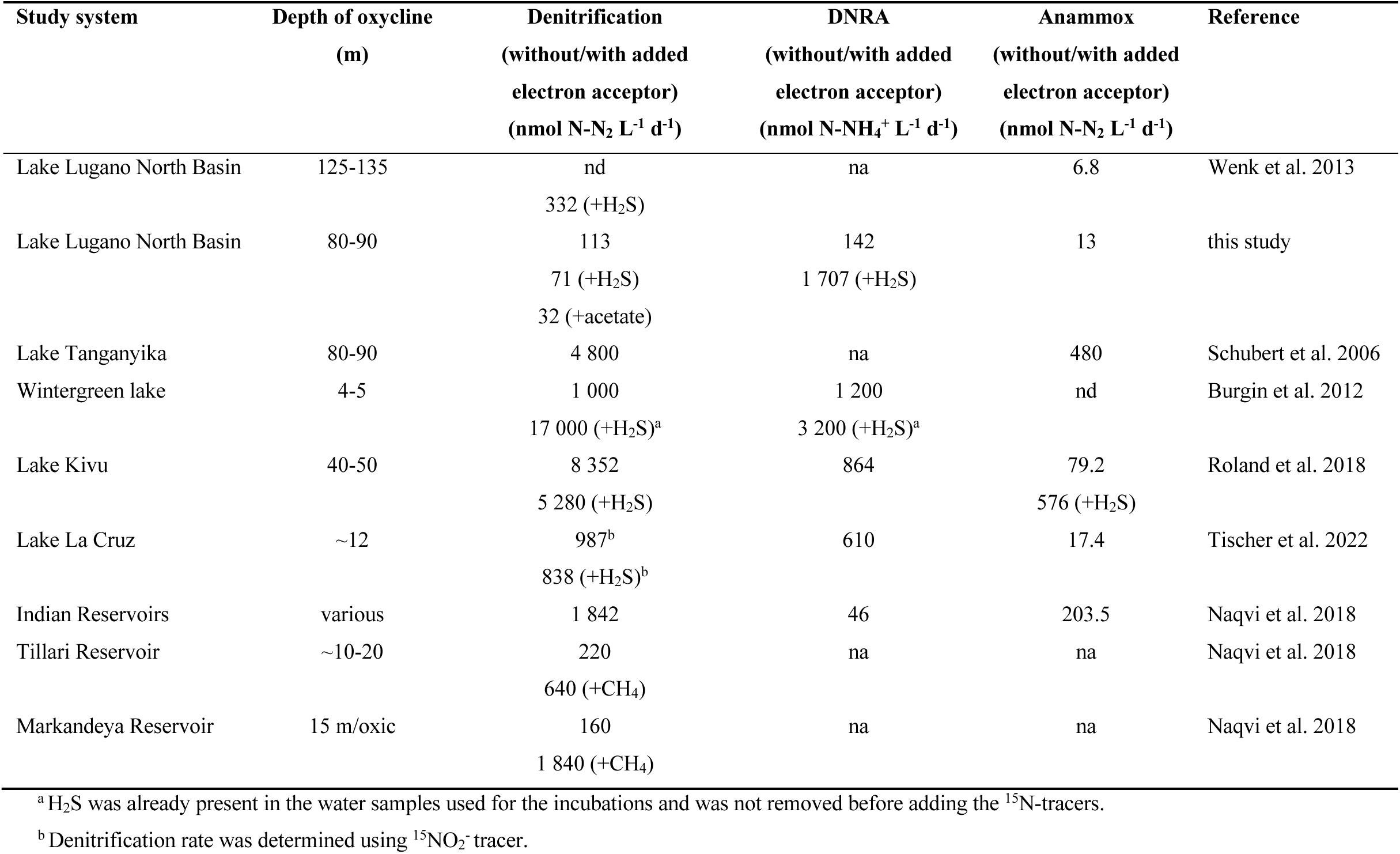
Potential rates of denitrification, with and without additional electron donor (ambient or added), dissimilatory nitrate reduction to ammonium (DNRA), and anammox in the water column of different lakes. From each study, the highest measured rates are reported. Rates of anammox include values determined with different ^15^N-substrates: ^15^NH_4_^+^, ^15^NO_2_^-^, or ^15^NO_3_^-^. nd = not detectable, na = not analyzed.

Lake Lugano has been heavily impacted by increased inputs of phosphorous (P) and N from municipal wastewater, particularly between the 1960s and the 1980s (Barbieri and Mosello 1992; Lepori et al. 2018; Studer et al. 2024). The fluxes of fixed N, and N cycling, in Lake Lugano have been the focus of research for many years (e.g., Barbieri and Mosello 1992; Lehmann et al. 2004). More recently, Wenk et al. (2013; 2014) investigated the modes of N loss in the water column of the northern basin, and reported that canonical organotrophic denitrification in the deep, meromictic northern basin of Lake Lugano plays only a minor role as N sink (Wenk et al. 2013). Instead, S-driven chemolithotrophic denitrification, and to a much lesser extent anammox, were proposed as the main N-removing processes in the North Basin’s redox transition zone (RTZ). However, the conditions that promote fixed-N elimination by lithotrophic (i.e., S-dependent) rather than organotrophic N_2_ production, and whether the controls on the modes of denitrification change seasonally (e.g., due to fluctuations in C_org_ export/substrate availability), remained largely unresolved. Furthermore, it is still uncertain to what extent N recycling occurs within the RTZ of the Lake Lugano North Basin, as possible modes of DNRA have not been determined. Wenk et al. (2014) found, based on their natural abundance N and O isotope measurements, that there is little scope for nitrate regeneration by microaerobic nitrification within the RTZ.

Here we combine ^15^N isotope-label incubation experiments and 16S rRNA gene amplicon sequence data covering several years of observation to i) shed light on the controls on, and importance of, microbial fixed N removal versus fixed N recycling (e.g., by DNRA and nitrifying organisms) within/along the RTZ in the meromictic Lake Lugano North Basin, and ii) identify the main S-oxidizing nitrate reducers, as well as other N- and S-transforming microorganisms involved. We examine whether the previously proposed predominance of S-driven denitrification in the basin’s RTZ is a permanent or a seasonal feature, and we aim to understand why a thermodynamically less favorable inorganic substrate, such as H_2_S, is preferred over organic matter (OM) as driver for N cycling.

## Methods

### Study site

Lake Lugano is a south-alpine lake on the Swiss-Italian border at an altitude of 271 m above sea level. A causeway built on a moraine separates the lake into the southern basin (93 m) and the deep and narrow (288 m) northern basin. Increasing eutrophication led to biogenic meromixis from ∼1960 onward. This state was only interrupted by two mixing events in 2005 and 2006, when cold and windy winters caused complete overturning and the transient oxygenation of the entire water column (Holzner et al. 2009).

### In situ profiling and sample collection

We studied the water column of the northern basin of Lake Lugano at the deepest spot (46°00’37.7”N, 9°01’14.9”E) off the village of Gandria in April, June, and October 2015, March, September, and November 2016, February and October 2017, and April 2018. A conductivity, temperature, depth (CTD) probe (Idronaut Ocean Seven 316Plus) was used to determine oxygen (O_2_) concentrations, temperature, conductivity, and, from July 2016 onward, chlorophyll *a* (Chl *a*). Monthly primary production and nitrate concentration data were obtained through a monitoring program conducted by the University of Applied Sciences and Arts of Southern Switzerland (www.cipais.org).

A RTZ separates the fully oxic upper water column from the anoxic monimolimnion. In this study, the upper RTZ boundary is set at the depth where O_2_ falls below 5 µM, and its lower boundary at the depth, where oxygen-sensitive reduced chemical compounds like Fe^2+^ or H_2_S rise above background levels. Samples were collected across the RTZ starting at around 80 m depth to a maximum depth of 155 to 165 m using 5 L Niskin bottles. Sample water was filled into sterile 1 L plastic or borosilicate bottles for DNA analysis and into 1 L borosilicate bottles for enrichment cultures, each without leaving any headspace. Water samples were kept cold and in the dark until further processing. For DNA analysis water samples (∼1.1-1.2 L) were filtered through 0.2 µm polycarbonate membrane filters (Cyclopore, Whatman) within 24 h of sample collection. For ^15^N-label incubations, water from the Niskin bottle was filled directly into sterile 160 mL serum vials (bubble free with ∼1-2 volumes overflow), and the vials were closed with grey rubber stoppers (VWR). The bottles were kept cold and in the dark at all times, until the incubation experiments were started in the home laboratory within 10 h after sampling. From each depth, additional samples were collected and prepared for various chemical analyses (see below).

### Chemical analyses

#### Nitrogen species

Water samples for the analysis of dissolved inorganic N concentrations were filtered (0.45 µm pore size) right after collection. NH_4_^+^ concentrations were determined using the colorimetrical indophenol reaction (Hansen and Koroleff 1999), and NO_2_^-^ using sulphanilamide and N-(1-Naphthyl)ethylenediamine (Hansen and Koroleff 1999). NO_x_ (i.e., NO_2_^-^ + NO_3_^-^) was determined using a NO_x_-Analyzer (Antek Model 745) involving the reduction of NO_x_ in a hot acidic V^3+^ solution to NO gas, and subsequent chemiluminescence detection (Braman and Hendrix 1989). We determined NO_3_^−^ concentrations by subtracting NO_2_^−^ from NO_x_.

#### Iron

Aliquots of unfiltered and filtered (0.2 µm) water samples were fixed with ∼150 mM (final concentration) nitric acid (HNO_3_) for the analysis of total and dissolved iron concentrations, respectively. Concentrations were quantified by inductively coupled plasma optical emission spectrometry (ICP-OES, Agilent Technologies) with a detection limit of ∼1.8 µM and 5-10% measurement uncertainty. We calculated concentrations of particulate Fe from the difference between total and dissolved Fe.

#### Sulfur compounds

We analyzed concentrations of dissolved H_2_S, S_2_O_3_^2-^, and SO_3_^2-^ according to Zopfi et al. (2008). Briefly, 450 µL unfiltered water samples were fixed in a mixture of 25 µM of HEPES-EDTA buffer (pH 8, 500 mM, 50 mM) and 25 µL of ∼45 mM monobromobimane in the dark. After 30 min, we stopped the derivatization reaction by adding 50 µL of 312 mM methanesulfonic acid. The samples were stored at -20 °C until analysis with reversed-phase high-performance liquid chromatography (RP-HPLC, Dionex) using a LiChrosphere 60RP select B column (125×4 mm, 5 μm; Merck) and a Waters 470 scanning fluorescence detector (excitation at 380 nm; detection at 480 nm). In certain cases, H_2_S concentrations were determined photometrically through the methylene blue reaction in unfiltered samples, immediately fixed with zinc acetate (0.5% final concentration, w/v) upon collection (Cline 1969). For the analysis of suspended S^0^, we filtered 60 mL of water sample through a glass microfiber (GF/F) filter (Whatman), and subsequently stabilized the S^0^ on the filter with 2-3 mL 5% (w/v) zinc acetate solution. The filters were stored at -20 °C until analysis. Using 2 mL of HPLC grade methanol, S^0^ was extracted overnight from the filters and subsequently quantified by RP-HPLC using a Knauer C18-column (Eurospher II, 100-5 C18 H, 125x4 mm) and UV-detection at 265 nm (Zopfi et al. 2008). SO_4_^2-^ concentrations were analyzed by ion chromatography and UV detection (940 Professional IC Vario, Metrohm).

#### Methane

We analyzed methane concentrations in November 2016 and October 2017, as described in Su et al. (2023). Briefly, water samples were collected in 120 mL serum bottles, crimp-sealed with buthyl rubber stoppers, and a 20 mL air headspace was created before fixing the sample with 5 mL of 12.5 M NaOH. Methane concentrations were measured using a gas chromatograph (SRI 8610C, SRI Instruments) with a flame ionization detector (Lehmann et al. 2004).

### Water column stability and turbulent flux calculations

The static stability of the water column was calculated as the Brunt-Väisälä frequency N^2^ (Wüest et al. 1992):

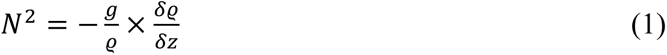

where g represents the gravitational acceleration (9.81 m/s^2^), ρ is the density of water, and δρ/δz denotes the density gradient over a specific depth increment. We calculated the pure water density as function of the measured temperature (T) with the following equation:

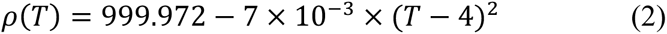

In a second step, we calculated the conductivity-dependent density as function of conductivity according to Wüest et al. (1992):

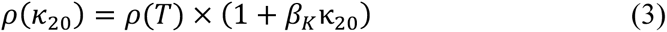

where the constant β_20_ is the relative change of density per unit electrical conductivity (7.05 x 10^-7^ [µS x cm^-1^]^-1^) and κ_20_ is the conductivity at 20°C. Turbulent diffusive fluxes of NO_3_^-^, NH_4_^+^, H_2_S, and CH_4_ towards the RTZ were determined according to Wenk et al. (2013):

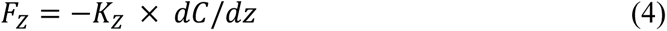

where *F_z_* represents the vertical solute flux, *K_z_* the vertical eddy diffusivity, and *dC/dz* the concentration gradient of the respective solute. Vertical eddy diffusivity was calculated as:

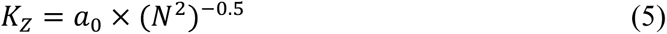

where a_0_ is a system-specific constant, which we adapted from Wenk et al. (2013) for the Lake Lugano North Basin (0.00014 cm^2^ s^-2^). The N^2^ and the *K_z_* values were calculated using the average CTD data integrated over 5 m intervals. To estimate NO_3_^-^ fluxes towards the RTZ from above, we used the average *K_z_* value from all samplings between 2015 and 2018 of 0.85 ± 0.10 m^2^ d^-1^ (standard deviation, SD) for the interval 80-110 m. Similarly, to calculate fluxes of NH_4_^+^, H_2_S, and CH_4_ from the deeper hypolimnion towards the RTZ, the average *K_z_* value for the depth interval 95-150 m (1.31 ± 0.22 SD m^2^ d^-1^) was used. These values for *K_z_* are within the range of previously reported vertical eddy diffusivities for the North Basin of Lake Lugano (Wüest et al. 1992; Wenk et al. 2013).

### DNA extraction, PCR amplification, Illumina sequencing, and data analysis

Filters with collected DNA were stored at -70 °C until extraction of DNA using the Fast DNA Spin Kit for Soil (MP Biomedicals). In addition to the 2015-2018 samples, we used samples already collected in 2009 and 2010 (see Su et al. 2023). The 16S rRNA gene library preparation, amplicon sequencing, and bioinformatic treatment of raw sequences are described in detail in Su et al. (2023). Briefly, the updated Earth Microbiome PCR primers for bacteria and archaea 515F-Y and 926R, targeting the V4 and V5 regions of the 16S rRNA gene, were used for the first PCR (Parada et al. 2015). Sample indices and Illumina adaptors were added by the second PCR before sequencing the purified amplicons on an Illumina MiSeq platform at the Genomics Facility Basel. Amplicon Sequence Variants (ASVs) were identified by denoising amplicons to zero-radius OTUs using the UNOISE algorithm in USEARCH v10.0.240 (Edgar 2010, 2013). Finally, we used SINTAX (Edgar 2016) and the SILVA 16S rRNA reference database v138 (Quast et al. 2013) for the taxonomic assignment of ASVs. We performed downstream sequence analysis and visualization in R v4.1.1 using the packages phyloseq v1.36.0 (McMurdie and Holmes 2013), vegan v2.5-7 (Oksanen et al. 2020), ggplot2 v3.3.5 (Wickham 2016), and dplyr v1.0.7 (Wickham et al. 2021). The sequence data were cleaned by removing mitochondrial and chloroplast sequences, as well as ASVs with unknown phylum-level taxonomy. We examined alpha diversity measures, specifically the observed ASV richness and Shannon diversity, using rarefied data (Su et al. 2023). In addition, we analyzed the temporal evolution of alpha diversity in each redox zone by determining Pearson correlation coefficients of the alpha diversity measure Observed using months as measure of time between sampling timepoints. Beta diversity was assessed using weighted UniFrac distances and principal coordinate analysis (PCoA). Based on the environmental conditions, we divided the samples into three groups: the oxic water column above the RTZ (“oxic”), the RTZ (“RTZ”), and the anoxic water layer below the RTZ (“anoxic”). Pairwise t-tests were performed by using the t.test() function in R to test for significance between the three different redox zones of the alpha diversity measures and of the weighted UniFrac distances by comparing the beta diversity distance pairs within the oxic zone samples, within the RTZ samples and within the anoxic samples. Permutational Multivariate Analysis of Variance (PERMANOVA) was performed using the adonis2() function in vegan to test for significant differences between the different redox zones and/or between timepoints, as well as well their interaction.

We visualized key phyla and classes based on relative abundances of 16S rRNA gene amplicons (taxa with a relative abundance of ≥1% in at least one of the samples). In addition, the temporal change of the microbial community at a given redox zone was monitored by selecting one sample per timepoint, respectively, from the oxic water column (5-10 m above the upper boundary of the RTZ), one from the RTZ (middle of the RTZ), and one from the anoxic water column (40-45 m below the lower boundary of the RTZ). We screened for S- and N-cycling taxonomic groups (Table S1) including ammonia-oxidizing archaea (Yang et al. 2021), ammonia-oxidizing bacteria (Kowalchuk and Stephen 2001), nitrite-oxidizing bacteria (Daims et al. 2016), anammox bacteria (Jetten et al. 2009), organotrophic nitrate/nitrite reducers (Zumft 1997; Pandey et al. 2020), S-oxidizing N reducers (Shao et al. 2010; Kojima and Fukui 2011), CH_4_-oxidizing denitrifiers (Kits et al. 2015), and S-reducers (Rosenberg et al. 2014). Additionally, we selected taxa with names that designate one of the specific before-mentioned metabolisms, e.g., “denitrificans”, “sulfuri”, or “desulf”. In our screening efforts, we also included genera that appeared to thrive and enrich in our incubation experiments with added NO_3_^-^, i.e., that showed a higher (≥2%) relative abundance (ΔRA = RA_end_ – RA_initial_) after incubation (limiting ourselves to microorganisms known for N reduction, denitrification, or DNRA genes; identified using www.kegg.jp) (Tischer et al. 2024/in prep). We only considered taxa with an abundance of ≥0.005% in at least one of the in-situ water-column samples.

### Batch incubation experiments and microbial enrichments

We performed batch incubations with lake water sampled between 2016 and 2018 in order to investigate NO_3_^-^ reduction with different electron donors. In 1-L borosilicate bottles with water samples, we introduced a N_2_ headspace (∼130 mL), and purged for 30 minutes with N_2_ to ensure anoxic incubation conditions. Subsequently, we added NO_3_^-^, NO_3_^-^ and H_2_S, or NO_3_^-^ and sodium acetate (NaAc) from sterile anoxic stock solutions. Targeted initial concentrations for NO_3_^-^ and NaAc were 25 µM, for H_2_S they ranged between 25 and 100 µM (Table 2). One bottle was left as a live control treatment without any amendments, and we incubated a dead control with NO_3_^-^ and H_2_S, killed with 15 mL 50% (w/v) ZnCl_2_, to ensure that no abiotic H_2_S oxidation was taking place (data not shown). Incubations were carried out at ∼8 °C in the dark with gentle agitation. We monitored the concentrations of dissolved N and S species by taking subsamples (∼10 mL) at specific time intervals. To prevent contamination with O_2_, we initially pressurized the incubation bottles to around 2 bars and used N_2_-flushed syringes for sampling. When H_2_S was consumed before the complete reduction of NO_3_^-^, we added additional H_2_S (∼25 µM final concentration). At the end of the experiment, we filtered the remaining water (approximately 300-800 mL) to collect biomass for DNA extraction and sequencing, as described above. This allowed us to identify microbial taxa that responded positively to the different substrate additions. We calculated NO_3_^-^ turnover rates based on the change in NO_3_^-^ concentration over time, using samples from two intervals, i) the first 2-3 days and ii) the first 4-5 days of incubation. We tested differences between treatments with a pairwise t-test and differences between intervals with an analysis of variance (ANOVA). Similarly, we calculated turnover rates for H_2_S, S^0^, SO_3_^2-^, S_2_O_3_^2-^, and SO_4_^2-^ using samples collected during the first 3-4 days of incubation. A *t*-test was performed on the rates to test for significance.

**Table 2.**
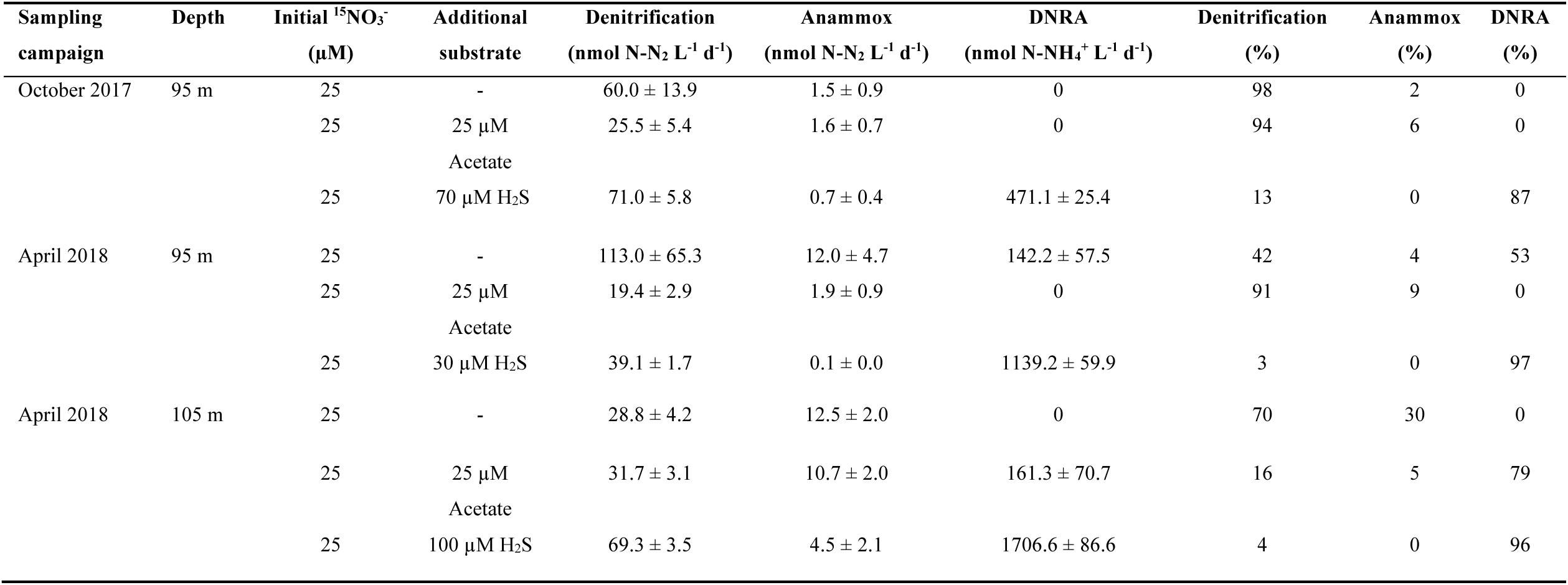
Incubation conditions and potential rates determined in ^15^N-tracer incubation experiments. Values are presented with standard error of means for triplicate or quintuplicate incubations. Percentages refer to the relative contribution of a given process with respect to the combined production of ^15^N-N_2_ and ^15^N-NH_4_^+^.

### ^15^N-label incubation experiments

The ^15^N-label incubation experiments were performed in October 2017 and April 2018 to determine rates of denitrification, anammox, and DNRA at different depths in the water column. We used a modified version of the protocol described in Wenk et al. (2013), adjusted for the additional quantification of ^15^NH_4_^+^. We introduced a 10 mL helium (He) headspace to the 160 mL sample vials, and purged for 10 minutes with He to remove potential traces of O_2_. Subsequently, we added ^15^N-NO_3_^-^ tracer from a sterile and anoxic stock solution of 7.5 mM ^15^N-KNO_3_ (99% ^15^N-KNO_3_, Cambridge Isotope Laboratories, Inc.), aiming for an initial ^15^NO_3_^-^ concentration of 25 µM. Additionally, H_2_S or Na-acetate (each 25 µM final concentration) was added in the respective treatments (Table 2). All incubations were performed in triplicate in October 2017 and in quintuplicate in April 2018. Incubations were performed in the dark, at ∼8 °C with gentle agitation. Samples for N_2_ isotope analysis were collected at five timepoints (∼12, 36, 60, 108, and 325 h) by sampling 2 mL of headspace, in exchange with He, using an airtight 3 mL Luer-lock plastic syringe (Braun) with a plastic valve flushed 3-times with He. The samples were stored in 3 mL exetainers after replacing sterile anoxic water with the gas sample. In addition to headspace samples, we took 5-mL liquid samples, filtered them using 0.2 µm membrane filters, and stored them frozen until analysis. The resulting negative pressure in the incubation vials was equilibrated by adding He. We determined ^15^N-N_2_ production (m/z 29/28 and 30/28 ratios) via isotope ratio mass spectrometry (IRMS; Flash-EA-ConfloIV-DELTA V Advanced, Thermo Scientific), and calculated the rates of denitrification and anammox according to the isotope pairing equations of Thamdrup and Dalsgaard (2002) and Thamdrup et al. (2006), where we used only the first three timepoints. The ^15^N-NH_4_^+^ samples were transformed to ^15^N_2_ by oxidation with hyprobromite (Risgaard-Petersen et al. 1995), and analyzed as described above. NH_4_^+^ standards of different concentrations and ^15^N/^14^N ratios were included in the analysis. We referenced the samples against air N_2_ and used the quantified ammonium-derived ^15^N_2_ to assess ^15^N-NH_4_^+^ concentrations in the liquid sample according to Cojean et al. (2020). DNRA rates were then calculated from the slope of the increase in ^15^N-NH_4_^+^ concentration over time. Given that anammox rates were low (Table 2), we assumed that the consumption of ^15^N-NH_4_^+^ produced by DNRA was negligible.

## Results

### Water column characteristics and hydrochemistry

Water column CTD profiles from the different sampling campaigns are shown in Fig. S1. Water column stability (Fig. S1e) was generally strongest in the subsurface waters, where temperature and density gradients were highest; no significant density gradient was observed at the RTZ (Fig. S1d). Water column O_2_ profiles indicate that all O_2_ was consumed approximately at a water depth of ∼80 and 90 m (four selected timepoints in Fig. 1), which was 35-45 m higher up in the water column compared to 2009 and 2010 (Wenk et al. 2013).

**Figure 1.**
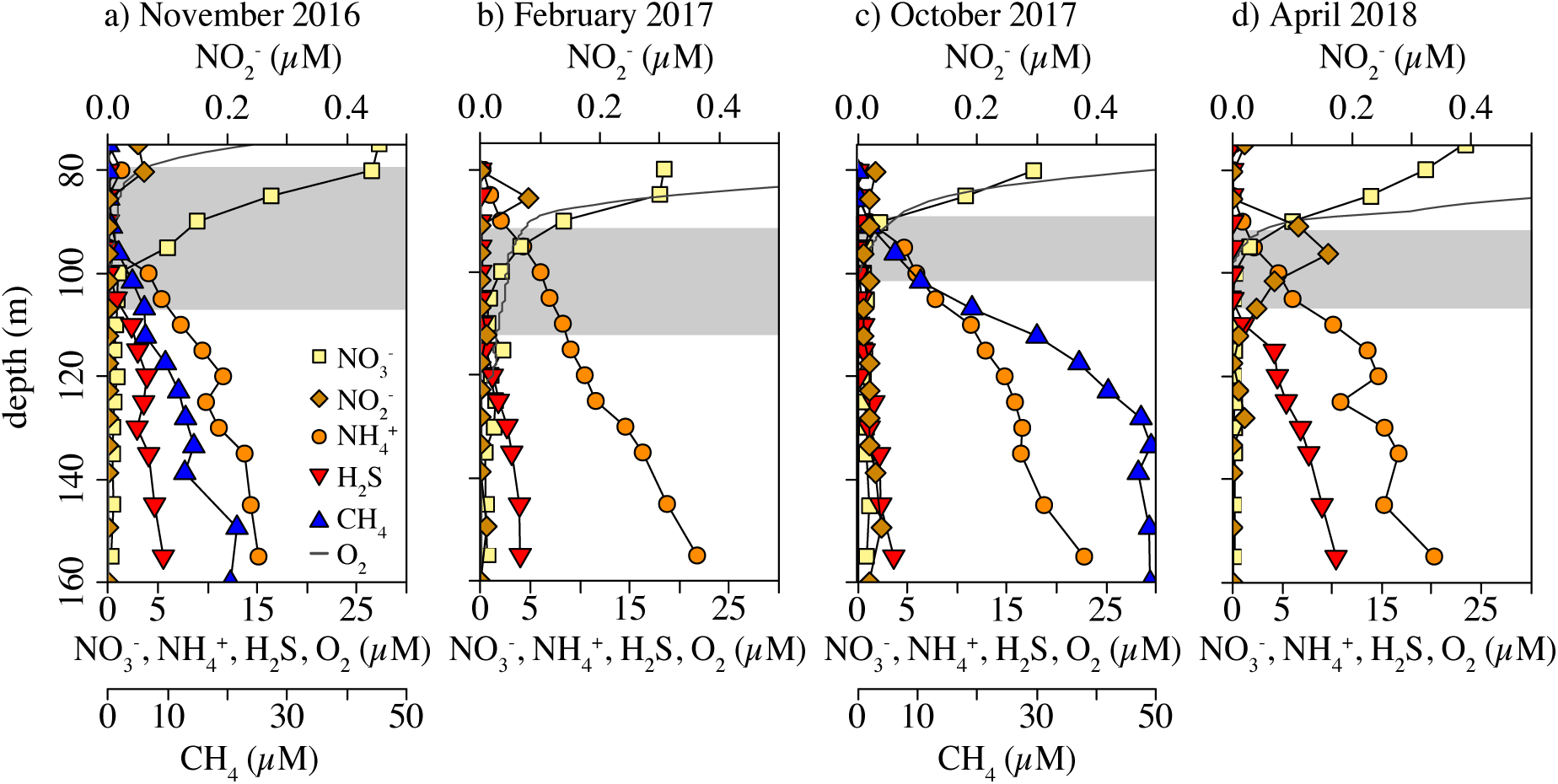
Concentration profiles at the water column redox transition zone (RTZ) of the Lake Lugano North Basin from four different sampling campaigns. The grey bars indicate the extension of the RTZ.

The biweekly O_2_ monitoring data from 2015 to 2018 showed highest concentrations between 0 and 20 m during spring and summer, while in late summer and autumn, the water column was generally less oxygenated with an O_2_ depletion at ∼20 to 30 m (Fig. S2a). Among the sampling dates when we collected water samples for incubation experiments, April 2018 was the only date with a distinct O_2_ peak (up to ∼500 µM at ∼15 m) in shallow waters (Fig. S1a). The Chl *a* concentrations in the surface waters were highest in spring and summer (Fig. S2b). The NO_3_^-^ concentration profiles of the entire water column showed highest concentrations between 20 and 50 m depth, ranging from 24 to 38 µM (Fig. S2b). Detailed concentration profiles at the RTZ (Fig. 1, Fig. S3) indicate that NO_3_^-^ decreased to <0.5 µM within the RTZ. For most timepoints, NO_2_^-^ did not exceed 0.05 µM in the RTZ. NH_4_^+^ concentrations started to increase around the depth of O_2_ depletion and reached concentrations between 15 and 25 µM at ∼155 m. NH_4_^+^ and NO_3_^-^ profiles generally overlapped, with concentrations of ∼1-2 µM each. H_2_S, on the other hand, was first detected approximately 15 m below the depth where NO_3_^-^reached its lowest levels <0.5 µM (Fig. 1, Fig. 2a-c), whereas S^0^, the most abundant product among the intermediates of H_2_S oxidation, was detectable up to the upper border of the RTZ at levels up to ∼0.25 µM (Fig. 2d-f). Low SO_3_^2-^ and S_2_O_3_^2-^ concentrations of <0.15 µM were detected in most of the anoxic water samples (Fig. 2d-f). In this regard, the October 2017 profiles stand out, as S^0^, S_2_O_3_^2-^, and SO_3_^2-^ consistently showed two distinct concentration peaks in the anoxic water column. Dissolved Fe^2+^ concentrations rose up to ∼3 µM around 160 m depth, while particulate Fe remained below 1 µM, reaching a maximum at 125-130 m in October 2017 and April 2018 (Fig. S3e). CH_4_ concentration profiles (November 2016 data have been previously published in Su et al. 2023; Fig. 1) clearly overlapped with NO_3_^-^ profiles. Below the RTZ, CH_4_ concentrations increased to up to ∼50 µM (e.g., October 2017). Turbulent diffusive solute fluxes towards the RTZ between 2015 to 2018 were on average 828 ± 315 SD µmol for NO_3_^-^ m^-2^ d^-1^, 379 ± 122 SD µmol for NH_4_^+^ m^-2^ d^-1^, 199 ± 87 SD µmol for H_2_S m^-2^ d^-1^, and 882 ± 597 SD µmol for CH_4_ m^-2^ d^-1^ (Table S2).

**Figure 2.**
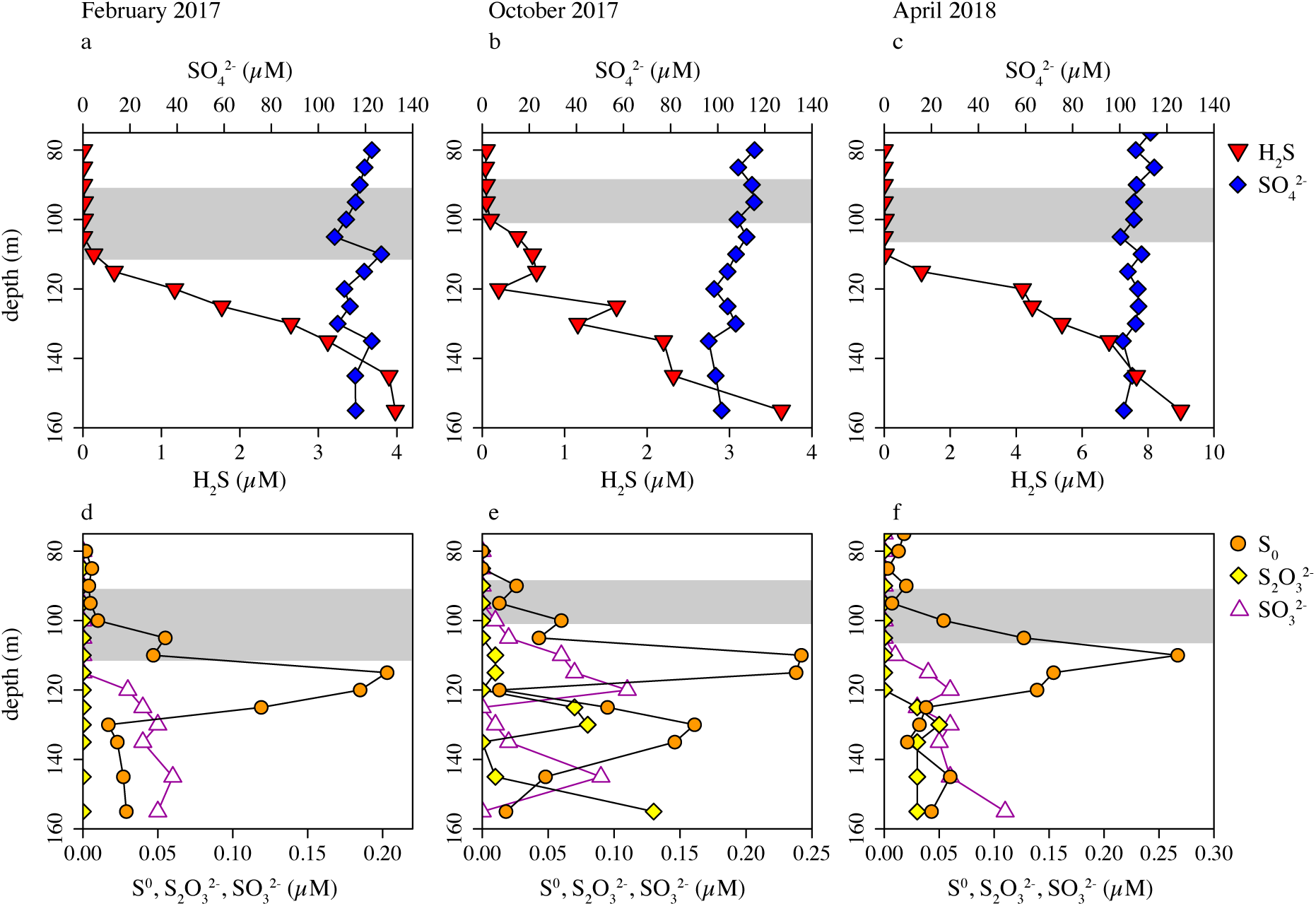
Depth profiles of sulfur compounds across and below the redox transition zone (RTZ) (grey bars), from February 2017 **(a+d)**, October 2017 (**b+e)**, and April 2018 **(c+f)**.

### Microbial community structures

The 16S rRNA gene sequencing revealed differences between the microbial communities in the oxic water column just above the RTZ, within the RTZ, and in the anoxic water below the RTZ with regard to their alpha and beta diversities (Fig. 3). The observed ASV richness was significantly higher in the anoxic water layers below the RTZ compared to the oxic water (pairwise t-test; t = -6.1, df = 19, *p* < 0.001) layer and the RTZ (pairwise t-test; t = -14.2, df = 93, *p* < 0.001). Interestingly, ASV richness increased with time during the multiannual survey of microbial community structures in the water column (Fig. S4), especially in the anoxic water below the RTZ (Pearson correlation coefficient; t = 3.0, n= 68, r = 0.34). The Shannon diversity index, which quantifies both richness and evenness, was higher in the oxic water compared to the RTZ (pairwise t-test; t = 4.2, df = 26, *p* < 0.001) and the anoxic water layer (pairwise t-test; t = 4.4, df = 19, *p* < 0.001). The community structure in the oxic water column seems to be more evenly distributed than in the RTZ and the anoxic zone. Particularly the communities in the anoxic layer show a high richness (i.e., a large number of different ASVs) but lower Shannon diversity, indicating a community with few high abundance and many very low abundance taxa.

**Figure 3.**
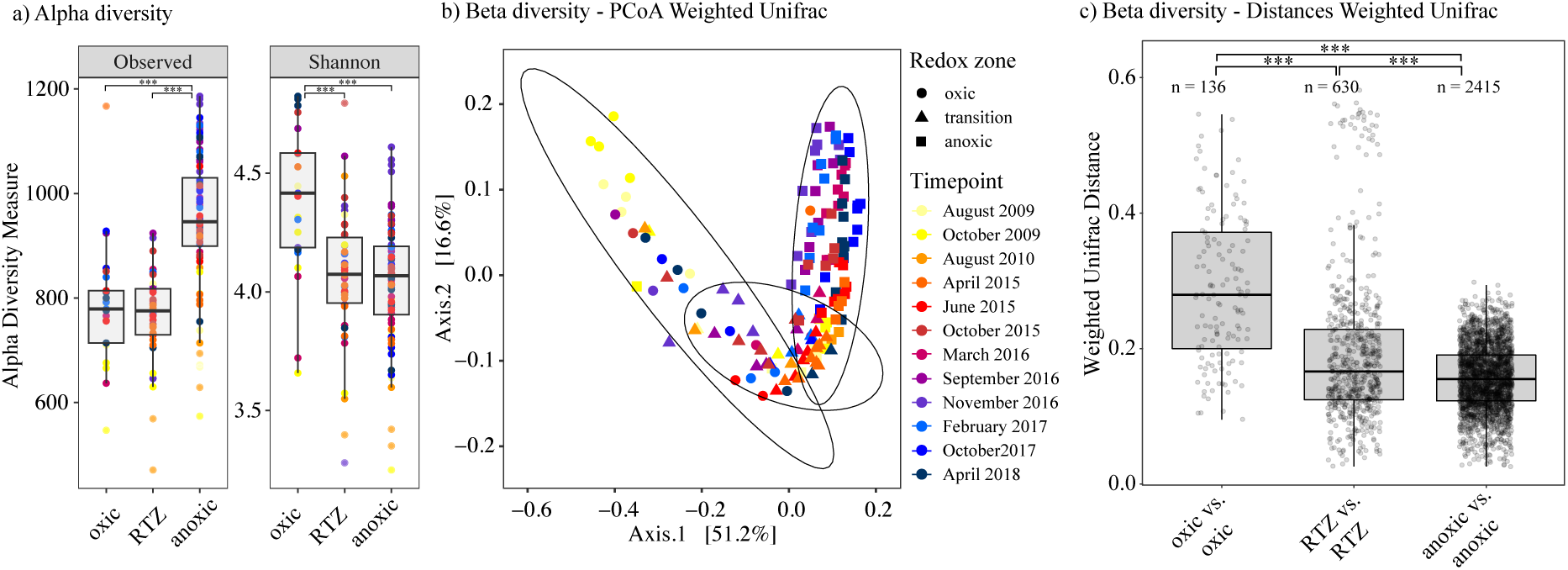
Characterization of the microbial communities in the water column of the Lake Lugano North Basin depending on redox regime (“oxic”, redox transition zone: “RTZ”, “anoxic”) and/or sampling timepoint using 16S rRNA gene amplicon sequencing variant (ASV) data from 2009 to 2018. **a)** Observed ASV richness and Shannon diversity, **b)** principal coordinate analysis (PCoA) of weighted UniFrac distances, and **c)** weighted UniFrac distances of microbial community structures within the three distinct redox zones. Stars indicate significance of pairwise t-tests (*p* < 0.001).

The PCoA showed that the microbial community structures followed the three redox zones, although the transitions were gradual and zones were overlapping (Fig. 3b). The community structures in the three defined redox zones were significantly different from each other (PERMANOVA; F_2,116_ = 114.3, R^2^ = 0.38, *p* < 0.001). Furthermore, significant differences were observed across timepoints (PERMANOVA; F_11,116_ = 13.3, R^2^ = 0.24, p < 0.001) and in the interaction between timepoints and redox zones (PERMANOVA; F_21,116_ = 5.5, R^2^ = 0.19, p < 0.001). The microbial community structure below the RTZ exhibited the least variability between different timepoints (Fig. 3b). This observation was confirmed by the pairwise Weighted UniFrac distances between samples within a specific redox zone, where the beta diversity below the RTZ was significantly lower than in the RTZ and in oxic-water samples (Fig. 3c; pairwise t-test; oxic vs. RTZ samples: t = 9.3, df = 196, *p* < 0.001; oxic vs. anoxic samples: t = 13.9, df = 138, *p* < 0.001; RTZ vs. anoxic samples: t = 8.0, df = 695, *p* < 0.001).

Phylogenetic analysis of 16S rRNA gene amplicons revealed Proteobacteria, Bacteroidota, Actinobacteriota, Chloroflexi, and Crenarchaeota as the most common phyla (Fig. S5a) and Actinobacteria, Bacteroidia, and Gammaproteobacteria as the most common classes (Fig S5b). Below the RTZ, the relative abundance of Desulfobacterota and Nanoarchaeota increased with depth in the water column. The composition of the microbial community (Fig. S6; phyla and classes) in the different redox zones over time showed that in the oxic water column, changes in relative abundance are more pronounced compared to the RTZ and the anoxic water column. Identified ASVs in the dataset associated to N- and S-cycling are summarized in Table S1 and Fig. S7-S11, whereas vertical distribution profiles of prominent representatives in the water column are shown in Fig 4. We detected the ammonia-oxidizing archaeon *Candidatus* Nitrosopumilus, the ammonia-oxidizing bacterium *Nitrosospira* and *Nitrosomonas*, and the nitrite-oxidizing bacterium *Nitrospira* with highest relative abundances at low O_2_ conditions above the RTZ (Fig. 4a-c). Furthermore, we detected the anammox-associated family Brocadiaceae (Fig. 4d). We also identified several taxa capable of N reduction, e.g., in the classes of Alphaproteobacteria, Gammaproteobacteria, and Campylobacteria (former Epsilonproteobacteria), which showed maximum relative abundances within the RTZ (examples in Fig. 4e-j). These taxa included organotrophic N reducers, such as *Denitratisoma, Sterolibacterium*, and *Dechloromonas*, S-oxidizing N reducers as *Sulfuritalea, Sulfuricurvum*, and *Sulfurimonas*, and NO_2_^-^-reducing, CH_4_-oxidizing *Ca.* Methylomirabilis. Some taxa showed two distinct maxima in their relative abundance, the first near the oxic-anoxic interface and another one further below within anoxic waters, e.g., *Sulfuritalea* and *Sulfurimonas* in March 2016. A diverse community of taxa capable to reduce SO_4_^2-^ and other S-compounds was present, mostly belonging to the Desulfobacterota (former Deltaproteobacteria), e.g., *Desulfobacca*, *Desulfocapsa* (Fig. 4k-l), and *Desulfovibrio.* In addition, we identified sulfate-reducing Firmicutes, including *Desulfosporosinu*s and *Desulfurispora*. Relative abundances of S-reducers generally increased with depth, but some genera exhibited distinct peaks in and/or below the RTZ, including *Desulfosporosinus* (Fig. S11a), *Desulfomonile* (Fig. S11g), and the versatile NO_3_^-^/Fe(III)/S^0^-reducing *Geobacter* (Fig. S11j).

**Figure 4.**
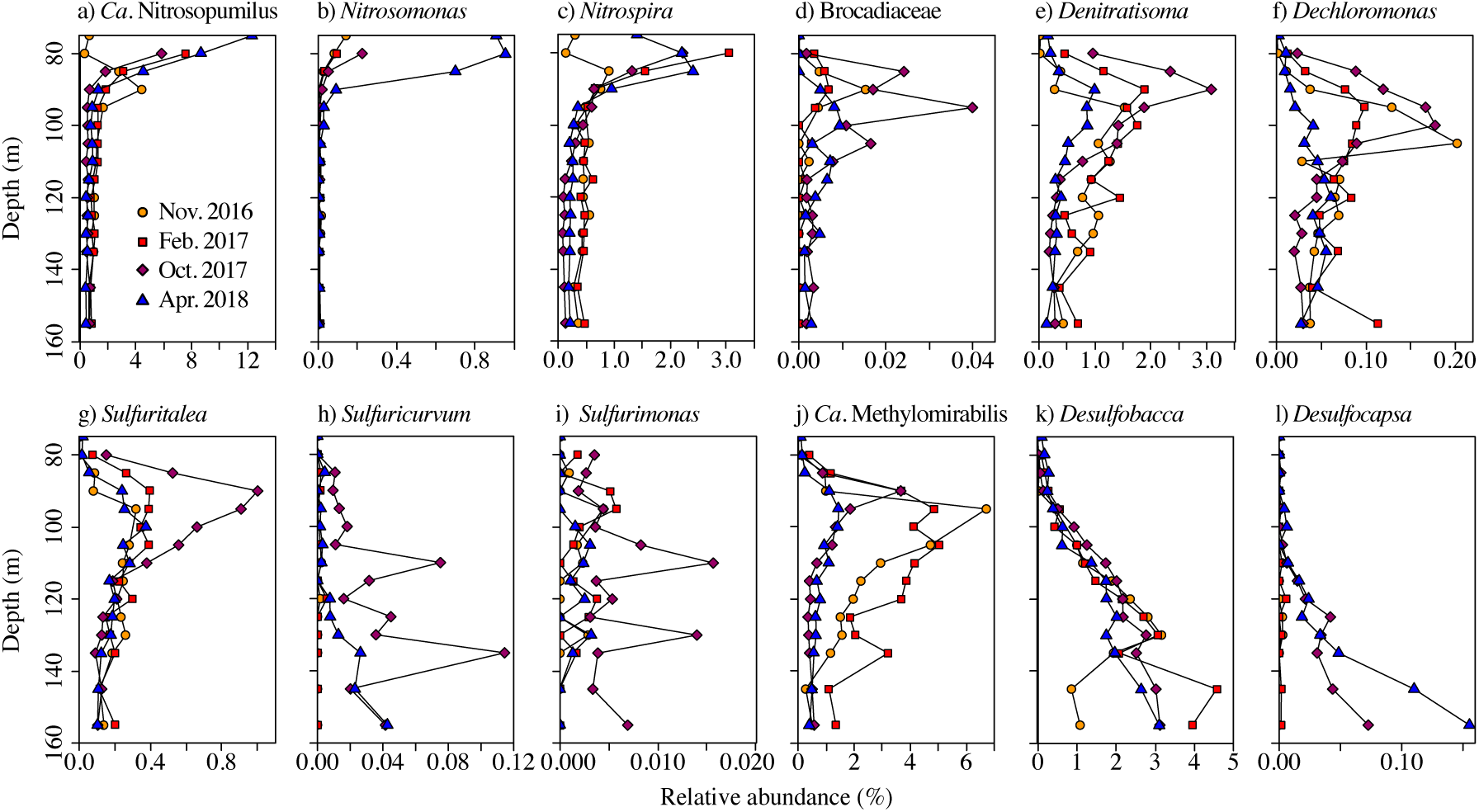
Depth distribution and temporal variability of the main taxa of N and S cycling microorganisms. Selected nitrifiers **(a-c)**, anammox-performing bacteria **(d)**, organotrophic-**(e, f)**, sulfidotrophic-**(g, h, i)** and methanotrophic N reducers **(j)**, and S-reducers **(k-l)** are presented. Data are based on relative abundances of 16S rRNA gene amplicons.

### Nitrogen transformation incubation experiments

In the unlabeled incubations with water from the RTZ (Fig. 5) with NO_3_^-^ added only, NO_3_^-^ consumption occurred, but was, in most cases, incomplete (reduction by ∼3-11 µM NO_3_^-^). Most of the NO_3_^-^ was only converted to NO_2_^-^ (Fig. 5d). Complete removal NO_3_^-^ (and subsequently also of NO_2_^-^) occurred only in one replicate experiment in February 2017 and one replicate experiment in April 2018 (Fig. 5A and D).

**Figure 5.**
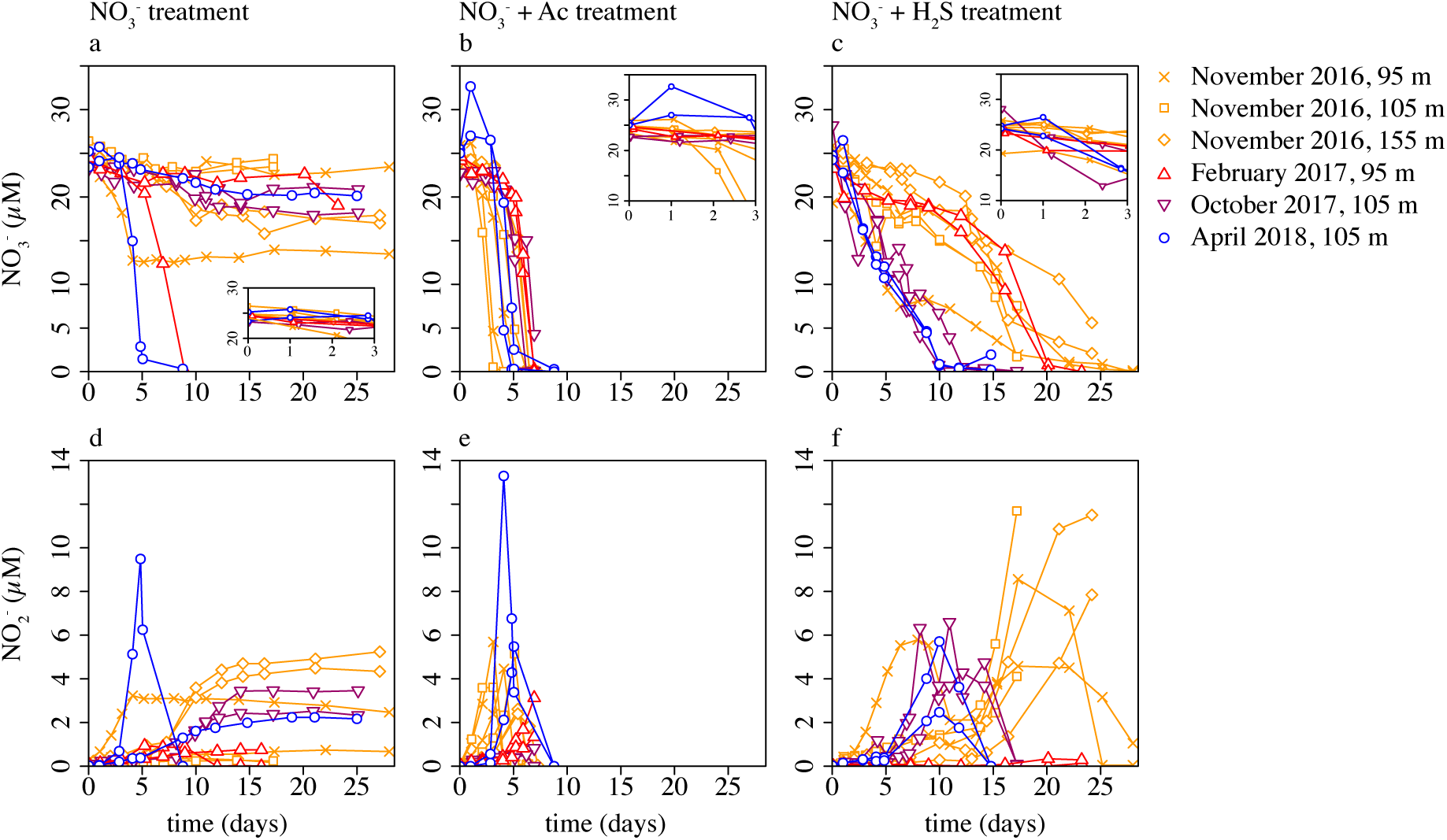
Concentrations of NO_3_^-^ **(a-c)** and NO_2_^-^ **(d-f**) in unlabeled incubation experiments performed with water samples from the redox transition zone (RTZ) in the North Basin of Lake Lugano, with added NO_3_^-^ **(a, d**), with NO_3_^-^ and acetate **(b, e)**, or with NO_3_^-^ and H_2_S **(c, f)**. Inserts show nitrate concentration changes within the first 3 days of the experiment.

The addition of acetate led to the relatively rapid and complete reduction of NO_3_^-^ within 4-5 days, after an initial lag phase (Fig. 5b). The addition of H_2_S caused complete NO_3_^-^ consumption within ∼10-30 days (Fig. 5c). The NO^3-^ drawdown was fastest, and almost linear, in water samples from the lower RTZ (105 m) in October 2017 and April 2018. NO_2_^-^ accumulation was transient in the incubations with acetate and H_2_S (Fig. 5e-f). The initial (i.e., first 2-3 days of incubation) NO_3_^-^ reduction rates (Fig. S12a) were highest for the NO_3_^-^ + H_2_S treatment with 1.4 ± 0.3 (standard error, SE) µmol NO_3_^-^ L^-1^ d^-1^. For the NO_3_^-^-only (0.6 ± 0.2 SE µmol NO_3_^-^ L^-1^ d^-1^) and the NO_3_^-^ + acetate treatment (0.9 ± 0.4 SE µmol NO_3_^-^ L^-1^ d^-1^), initial NO_3_^-^ reduction rates were lower. Whereas after the initial phase (Fig. S9b), NO_3_^-^ consumption rates in the NO_3_^-^-only treatment (0.7 ± 0.4 SE µmol NO_3_^-^ L^-1^ d^-1^) and the NO_3_^-^ + H_2_S treatment (1.7 ± 0.3 SE µmol NO_3_^-^ L^-1^ d^-1^) remained similar, NO_3_^-^ consumption rates in the NO_3_^-^ + acetate treatment increased to 3.2 ± 0.6 SE µmol NO_3_^-^ L^-1^ d^-1^ (ANOVA; F_1,22_ = 9.8, R^2^ = 0.31, *p* = 0.005). During the course of the incubations, H_2_S was consumed and S^0^ and S_2_O_3_^2-^ accumulated, but no substantial amounts of SO_4_^2-^ were produced (Table S3, Fig. S13).

### ^15^N incubation rate measurements

In the ^15^N-incubation experiments conducted in October 2017 and April 2018, denitrification rates (i.e., N_2_ production by denitrification) ranged between 28.8 and 113.0 nmol N-N_2_ L^-1^ d^-1^ in the treatments with ^15^NO_3_^-^ and no added electron donors (Table 2). Hence, N_2_ production rates were almost an order of magnitude lower than NO_3_^-^ consumption rates determined in parallel unlabeled experiments (see above), which can be explained by the accumulation of NO_2_^-^, which had also occurred in the unlabeled incubations (Fig. 5d). A representative example of a ^15^N-label incubation experiment from April 2018 is presented in Fig. S14. Amended acetate and H_2_S did not significantly change the measured denitrification rates (ANOVA; F_2,6_ = 2.1, R^2^ = 0.41, *p* = 0.21), which ranged between 19.4 and 31.7 nmol N-N_2_ L^-1^ d^-1^ for the ^15^NO_3_^-^ + acetate treatment and between 39.1 and 71.0 nmol N-N_2_ L^-1^ d^-1^ for the ^15^NO_3_^-^ + H_2_S treatment. Qualitatively consistent with the observed NO_3_^-^ reduction dynamics in the unlabeled incubations, acetate-supported ^30^N_2_ production started only after a lag phase of more than three days (Fig. S14e). In the ^15^NO_3_^-^ and ^15^NO_3_^-^ + acetate incubations, DNRA rates were mostly undetectable. Only in April 2018 (95 m, NO_3_^-^ treatment: 142.2 nmol ^15^N-NH_4_^+^ L^-1^ d^-1^ and 105 m, ^15^NO_3_^-^ + acetate treatment: 161.3 nmol ^15^N-NH_4_^+^ L^-1^ d^-1^), they were significant and contributed more than denitrification to the total NO_3_^-^ turnover. In contrast, the addition of H_2_S resulted in high DNRA rates between 471.1 and 1706.6 nmol ^15^N-NH_4_^+^ L^-1^ d^-1^, representing 87-97% of the total ^15^N-N_2_ + ^15^N-NH_4_^+^ produced. Anammox rates were generally low in all treatments, ranging from 0.1 up to 12.5 nmol ^15^N-N_2_ L^-1^ d^-1^. They accounted for 0 to 30% (but mostly <10%; Table 2) of the produced ^15^N-N_2_ + ^15^N-NH_4_^+^, without any significant differences between treatments (ANOVA; F_2,6_ = 1.5, R^2^ = 0.33, *p* = 0.30), and were not considered further in the discussion.

## Discussion

The results from unlabeled and ^15^N-label incubation experiments and 16S rRNA gene sequencing data provide conclusive evidence for active N-cycling around the RTZ, closely coupled to the S- and C-cycles. The overlapping concentration profiles of NH_4_^+^ and NO_3_^-^ with complete consumption of both substrates within the RTZ suggest that ammonia oxidation (aerobic and anaerobic) and nitrate reduction take place simultaneously, and in close vicinity. In the following sections, we discuss the relevance of H_2_S and organic compounds (including CH_4_) for N reduction, the N reduction pathway they fuel, as well as potential microbial players involved. We thereby elucidate the importance of closely coupled (and possibly cryptic) N- and S-cycling within the RTZ.

### Role of organic electron donors for N reduction

Limited availability of reactive organic material (OM) for canonical denitrification was indicated in the unlabeled incubation experiments, where, in most cases, only a relatively small fraction of the added NO_3_^-^ was reduced. In two incubations, NO_3_^-^ was consumed completely within about one week, but NO_3_^-^ reduction rates were low until two to three days into the experiment. In these bottles, we may have trapped sinking organic aggregates. In situ, they would have passed the RTZ in less than 2 days, based on sinking rates reported by Grossart and Simon (1998), but in the incubations, the organic substrate within the aggregates could be exploited more efficiently by the organotrophic denitrifying microorganisms present. The ^15^N-label incubation experiments confirm an overall relatively low denitrification activity. Even during the algal bloom in April 2018 (Fig. S2), when large amounts of OM were produced (46.5 g C m^-2^; www.cipais.org), ^15^NO_3_^-^ added to water samples did not result in higher denitrification (or DNRA) rates. Also, the addition of acetate (an important intermediate of central metabolisms and an important compound during OM degradation) as electron donor only enhanced NO_3_^-^ reduction after a lag phase, which was similarly observed by Wenk et al. (2013). This suggests that organotrophic denitrifiers are present within the Lake Lugano RTZ, but are not very active. They respond, however, to organic-substrate addition (e.g., acetate) with growth and, hence, increased N reduction.

We hypothesized that in spring/summer, during periods of higher OM production and export, organotrophic denitrification rates are higher compared to other seasons. However, this was not evident in our data set. Readily degradable OM, produced in the primary production zone at ∼5-25 m depth, as indicated by maximum O_2_ and Chl *a* concentrations and low-nitrate conditions during algae blooms in April (Fig. S1, Fig. S2), is likely scarce in the RTZ for several reasons. A large portion of the freshly produced OM is already consumed in subsurface waters through aerobic respiration before even reaching the RTZ at 80-110 m depth. This is indicated by a strong decline of dissolved O_2_ in and below the primary production zone in summer and autumn (Fig. S1a). As a consequence, only a small fraction of the exported OM, with a less accessible (i.e., less susceptible to hydrolysis and microbial degradation), organic compound composition, makes it to the RTZ in the relatively deep northern basin of Lake Lugano, also during periods of high primary productivity. Moreover, the residence time of sinking detrital OM in the RTZ might be too short for microorganisms to exploit the OM (Grossart and Simon 1998). Indeed, an unrestrained transit of particles through the RTZ is likely because the RTZ in the Lake Lugano North Basin is not associated with a notable density gradient and has a weak water column stability (Fig. S1e), thus preventing any significant retention of sinking OM aggregates (Alldredge and Cracker 1995). In summary, also during the productive season, availability of C_org_ in the RTZ is likely low, limiting heterotrophic denitrification.

We note that our ^15^NO_3_^-^ addition experiments without any additional electron donors, while consistent with previous reports by Wenk et al. (2014), might slightly underestimate the in situ rates of denitrification, because volatile electron donors such as CH_4_ and H_2_S were likely removed during the purging. Nevertheless, denitrification rates in the Lake Lugano North Basin (up to 113 nmol N-N_2_ L^-1^ d^-1^) are at least one order of magnitude lower compared to several other lakes (Table 1). For instance, maximum denitrification rates reached >8000 nmol N-N_2_ L^-1^ d^-1^ in tropical Lake Kivu (Roland et al. 2018), up to ∼4800 nmol N-N_2_ L^-1^ d^-1^ in Lake Tanganyika (Schubert et al. 2006), and up to 1000 nmol N-N_2_ L^-1^ d^-1^ in Wintergreen Lake (Burgin et al. 2012). The higher rates observed in other lakes could be due to increased temperature (e.g., Lake Kivu) and/or a shorter distance between the primary production zone and the RTZ, where less OM is degraded before reaching the active denitrification zone. For example, in Lake Kivu the RTZ lies between 40 and 50 m (Roland et al. 2018) and in Wintergreen Lake at 4-5 m depth (Burgin et al. 2012). Moreover, in relatively shallow RTZs, light penetration can enable OM production in situ, and sustain organotrophic denitrification, as observed in Lake La Cruz. (Oswald et al. 2016; Tischer et al. 2022). Lake Tanganyika has a similar O_2_-depletion depth as Lake Lugano of 80-90 m (Schubert et al. 2006) and similar mean annual daily rates of primary production of 1 g C m^-2^ d^-1^ (Hecky and Fee 1981). Yet, also in Lake Tanganyika, there is evidence for a deep secondary, anoxic Chl *a* maximum suggesting phototrophic growth near the RTZ, at least in the northern part of the basin, which may partly support denitrification (and anammox) (Ehrenfels et al. 2023). Moreover, besides the higher Chl *a* content at greater depths in Lake Tanganyika, and higher water column temperatures, other factors may be particularly conducive to higher rates of heterotrophic denitrification. More specifically, a relatively deep thermocline at 70 m depth (Mziray et al. 2018) likely subserves an increased OM availability at the RTZ by promoting the retention of OM aggregates along the density gradient close to the RTZ.

### Sulfur-dependent N reduction

We demonstrate a high potential of S-dependent DNRA in the ^15^N-label experiments with up to 1707 nmol NH_4_^+^ L^-1^ d^-1^. Active S-dependent N reduction is also indicated by the presence of intermediates of S-oxidation, especially of S^0^ and S_2_O_3_^2-^, which accumulated in the unlabeled incubation experiments and were present in situ with up to ∼0.25 µM S^0^ and ∼0.10 µM S_2_O_3_^2-^. Contrary to S-dependent DNRA, S-dependent denitrification showed low potential to contribute to nitrate reduction, as in situ denitrification rates were generally low (up to 71 nmol N-N_2_ L^-1^ d^-1^), and were not stimulated in the incubations with added ^15^NO_3_^-^ plus H_2_S, compared to the ^15^NO_3_^-^-only treatments.

Measured rates in the water column of other lakes revealed significantly different relative and absolute importance of DNRA and denitrification coupled to S-oxidation and OM-oxidation, respectively. For example, in Lake Kivu (Roland et al. 2018) and in Wintergreen Lake (Burgin et al. 2012), both organotrophic and S-dependent denitrification and DNRA (partly coupled to S-oxidation) have been detected. Yet, in both cases, DNRA accounted for only up to ∼15% of the total ^15^N-turnover, while in Lake Lugano DNRA potentially contributed up to 97%. The OM limitation in the RTZ in Lake Lugano may competitively favor the activity of S-dependent N reduction. However, ambient sulfide concentrations were below the detection limit within the RTZ, and high DNRA rates were only observed upon H_2_S addition. Hence, the dominance of S-dependent DNRA in the water column under in situ conditions remains to be confirmed, for instance, by quantifying the in situ expression of the *nrfA* gene, a molecular marker for DNRA (Pandey et al. 2020).

The 16S rRNA gene sequencing analysis revealed a number of bacterial genera in the RTZ, which likely perform the observed S-dependent N reduction, like *Sulfuritalea*, with a relatively high abundance of up to 1%, as well as less abundant taxa such as *Sulfuricurvum* and *Sulfurimonas*. We speculate that the N-reducing *Dechloromonas*, which also possesses genes for the Sox sulfur-oxidation enzyme system (Luo et al. 2018), and proliferated in the unlabeled incubations with added NO_3_^-^ and H_2_S (Tischer et al. 2024/in prep.), may also be an important S-oxidizer in the RTZ, in addition to performing organotrophic denitrification.

Based on 16S rRNA gene sequencing data, the observed sulfur speciation in the water column, and the H_2_S-addition experiments, we suggest that the most active zone for S-oxidizing N reducers may be a “cryptic-sulfur-cycling” zone, where S-oxidation and reduction are so tightly coupled that free H_2_S is undetectable. Cryptic sulfur cycling has already been reported for marine oxygen minimum zones, albeit with much lower rates (Canfield et al. 2010; Callbeck et al. 2018). In the North Basin of Lake Lugano, in the 10 to 40 m thick anoxic water layer between the water depths where O_2_ disappears and where H_2_S begins to accumulate, respectively, we detected S^0^, a high-rate potential for S-dependent DNRA, and maximum relative abundances of *Sulfuritalea*. We hence argue that H_2_S diffusing towards the RTZ is quantitatively oxidized by nitrate-reducing microbes performing DNRA. The volumetric sulfide oxidation rate that would be required to explain the observed flux of H_2_S toward the RTZ (85 to 304 µmol d^-1^ m^-2^) was, on average, 0.013 µmol H_2_S L^-1^ d^-1^ (assuming a 15 m thick layer of S-dependent denitrification activity; Table S2). The H_2_S consumption in the unlabeled anoxic incubation experiments, however, shows, with an average of 4.6 µmol H_2_S L^-1^ d^-1^, a more than 350-times higher potential of H_2_S utilization than what would be required to completely oxidize the upward diffusing H_2_S in the water column. Furthermore, the presence of a community of S-reducing taxa, dominated by *Desulfobacca* (Fig. 4k), within, and below the RTZ, underscores the high potential for producing reduced S species in the same water-column region where the high S oxidation potential was observed. Hence, even in the non-sulfidic anoxic water column, microbial sulfate reduction represents a continuous source of sulfide, which is immediately consumed by S-oxidizing microorganisms. This process thus supports cryptic S-dependent DNRA, and to a lesser extent, S-dependent denitrification.

### Scope for methane-dependent N reduction

CH_4_ may serve as a viable alternative electron donor for microbial N reduction, indicated by its depletion within the RTZ and the elevated abundance and activity of *Ca.* Methylomirabilis in link with CH_4_ oxidation stimulation by NO_2_^-^ (Su et al. 2023). Methane and nitrate concentration profiles overlap in a ∼10 m-thick water layer (Fig. 1) where peaking relative abundances of *Ca.* Methylomirabilis were observed, underscoring that the basic hydro-biogeochemical and microbiological requirements for nitrite/nitrate reduction with methane are fulfilled. In addition to anaerobic N reduction by *Ca.* Methylomirabilis, micro-aerobic CH_4_-oxidation coupled to N reduction might occur, potentially by *Crenothrix*, which is abundant in the upper RTZ, as discussed in Su et al. (2023).

We did not perform CH_4_ addition experiments to determine the potential for CH_4_-dependent nitrate/nitrite reduction in the water column. Nevertheless, some indication for such a microbial pathway is provided by the methane oxidation rates reported by Su et al. (2023), which were ∼0.05-0.06 µmol L^-1^ d^-1^, translating to a maximum CH_4_-dependent NO_2_^-^ reduction rate of ∼0.08 µmol NO_2_^-^ L^-1^ d^-1^ (assuming a stochiometric ratio of 3:5 for CH_4_:NO_2_^-^ consumption; Ettwig et al. 2010). This rate would be in the same range as some of the denitrification rates determined in the presence of H_2_S. However, it must be considered as an upper-limit estimate because it assumes that all CH_4_ is oxidized with NO_3_^-^ as electron acceptor, and even this maximum possible rate is clearly lower than the observed DNRA rates determined under the same experimental conditions (Table 2). Moreover, parallel investigations by our group assessing the role of CH_4_ in N_2_O formation during denitrification in the same lake basin (T. Einzmann, unpubl.), showed that reductive N_2_O production is not stimulated by CH_4_, suggesting, indeed, a subordinate role of CH_4_ as electron donor for denitrification.

### Cryptic N cycling through nitrification and DNRA coupling

We suggest that N recycling via S-dependent DNRA and nitrification is a key mechanism in the RTZ of the Lake Lugano North Basin, which helps to retain large amounts of bioavailable N, while N_2_ production by denitrification and anammox removes only a minor portion of the fixed N. High activity of ammonia and nitrite oxidation close to the oxic-anoxic interface is evidenced by the presence of a diverse and abundant community of ammonia and nitrite-oxidizers, including *Ca.* Nitrosopumilus. This archaeal ammonia oxidizer is common in marine environments (Könneke et al. 2005), but has also been observed more recently in deep freshwater bodies such as Lake Constance and Lake Maggiore (Coci et al. 2015; Klotz et al. 2022). Although nitrification rates remain to be quantified, maximum relative abundances close to the upper boundary of the RTZ indicate that *Nitrosomonas* or *Nitrosospira* encounter excellent conditions for nitrification, with replete O_2_ and a constant supply of NH_4_^+^. *Nitrospira* might perform complete ammonia oxidation (comammox) from NH_4_^+^ to NO_3_^-^ (Sakuola et al. 2021), and both *Nitrosospira* and *Nitrosomonas* can perform nitrifier denitrification, where NO_2_^-^ produced from NH_3_ is subsequently reduced to NO and N_2_O under low O_2_ conditions (Wrage et al. 2004; Shaw et al. 2006). Studies on other stratified lakes have shown evidence of active nitrification occurring in the respective RTZs (Christofi et al. 1981; Pajares et al. 2017). More specifically, in the RTZ of a monomictic tropical lake, Pajares et al. (2017) demonstrated the co-occurrence of nitrification, DNRA, and denitrification genes, highlighting the importance of internal N recycling in the lake.

Nitrification in the RTZ of the Lake Lugano North Basin is likely more important than previously reported by Wenk et al. (2013), who interpreted the non-overlapping profiles of O_2_ and NH_4_^+^ (i.e., the quantitative consumption of NH_4_^+^ below the redoxcline) as evidence that ammonium oxidation primarily occurs anaerobically, driven by anammox bacteria. However, overlapping O_2_ and NH_4_^+^ concentration profiles for 2015 to 2018 (this study), recent results from dual nitrate N and O isotope measurements (Tischer et al. 2024/in prep.), and an observed community shift of certain nitrifying taxa like *Nitrosospira* and Nitrosomonadaceae (Table S1), point towards an increased role of nitrification in the RTZ of Lake Lugano’s North Basin over the past decade.

We also argue that, independent of the relatively low turbulent-diffusive fluxes of N compounds towards to RTZ (Table S2), the overall N turnover may be quite high, yet cryptic, due to efficient recycling between the ammonium and nitrate pools via DNRA and nitrification. Indeed, an average net consumption rate of not more than 0.055 µmol NO_3_^-^ L^-1^ d^-1^ would have been necessary to explain the low observed nitrate flux towards the RTZ. In the NO_3_^-^-only treatment of the unlabeled incubations, however, potential NO_3_^-^ reduction rates were much higher (average 0.7 µmol NO_3_^-^ L^-1^ d^-1^), suggesting that the flux-based rate estimates underestimate the potential for NO_3_^-^ reduction and that nitrate reduction is closely coupled to microaerobic nitrate regeneration. The latter, in turn, may be supported by DNRA, especially by S-dependent DNRA, which showed a high potential in the ^15^N-label incubations. Cryptic N cycling, i.e., without measurable intermediates as nitrite or ammonium, has been demonstrated in oxic riverbeds (Ooyang et al. 2021). In addition, Lam et al. (2009) demonstrated that a substantial fraction of anammox in the Peruvian oxygen minimum zone is supported by DNRA, without significant accumulation of NH_4_^+^. In the Lake Lugano North Basin of today, anammox arguably plays a subordinate role. Here, it is a close coupling between DNRA, nitrification, and denitrification that efficiently turns over fixed N and ultimately eliminates parts of it as N_2_. As with the “cryptic S cycle” mentioned above, we propose a closely linked “cryptic N turnover” involving S-dependent denitrification and DNRA in the RTZ water layer, where NO_3_^-^ and H_2_S are not detected and NO_2_^-^ usually does not accumulate.

### Electron donors from sediment fuel water column N cycle

We argue that N reduction (denitrification and DNRA) in the North Basin of Lake Lugano is mainly driven by the constant supply of H_2_S and CH_4_, because readily accessible OM for organotrophic denitrification is scarce in the RTZ, as conceptualized in Figure 6. As mentioned before, most of the OM from primary production is either already degraded in the upper oxic water column, or is not readily available as electron donor in the RTZ because sinking aggregates are not retained, and/or organic particles reaching the deeper hypolimnion are more refractory/stable (i.e., less prone to microbial attack). Instead, OM will accumulate in the permanently anoxic lake sediments, where it is further degraded through fermentation, SO_4_^2-^ reduction, and methanogenesis, effectively transferring electrons to the benthic H_2_S and CH_4_ pools (Blees et al. 2014). Microorganisms in the sediment are more abundant and the microbial community is more diverse compared to the water column (Bartosiewicz et al. 2024). The microorganisms have essentially unlimited time to adapt their metabolism to, and exploit, specific compounds. In this context, the sediments and their natural microbial community may be seen as “electron donor refinery”, which turn a non-steady flow of less accessible electron donor compounds from the water column into a sustained production of H_2_S and CH_4_. These metabolites then diffuse out of the sediments towards the RTZ, where they serve as continuous and mostly season-independent electron donors for chemolithotrophic and CH_4_-dependent N reduction, respectively. The sustained benthic flux of electron-rich substrates thus allows for the establishment of a stable assemblage of N- and S-cycling microorganisms structured along a spatial sequence from oxic, to low O_2_, to anoxic conditions, including key organisms such as nitrifying *Ca.* Nitrosopumilus, *Nitrosomonas*, and *Nitrospira* and N reducing *Denitratisoma*, *Sulfuritalea*, and *Ca.* Methylomirabilis. There was little variation within these guilds of microorganisms for the duration of our study, probably due to the stable environmental conditions and reduced or absent grazing pressure in the lower part of the RTZ and the anoxic water layer. The concept of low, yet very constant, supply of H_2_S and CH_4_ is supported by the low weighted UniFrac distances within the anoxic zone (Fig. 3c), indicating that the microbial community is more stable compared to the oxic water column, where grazing by e.g., protists and fluctuations in OM supply lead to a more variable community composition. We propose that this ultimately explains the dominance of S-, and possibly CH_4_, driven NO_3_^-^/NO_2_^-^ reduction over canonical organotrophic denitrification within the RTZ of the Lake Lugano North Basin.

**Figure 6.**
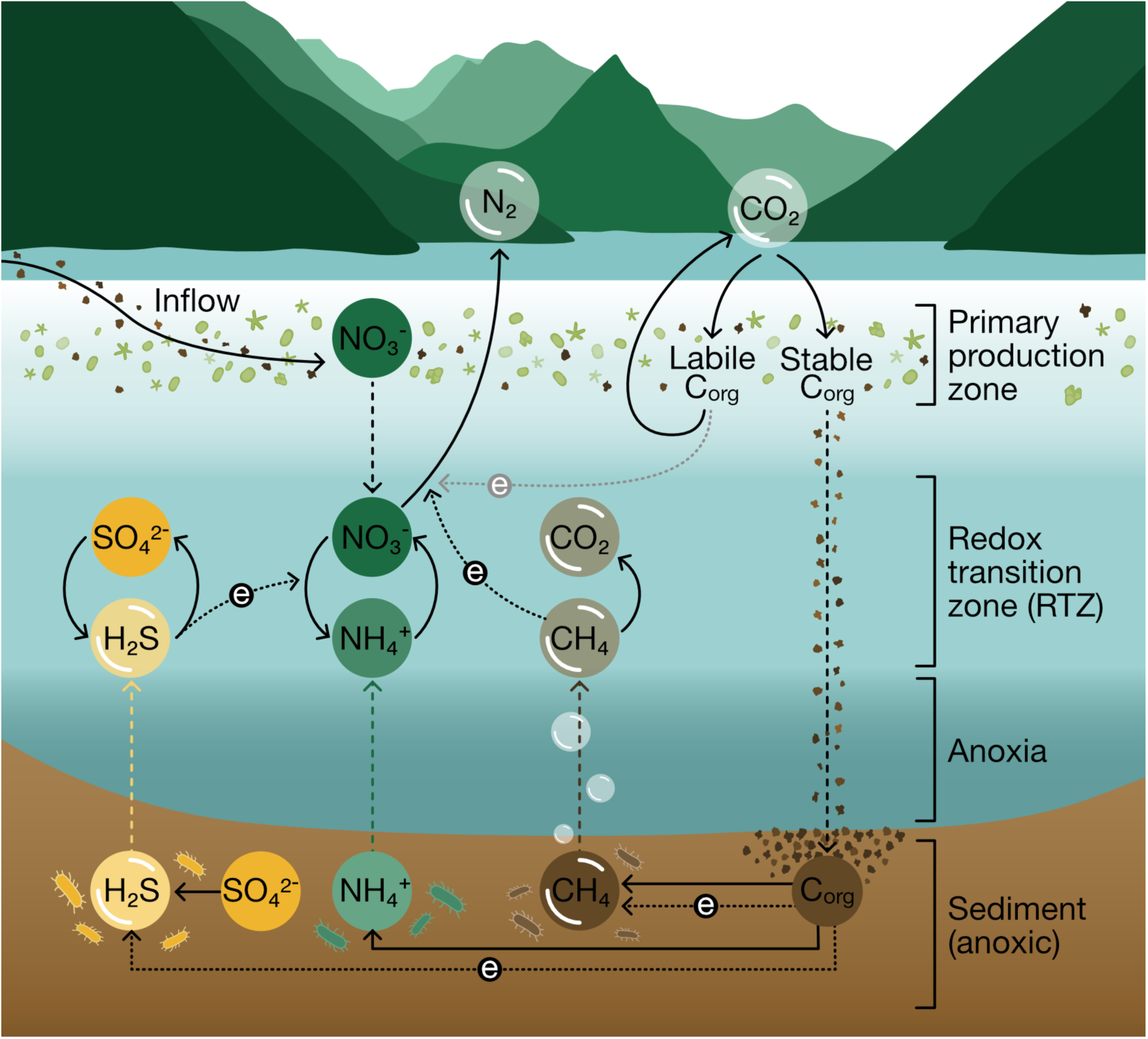
Schematic illustration of N formation processes in the Lake Lugano North Basin coupled to the S cycle and the C-cycle. Solid lines indicate transformation processes, dashed lines indicate transport processes or symbolize the flow of electrons. Easily accessible fractions of fresh organic carbon (labile C_org_) produced in the primary production zone are mostly degraded in the oxic water column before reaching the redox transition zone (RTZ), while the less accessible or more stable fractions (stable C_org_) are incompletely oxidized and sink to the ground where they are degraded under anoxic conditions through fermentation, S-reduction and methanogenesis. Sulfide and methane, along with NH_4_^+^ from organic N mineralization, diffuse to the RTZ, where they are used steadily as electron donors for denitrification and dissimilatory nitrate reduction to ammonium (DNRA). This way, the sediments serve as “electron donor refinery” that generates a steady flux of substrates for denitrification. Within the RTZ, N-cycling via nitrification and DNRA and S-cycling via S-oxidation and S-reduction likely takes place.

## Conclusions and implications

We investigated cycling and elimination of fixed N, coupled to the S- and C-cycles, in the RTZ of the Lake Lugano North Basin. Incubation experiments, rate measurements, and 16S rRNA gene sequencing data uncovered a complex interaction of multiple oxidative and reductive N transformation processes, supported by a stable community of relatively few but abundant key microorganisms. We provide evidence that the meromictic North Basin of Lake Lugano, with its deep oxycline, creates conditions that favor sulfur- and methane-driven N reduction over canonical organotrophic denitrification. In particular, H_2_S appears to play an important role in DNRA within the RTZ. The factors that determine the relative importance of denitrification and DNRA under environmentally relevant substrate concentrations can vary and are not fully understood (Pandey et al. 2020). While in other environments (e.g., lacustrine sediments), dissolved H_2_S has been shown to stimulate denitrification but not DNRA (Cojean et al. 2020), we demonstrate here that H_2_S can strongly stimulate DNRA in a water column with limited OM availability.

Nitrification, denitrification, DNRA, and reduction of S species take place in basically the same water mass around the oxic-anoxic interface, supporting an interactive, and to large parts “cryptic”, S- and N-cycling. This is the result of a diverse microbial community of S- and N transforming taxa, which can benefit from mutualistic interactions, e.g., through the continuous exchange of reduced and oxidized compounds. The close coupling and partitioning of the different NO_3_^-^-transforming metabolisms will determine whether fixed-N is recycled and remains in the system or is eliminated from it. Because DNRA (and nitrification), in contrast to denitrification, results in the retention of reactive N in the environment, our findings have important implications for the N budget of the Lake Lugano North Basin. More than 50 years ago, high inputs of P and N have shifted the lake into a meromictic and meso-eutrophic state. Despite a substantial reduction of external P inputs over the last 40 years (Barbieri and Mosello 1992), the trophic state of the lake has only weakly improved (Lepori et al. 2018). The relatively low fixed-N removal potential in the North Basin, coupled with efficient internal N cycling via DNRA, might contribute to maintaining elevated nitrate levels in the water column, thereby slowing down the restoration process of the lake.

## Data availability

Raw sequence data are made available at NCBI under the BioProjectID PRJNA672280 with the accession numbers SRR12936362 through SRR12936382; MH111698 through MH113143. Water column chemistry and experimental data are available on the Open Science Framework at https://osf.io/gfx7w/.

## Acknowledgements

We thank Marco Simona, Stefano Beatrizotti, Adeline Cojean-Egger, Maciej Bartosiewicz, and Lea Steinle for their help during the sampling campaigns on Lake Lugano. Long-term oxygen, nitrate, and primary productivity data were generated within a research program funded by the Dipartimento del Territorio (Department of Environment) of the Canton of Ticino, Switzerland, and the International Commission for the Protection of Italian-Swiss Waters (CIPAIS). We are also grateful to Thomas Kuhn and Judith Kobler for providing technical support in the laboratory. We also thank Teresa Einzmann for providing unpublished experimental data from Lake Lugano. The research was funded by the Swiss National Science Foundation project 153055 granted to JZ and MFL.

## Author contribution statement

JZ and MFL conceived the research project. JT, JZ, and MFL conceptualized research and experimental design. JT, JZ, MFL, GS and FL conducted sampling campaigns and contributed to the data acquisition of physicochemical parameters in the water column of Lake Lugano. The lake water incubation experiments were conducted by JT and analyzed by JT and JZ. JT, JZ, and GS carried out the data analysis of 16S rRNA sequencing data. JT wrote the paper, with substantial input from JZ, MFL, GS, and FL. JT, JZ, and MFL are accountable for the integrity of the data, analysis, and presentation of the findings.

## References

1. Alldredge, A. L., and K. M. Cracker. 1995. Why do sinking mucilage aggregates accumulate in the water column? Sci Total Environ 165: 15–22. doi:10.1016/0048-9697(95)04539-D

2. Barbieri, A., and R. Mosello. 1992. Chemistry and trophic evolution of Lake Lugano in relation to nutrient budget. Aquat Sci 54: 219–237. doi:10.1007/BF00878138

3. Bartosiewicz, M., A. Przytulska, A. Birkholz, J. Zopfi, and M. F. Lehmann. 2024. Controls and significance of priming effects in lake sediments. Glob Chang Biol 30. doi:10.1111/GCB.17076

4. Blees, J. and others. 2014. Micro-aerobic bacterial methane oxidation in the chemocline and anoxic water column of deep south-Alpine Lake Lugano (Switzerland). Limnol Oceanogr 59: 311–324. doi:10.4319/lo.2014.59.2.0311

5. Braman, R. S., and S. A. Hendrix. 1989. Nanogram nitrite and nitrate determination in environmental and biological materials by vanadium(III) reduction with chemiluminescence detection. Anal Chem 61: 2715–2718. doi:10.1021/ac00199a007

6. Brunet, R., and L. Garcia-Gil. 1996. Sulfide-induced dissimilatory nitrate reduction to ammonia in anaerobic freshwater sediments. FEMS Microbiol Ecol 21: 131–138. doi:10.1016/0168-6496(96)00051-7

7. Burgin, A., S. Hamilton, S. Jones, and J. Lennon. 2012. Denitrification by sulfur-oxidizing bacteria in a eutrophic lake. Aquatic Microbial Ecology 66: 283–293. doi:10.3354/ame01574

8. Burgin, A. J., and S. K. Hamilton. 2007. Have we overemphasized the role of dentitrification in aquatic ecosystems? A review of nitrate removal pathways. Front Ecol Environ 5: 89–96. doi:10.1890/1540-9295(2007)5[89:HWOTRO]2.0.CO;2

9. Callbeck, C. M. and others. 2018. Oxygen minimum zone cryptic sulfur cycling sustained by offshore transport of key sulfur oxidizing oxidizing bacteria. Nat Commun 9: 1–11. doi:10.1038/s41467-018-04041-x

10. Canfield, D. E., F. J. Stewart, B. Thamdrup, L. De Brabandere, T. Dalsgaard, E. F. Delong, N. P. Revsbech, and O. Ulloa. 2010. A cryptic sulfur cycle in oxygen-minimum-zone waters off the Chilean coast. Science 330: 1375–1378. doi:10.1126/science.1196889

11. Christofi, N., T. Preston, and W. D. P. Stewart. 1981. Endogenous nitrate production in an experimental enclosure during summer stratification. Water Res 15: 343–349. doi:10.1016/0043-1354(81)90039-7

12. Cline, J. D. 1969. Spectrophotometric determination of hydrogen sulfide in natural waters. Limnol Oceanogr 14: 454–458. doi:10.4319/lo.1969.14.3.0454

13. Coci, M., N. Odermatt, M. M. Salcher, J. Pernthaler, and G. Corno. 2015. Ecology and distribution of Thaumarchaea in the deep hypolimnion of Lake Maggiore. Archaea 2015. doi:10.1155/2015/590434

14. Cojean, A. N. Y., M. F. Lehmann, E. K. Robertson, B. Thamdrup, and J. Zopfi. 2020. Controls of H_2_S, Fe^2+^, and Mn^2+^ on Microbial NO_3_^-^-reducing processes in sediments of an eutrophic lake. Front Microbiol 11: 1–17. doi:10.3389/fmicb.2020.01158

15. Daims, H., S. Lücker, and M. Wagner. 2016. A new perspective on microbes formerly known as nitrite-oxidizing bacteria. Trends Microbiol 24: 699–712. doi:10.1016/j.tim.2016.05.004.A

16. Dong, L. F., M. N. Sobey, C. J. Smith, I. Rusmana, W. Phillips, A. Stott, A. M. Osborn, and D. B. Nedwell. 2011. Dissimilatory reduction of nitrate to ammonium, not denitrification or anammox, dominates benthic nitrate reduction in tropical estuaries. Limnol Oceanogr 56: 279–291. doi:10.4319/lo.2011.56.1.0279

17. Edgar, R. C. 2010. Search and clustering orders of magnitude faster than BLAST. 26: 2460–2461. doi:10.1093/bioinformatics/btq461

18. Edgar, R. C. 2013. UPARSE : highly accurate OTU sequences from microbial amplicon reads. 10. doi:10.1038/nmeth.2604

19. Edgar, R. C. 2016. SINTAX: a simple non-Bayesian taxonomy classifier for 16S and ITS sequences. Ehrenfels, B. and others. 2023. Hydrodynamic regimes modulate nitrogen fixation and the mode of diazotrophy in Lake Tanganyika. Nature Communications 2023 14:1 14: 1–13. doi:10.1038/s41467-023-42391-3

20. Ettwig, K. F. and others. 2010. Nitrite-driven anaerobic methane oxidation by oxygenic bacteria. Nature 464: 543–548. doi:10.1038/nature08883

21. Fahey, R. C., and G. L. Newton. 1987. Determination of low-molecular-weight thiols using monobromobimane fluorescent labeling and high-performance liquid chromatography. Methods Enzymol 143: 85–96. doi:10.1016/0076-6879(87)43016-4

22. Grossart, H.-P., and M. Simon. 1998. Significance of limnetic organic aggregates (lake snow) for the sinking flux of particulate organic matter in a large lake. Aquatic Microbial Ecology 15: 115–125. doi:10.3354/ame015115

23. Gruber, N., and J. N. Galloway. 2008. An Earth-system perspective of the global nitrogen cycle. Nature 451: 293–296. doi:10.1038/nature06592

24. Hansen, H. P., and F. Koroleff. 1999. Determination of nutrients, p. 159–228. In K. Grasshoff, K. Kremling, and M. Ehrhardt [eds.], Methods of Seawater Analysis. Wiley.

25. Hecky, R. E., and E. J. Fee. 1981. Primary production and rates of algal growth in Lake Tanganyika. Limnol Oceanogr 26: 532–547. doi:10.4319/lo.1981.26.3.0532

26. Holzner, C. P., W. Aeschbach-Hertig, M. Simona, M. Veronesi, D. M. Imboden, and R. Kipfer. 2009. Exceptional mixing events in meromictic Lake Lugano (Switzerland/Italy), studied using environmental tracers. Limnol Oceanogr 54: 1113–1124. doi:10.4319/lo.2009.54.4.1113

27. Howarth, R. W. and others. 1996. Regional nitrogen budgets and riverine N & P fluxes for the drainages to the North Atlantic Ocean: Natural and human influences. Biogeochemistry 35: 75–139. doi:10.1007/BF02179825

28. Hulth, S., R. C. Aller, D. E. Canfield, T. Dalsgaard, P. Engström, F. Gilbert, K. Sundbäck, and B. Thamdrup. 2005. Nitrogen removal in marine environments: Recent findings and future research challenges. Mar Chem 94: 125–145. doi:10.1016/j.marchem.2004.07.013

29. Jetten, M. S. M., L. Van Niftrik, M. Strous, B. Kartal, J. T. Keltjens, and H. J. M. Op den Camp. 2009. Biochemistry and molecular biology of anammox bacteria. Crit Rev Biochem Mol Biol 44: 65–84. doi:10.1080/10409230902722783

30. Kelso, B. H. L., R. V Smith, R. J. Laughlin, and S. D. Lennox. 1997. Dissimilatory nitrate reduction in anaerobic sediments leading to river nitrite accumulation. 63: 4679–4685. doi:10.1128/aem.63.12.4679-4685.1997

31. Kits, K. D., M. G. Klotz, and L. Y. Stein. 2015. Methane oxidation coupled to nitrate reduction under hypoxia by the Gammaproteobacterium *Methylomonas denitrificans*, sp. nov. type strain FJG1. Environ Microbiol 17: 3219–3232. doi:10.1111/1462-2920.12772

32. Klotz, F., K. Kitzinger, D. K. Ngugi, P. Büsing, S. Littmann, M. M. M. Kuypers, and M. Pester. 2022. Quantification of archaea-driven freshwater nitrification from single cell to ecosystem levels. ISME Journal 16: 1647–1656. doi:10.1038/s41396-022-01216-9

33. Kojima, H., and M. Fukui. 2011. *Sulfuritalea hydrogenivorans* gen. nov., sp. nov., a facultative autotroph isolated from a freshwater lake. Int J Syst Evol Microbiol 61: 1651–1655. doi:10.1099/ijs.0.024968-0

34. Könneke, M., A. E. Bernhard, J. R. de la Torre, C. B. Walker, J. B. Waterbury, and D. A. Stahl. 2005. Isolation of an autotrophic ammonia-oxidizing marine archaeon. Nature 437: 543–546. doi:10.1038/nature03911

35. Kowalchuk, G. A., and J. R. Stephen. 2001. Ammonia-oxidizing bacteria: A model for molecular microbial ecology. Annu Rev Microbiol 55: 485–529. doi:10.1146/annurev.micro.55.1.485

36. Lehmann, M. F., S. M. Bernasconi, J. A. McKenzie, A. Barbieri, M. Simona, and M. Veronesi. 2004. Seasonal variation of the δ^13^C and δ^15^N of particulate and dissolved carbon and nitrogen in Lake Lugano: Constraints on biogeochemical cycling in a eutrophic lake. Limnol Oceanogr 49: 415–429. doi:10.4319/lo.2004.49.2.0415

37. Lepori, F., M. Bartosiewicz, M. Simona, and M. Veronesi. 2018. Effects of winter weather and mixing regime on the restoration of a deep perialpine lake (Lake Lugano, Switzerland and Italy). Hydrobiologia 824: 229–242. doi:10.1007/s10750-018-3575-2

38. Luo, J., X. Tan, K. Liu, and W. Lin. 2018. Survey of sulfur-oxidizing bacterial community in the Pearl River water using soxB, sqr, and dsrA as molecular biomarkers. 3 Biotech 8: 1–12. doi:10.1007/s13205-017-1077-y

39. McMurdie, P. J., and S. Holmes. 2013. phyloseq: an R package for reproducible interactive analysis and graphics of microbiome census data. PLoS One 8: 1–11. doi:10.1371/journal.pone.0061217

40. Mziray, P., I. A. Kimirei, P. A. Staehr, C. V Lugomela, W. L. Perry, D. Trolle, C. M. O. Reilly, and H. F. Mgana. 2018. Seasonal patterns of thermal stratification and primary production in the northern parts of Lake Tanganyika. J Great Lakes Res 44: 1209–1220. doi:10.1016/j.jglr.2018.08.015

41. Naqvi, S. W. A. and others. 2018. Methane stimulates massive nitrogen loss from freshwater reservoirs in India. Nat Commun 9: 1–10. doi:10.1038/s41467-018-03607-z

42. Oksanen, J. and others. 2020. R package vegan: community ecology package.

43. Ooyang, L., B. Thamdrup, and M. Trimmer. 2021. Coupled nitrification and N_2_ gas production as a cryptic process in oxic riverbeds. Nat Commun 1–8. doi:10.1038/s41467-021-21400-3

44. Oswald, K. and others. 2016. Methanotrophy under versatile conditions in the water column of the ferruginous meromictic Lake La Cruz (Spain). Front Microbiol 7. doi:10.3389/fmicb.2016.01762

45. Pajares, S., M. M. Macek, and J. Alcocer. 2017. Vertical and seasonal distribution of picoplankton and functional nitrogen genes in a high-altitude warm-monomictic tropical lake. Freshw Biol 62: 1180–1193. doi:10.1111/fwb.12935

46. Pandey, C. B., U. Kumar, M. Kaviraj, K. J. Minick, A. K. Mishra, and J. S. Singh. 2020. Science of the Total Environment DNRA: A short-circuit in biological N-cycling to conserve nitrogen in terrestrial ecosystems. Science of the Total Environment 738: 139710. doi:10.1016/j.scitotenv.2020.139710

47. Quast, C., E. Pruesse, P. Yilmaz, J. Gerken, T. Schweer, F. O. Glo, and P. Yarza. 2013. The SILVA ribosomal RNA gene database project: improved data processing and web-based tools. Nucleic Acids Res 41: 590–596. doi:10.1093/nar/gks1219

48. Raghoebarsing, A. A. and others. 2006. A microbial consortium couples anaerobic methane oxidation to denitrification. Nature 440: 918–921. doi:10.1038/nature04617

49. Risgaard-Petersen, N., S. Rysgaard, and N. P. Revsbech. 1995. Combined microdiffusion-hypobromite oxidation method for determining nitrogen-15 isotope in ammmonium. Soil Science Society of America Journal 59: 1077–1080. doi:10.2136/sssaj1995.03615995005900040018x

50. Roland, F. A. E., F. Darchambeau, A. V. Borges, C. Morana, L. De Brabandere, B. Thamdrup, and S. A. Crowe. 2018. Denitrification, anaerobic ammonium oxidation, and dissimilatory nitrate reduction to ammonium in an East African Great Lake (Lake Kivu). Limnol Oceanogr 63: 687–701. doi:10.1002/lno.10660

51. Rosenberg, E., E. F. DeLong, S. Lory, E. Stackebrandt, and F. Thompson. 2014. The Prokaryotes: Deltaproteobacteria and Epsilonproteobacteria, 4th ed. Springer.

52. Sakuola, D., H. Koch, J. Frank, M. S. M. Jetten, M. A. H. J. van Kessel, and S. Lücker. 2021. Enrichment and physiological characterization of a novel comammox *Nitrospira* indicates ammonium inhibition of complete nitrification. ISME J 15: 1010–1024. doi:10.1038/s41396-020-00827-4

53. Schubert, C. J., E. Durisch-Kaiser, B. Wehrli, B. Thamdrup, P. Lam, and M. M. M. Kuypers. 2006. Anaerobic ammonium oxidation in a tropical freshwater system (Lake Tanganyika). Environ Microbiol 8: 1857–1863. doi:10.1111/j.1462-2920.2006.001074.x

54. Seitzinger, S. P. 1988. Denitrification in freshwater and coastal marine ecosystems: Ecological and geochemical significance. Limnol Oceanogr 33: 702–724. doi:10.1057/9780230306912

55. Shao, M. F., T. Zhang, and H. H. P. Fang. 2010. Sulfur-driven autotrophic denitrification: Diversity, biochemistry, and engineering applications. Appl Microbiol Biotechnol 88: 1027–1042. doi:10.1007/s00253-010-2847-1

56. Shapleigh, J. P. 2013. Denitrifying Prokaryotes, p. 405–425. In E. et Al Rosenberg [ed.], The Prokaryotes – Prokaryotic Physiology and Biochemistry. Springer-Verlag Berlin Heidelberg.

57. Shaw, L. J., G. W. Nicol, Z. Smith, J. Fear, J. I. Prosser, and E. M. Baggs. 2006. *Nitrosospira* spp. can produce nitrous oxide via a nitrifier denitrification pathway. Environ Microbiol 8: 214–222. doi:10.1111/j.1462-2920.2005.00882.x

58. Studer, A. S., L. Wörmer, H. Vogel, N. Dubois, M. Bartosiewicz, K. U. Hinrichs, F. Lepori, and M. F. Lehmann. 2024. First lacustrine application of the diatom-bound nitrogen isotope paleo-proxy reveals coupling of denitrification and N_2_ fixation in a hyper-eutrophic lake. Limnol Oceanogr 1–13. doi:10.1002/lno.12627

59. Su, G., M. F. Lehmann, J. Tischer, Y. Weber, F. Lepori, J.-C. Walser, H. Niemann, and J. Zopfi. 2023. Water column dynamics control nitrite-dependent anaerobic methane oxidation by *Candidatus* “Methylomirabilis” in stratified lake basins. ISME J 17: 693– 702. doi:10.1038/s41396-023-01382-4

60. Su, G., J. Zopfi, H. Yao, L. Steinle, H. Niemann, and M. F. Lehmann. 2020. Manganese/iron-supported sulfate-dependent anaerobic oxidation of methane by archaea in lake sediments. Limnol Oceanogr 65: 863–875. doi:10.1002/lno.11354

61. Thamdrup, B., and T. Dalsgaard. 2002. Production of N_2_ through anaerobic ammonium oxidation coupled to nitrate reduction in marine sediments. Appl Environ Microbiol 68: 1312–1318. doi:10.1128/AEM.68.3.1312

62. Thamdrup, B., T. Dalsgaard, M. M. Jensen, O. Ulloa, L. Farias, and R. Escribano. 2006. Anaerobic ammonium oxidation in the oxygen-deficient waters off northern Chile. Limnol Oceanogr 51: 2145–2156. doi:10.4319/lo.2006.51.5.2145

63. Tischer, J. and others. 2022. Isotopic signatures of biotic and abiotic N_2_O production and consumption in the water column of meromictic, ferruginous Lake La Cruz (Spain). Limnol Oceanogr 67: 1760–1775. doi:10.1002/lno.12165

64. Tischer, J., J. Zopfi, C. Frey, J. Venetz, L. Burgdorfer, O. Rehmann, and M. F. Lehmann. 2025. *Isotope effects of nitrate reduction by natural lake water communities* [Manuscript in preparation]. Department of Environmental Sciences, University of Basel.

65. Wenk, C. B., J. Blees, J. Zopfi, M. Veronesi, A. Bourbonnais, C. J. Schubert, H. Niemann, and M. F. Lehmann. 2013. Anaerobic ammonium oxidation (anammox) bacteria and sulfide-dependent denitrifiers coexist in the water column of a meromictic south-alpine lake. Limnology Oceanography 58: 1–12. doi:10.4319/lo.2013.58.1.0001

66. Wenk, C. B., J. Zopfi, J. Blees, M. Veronesi, H. Niemann, and M. F. Lehmann. 2014. Community N and O isotope fractionation by sulfide-dependent denitrification and anammox in a stratified lacustrine water column. Geochim Cosmochim Acta 125: 551–563. doi:10.1016/j.gca.2013.10.034

67. Wickham, H. 2016. ggplot2: elegant graphics for data analysis, Springer-Verlag New York.

68. Wickham, H., R. François, L. Henry, and K. Müller. 2021. R package dplyr: A Grammar of Data Manipulation.

69. Wrage, N., G. L. Velthof, O. Oenema, and H. J. Laanbroek. 2004. Acetylene and oxygen as inhibitors of nitrous oxide production in *Nitrosomonas europaea* and *Nitrosospira briensis*: a cautionary tale. FEMS Microbiol Ecol 47: 13–18. doi:10.1016/S0168-6496(03)00220-4

70. Wüest, A., W. Aeschbach-Hertig, H. Baur, M. Hofer, R. Kipfer, and M. Schurter. 1992. Density structure and tritium-helium age of deep hypolimnetic water in the northern basin of Lake Lugano. Aquat Sci 54: 205–218. doi:10.1007/BF00878137

71. Yang, Y. and others. 2021. The evolution pathway of ammonia-oxidizing archaea shaped by major geological events. Mol Biol Evol 38: 3637–3648. doi:10.1093/molbev/msab129

72. Yao, X. and others. 2024. Methane-dependent complete denitrification by a single *Methylomirabilis* bacterium. Nat Microbiol 9: 464–476. doi:10.1038/s41564-023-01578-6

73. Zopfi, J., M. E. Böttcher, and B. B. Jørgensen. 2008. Biogeochemistry of sulfur and iron in Thioploca-colonized surface sediments in the upwelling area off Central Chile. Geochim Cosmochim Acta 72: 827–843. doi:10.1016/j.gca.2007.11.031

74. Zumft, W. G. 1997. Cell biology and molecular basis of denitrification. Microbiology and molecular biology reviews 61: 533–616. doi:10.1128/.61.4.533-616.1997

