## Supporting Information for "Microbial nitrogen removal versus recycling in the redox transition zone of a meromictic lake and its coupling to sulfur"

### Table of content

**Table S1.** Taxa associated to the N-, S-, and C-cycles.
**Table S2.** Nutrient fluxes ($F_Z$) to the redox transition zone (RTZ) and in situ rates necessary to explain these fluxes, assuming a 15 m thick reaction layer.
**Table S3.** S species consumption and production rates during the first 3-4 days of incubation with unlabeled $NO_3^-$ and $H_2S$.
**Figure S1.** Water column characteristics of the Lake Lugano North Basin.
**Figure S2.** Dissolved $O_2$ and chl *a* concentrations in the Lake Lugano North Basin.
**Figure S3.** Concentration profiles of N species, $H_2S$, and Fe species in the Lake Lugano North Basin.
**Figure S4.** Temporal evolution of the observed amplicon sequencing variant (ASV) richness ("Observed") in rarefied datasets from three adjacent redox zones in the water column of in the North Basin of Lake Lugano.
**Figure S5.** Composition of most abundant taxa in the water column of the Lake Lugano North Basin, based on amplicon sequencing variant (ASV) data from 2009 to 2018.
**Figure S6.** Time series of most abundant taxa in the Lake Lugano North Basin, based on amplicon sequencing variant (ASV) data from 2009 to 2018.
**Figure S7.** Time series of relative-abundance profiles of amplicon sequence variants (ASVs) of ammonia-oxidizing archaea.
**Figure S8**. Time series of relative-abundance profiles of amplicon sequencing variants (ASVs) of nitrite-oxidizing and complete ammonia oxidation (comammox) performing *Nitrospira* sp.
**Figure S9.** Time series of relative-abundance profiles of amplicon sequence variants (ASVs) of potential anammox bacteria.
**Figure S10.** Time series of relative-abundance profiles of amplicon sequence variants (ASVs) of potential N-reducing bacteria.
**Figure S11.** Time series of relative-abundance profiles of amplicon sequencing variants (ASVs) of potential S-reducing bacteria and syntrophs.
**Figure S12.** $NO_3^-$ reduction rates of incubation experiments with added unlabeled $NO_3^-$.
**Figure S13.** Temporal dynamics of $H_2S$, $SO_4^{2-}$, $S^0$, and $S_2O_3^{2-}$ in incubation experiments with unlabeled $NO_3^-$ und $H_2S$ added to anoxic water.
**Figure S14.** Exemplary data of an $^{15}N$-label incubation experiment from April 2018, 95 m.

**Table S1.** Taxa associated to the N-, S-, and C-cycles, including ammonia-oxidizing archaea (AOA), ammonia-oxidizing bacteria (AOB), nitrite-oxidizing bacteria (NOB), anammox bacteria, N-reducing bacteria, and S-reducing bacteria in the Lake Lugano North Basin, based on amplicon sequence variant (ASV) data from 2009 to 2018. Only taxa with at least 0.005% relative abundance in at least one of the samples were included. Taxa in bold have a maximum relative abundance of >0.5% and are considered key taxa of the Lake Lugano water-column communities across the redox transition zone (RTZ). The list presents all detected taxa according to the search criteria outlined in the *Methods* section. Max. RA = maximum relative abundance of the taxon in the analyzed in situ samples.

| Type of organism/ metabolism | Genus | Family | Order | Class | Phylum | Max. RA (%) | Literature/Search criterion |
|---|---|---|---|---|---|---|---|
| **AOA** | ***Candidatus* Nitrosopumilus** | **Nitrosopumilaceae** | **Nitrosopumilales** | **Nitrososphaeria** | **Crenarchaeota** | **26.2** | Yang et al. 2021 |
| **AOB** | ***Nitrosomonas*** | **Nitrosomonadaceae** | **Burkholderiales** | **Gammaproteobacteria** | **Proteobacteria** | **0.955** | Yang et al. 2021 |
| AOB | *Nitrosospira* | Nitrosomonadaceae | Burkholderiales | Gammaproteobacteria | Proteobacteria | 0.015 | Yang et al. 2021 |
| **AOB** | **-** | **Nitrosomonadaceae** | **Burkholderiales** | **Gammaproteobacteria** | **Proteobacteria** | **4.2** | Yang et al. 2021 |
| **NOB** | ***Nitrospira*** | **Nitrospiraceae** | **Nitrospirales** | **Nitrospiria** | **Nitrospirota** | **3.1** | Yang et al. 2021 |
| **Anammox** | ***Candidatus* Anammoximicrobium** | Pirellulaceae | Pirellulales | **Planctomycetes** | **Planctomycetota** | **0.87** | Jetten et al. 2009 |
| Anammox | - | Brocadiaceae | Brocadiales | Brocadiae | Planctomycetota | 0.040 | Jetten et al. 2009 |
| Organotrophic N-reducer | *Hyphomicrobium* | Hyphomicrobiaceae | Rhizobiales | Alphaproteobacteria | Proteobacteria | 0.083 | Martineau et al. 2015 |
| Organotrophic N-reducer | *Rhodobacter* | Rhodobacteraceae | Rhodobacterales | Alphaproteobacteria | Proteobacteria | 0.17 | Zumft 1997 |
| Organotrophic N-reducer | *Shewanella* | Shewanellaceae | Alteromonadales | Gammaproteobacteria | Proteobacteria | 0.079 | Pandey et al. 2020 |
| S-oxidizing N-reducer | *Beggiatoa*[a] | Beggiatoaceae | Beggiatoales | Gammaproteobacteria | Proteobacteria | 0.007 | Shao et al. 2010 |
| **Organotrophic N-reducer** | ***Denitratisoma*** | **Rhodocyclaceae** | **Burkholderiales** | **Gammaproteobacteria** | **Proteobacteria** | **4.1** | Kojima and Fukui 2011 |
| Organotrophic N-reducer | *Candidatus* Accumulibacter | Rhodocyclaceae | Burkholderiales | Gammaproteobacteria | Proteobacteria | 0.26 | Sparacino-Watkins et al. 2014 |
| **Organotrophic N-reducer** | ***Sterolibacterium*** | **Rhodocyclaceae** | **Burkholderiales** | **Gammaproteobacteria** | **Proteobacteria** | **1.0** | Kojima and Fukui 2011 |
| Organotrophic N-reducer | *Thauera* | Rhodocyclaceae | Burkholderiales | Gammaproteobacteria | Proteobacteria | 0.009 | Xia et al. 2019 |
| **C+S-ox. N-reducer** | ***Dechloromonas*** | **Rhodocyclaceae** | **Burkholderiales** | **Gammaproteobacteria** | **Proteobacteria** | **1.9** | Salinero et al. 2009, incubation experiment |
| **S-oxidizing N-reducer** | ***Sulfuritalea*** | **Rhodocyclaceae** | **Burkholderiales** | **Gammaproteobacteria** | **Proteobacteria** | **1.0** | Kojima and Fukui 2011 |
| **Organotrophic N-reducer** | **-** | **Rhodocyclaceae** | **Burkholderiales** | **Gammaproteobacteria** | **Proteobacteria** | **5.2** | Kojima and Fukui 2011 |
| Organotrophic N-reducer | *Ralstonia* | Burkholderiaceae | Burkholderiales | Gammaproteobacteria | Proteobacteria | 0.034 | Zumft 1997 |

### Table S1 (continued)

| Type of organism/ metabolism | Genus | Family | Order | Class | Phylum | Max. RA (%) | Literature/Search criterion |
|---|---|---|---|---|---|---|---|
| S-oxidizing N-reducer | *Sulfuricella* | Sulfuricellaceae | Burkholderiales | Gammaproteobacteria | Proteobacteria | 0.010 | Kojima and Fukui 2011 |
| S-oxidizing N-reducer | *Thiobacillus*[a] | Hydrogenophilaceae | Burkholderiales | Gammaproteobacteria | Proteobacteria | 0.071 | Shao et al. 2010 |
| S/H-oxidizing N-reducer | *Hydrogenophaga* | Comamonadaceae | Burkholderiales | Gammaproteobacteria | Proteobacteria | 0.028 | incubation experiment, Willems et al. 1989 |
| **Organotrophic N-reducer** | ***Methylotenera*[a]** | **Methylophilaceae** | **Burkholderiales** | **Gammaproteobacteria** | **Proteobacteria** | **6.3** | Mustakhimov et al. 2013 |
| **$CH_4$-oxidizing N-reducer** | ***Methylobacter*[a]** | **Methylomonadaceae** | **Methylococcales** | **Gammaproteobacteria** | **Proteobacteria** | **16.3** | Kits et al. 2015 |
| **$CH_4$-oxidizing N-reducer** | ***Crenothrix*[b]** | **Methylomonadaceae** | **Methylococcales** | **Gammaproteobacteria** | **Proteobacteria** | **5.9** | Oswald et al. 2017 |
| Organotrophic N-reducer | *Halomonas* | Halomonadaceae | Oceanospirillales | Gammaproteobacteria | Proteobacteria | 0.020 | Zumft 1997 |
| **Organotrophic N-reducer** | ***Pseudomonas*[a]** | **Pseudomonadaceae** | **Pseudomonadales** | **Gammaproteobacteria** | **Proteobacteria** | **4.5** | Zumft 1997 |
| Organotrophic N-reducer | *Bdellovibrio* | Bdellovibrionaceae | Bdellovibrionales | Bdellovibrionia | Bdellovibrionota | 0.17 | Shapleigh 2013 |
| S-oxidizing N-reducer | *Sulfurimonas* | Sulfurimonadaceae | Campylobacterales | Campylobacteria | Campilobacterota | 0.016 | Shao et al. 2010 |
| S-oxidizing N-reducer | *Sulfuricurvum* | Sulfurimonadaceae | Campylobacterales | Campylobacteria | Campilobacterota | 0.11 | Shao et al. 2010 |
| Organotrophic N-reducer | *Bacillus* | Bacillaceae | Bacillales | Bacilli | Firmicutes | 0.024 | Zumft 1997 |
| **Organotrophic N-reducer** | ***Flavobacterium*[a]** | **Flavobacteriaceae** | **Flavobacteriales** | **Bacteroidia** | **Bacteroidota** | **3.2** | Zumft 1997, incubation experiment |
| **$CH_4$-oxidizing N-reducer** | ***Candidatus* Methylomirabilis** | **Methylomirabilaceae** | **Methylomirabilales** | **Methylomirabilia** | **Methylomirabilota** | **8.5** | Haroon et al. 2013 |
| S-reducer | *Desulfosporosinus* | Desulfitobacteriaceae | Desulfitobacteriales | Desulfitobacteriia | Firmicutes | 0.020 | "Desulf" |
| S-reducer | *Desulfurispora* | Desulfotomaculales | Desulfotomaculales | Desulfotomaculia | Firmicutes | 0.041 | Kaksonen et al. 2007 |
| **S-reducer** | ***Desulfobacca*** | **Desulfobaccaceae** | **Desulfobaccales** | **Desulfobaccia** | **Desulfobacterota** | **4.6** | "Desulf" |
| **S-reducer** | **-** | **-** | **Desulfobacterales** | **Desulfobacteria** | **Desulfobacterota** | **6.5** | "Desulf" |
| S-reducer | *Desulfobulbus* | Desulfobulbaceae | Desulfobulbales | Desulfobulbia | Desulfobacterota | 0.031 | "Desulf" |
| S-reducer | *Desulfocapsa* | Desulfocapsaceae | Desulfobulbales | Desulfobulbia | Desulfobacterota | 0.16 | "Desulf" |
| S-reducer | *Desulfomonile* | Desulfomonilaceae | Desulfomonilales | Desulfomonilia | Desulfobacterota | 0.016 | "Desulf", incubation experiment |
| S-reducer | *Desulfovibrio* | Desulfovibrionaceae | Desulfovibrionales | Desulfovibrionia | Desulfobacterota | 0.083 | (Hao et al. 1996) |
| S-reducer | *Desulfomicrobium* | Desulfomicrobiaceae | Desulfovibrionales | Desulfovibrionia | Desulfobacterota | 0.019 | "Desulf" |
| Fe/S-reducer | *Geobacter* | Geobacteraceae | Geobacterales | Desulfuromonadia | Desulfobacterota | 0.093 | Kojima and Fukui 2011 |
| Syntroph fermenter | **-** | **-** | **Syntrophales** | **Syntrophia** | **Desulfobacterota** | **1.1** | "Desulf" |
| Syntroph fermenter | - | - | Syntrophobacterales | Syntrophobacteria | Desulfobacterota | 0.23 | Kuever 2014 |
| Syntroph fermenter | *Syntrophobacter* | Syntrophobacteraceae | Syntrophobacterales | Syntrophobacteria | Desulfobacterota | 0.005 | Shapleigh 2013 |

[a] Not all known species of this taxon are known for performing N-reduction or possessing N-reduction genes, respectively.
[b] N-reduction coupled to $CH_4$-oxidation by *Crenothrix* sp. was suggested in Oswald et al. 2017, but has not been confirmed yet.

**Table S2.** Nutrient fluxes ($F_Z$) to the redox transition zone (RTZ) and in situ rates necessary to explain these fluxes, assuming a 15 m thick reaction layer.

| **Sampling timepoint** | **Flux of compound $F_Z$ (µmol $d^{-1}$ $m^{-2}$)** | | | | **Required net in situ consumption rate (µmol $L^{-1}$ $d^{-1}$)** | | | |
|---|---|---|---|---|---|---|---|---|
| | **$NO_3^-$** | **$NH_4^+$** | **$H_2S$** | **$CH_4$** | **$NO_3^-$** | **$NH_4^+$** | **$H_2S$** | **$CH_4$** |
| April 2015 | 769 | 298 | 165 | | 0.051 | 0.020 | 0.011 | |
| June 2015 | 443 | 194 | 298 | | 0.030 | 0.013 | 0.020 | |
| October 2015 | 671 | 319 | 153 | | 0.045 | 0.021 | 0.010 | |
| March 2016 | 603 | 530 | 125 | | 0.040 | 0.035 | 0.008 | |
| September 2016 | 931 | 326 | 304 | | 0.062 | 0.022 | 0.020 | |
| November 2016 | 1150 | 577 | 241 | 460 | 0.077 | 0.038 | 0.016 | 0.031 |
| February 2017 | 567 | 319 | 119 | | 0.038 | 0.021 | 0.008 | |
| October 2017 | 1446 | 391 | 85 | 1304 | 0.096 | 0.026 | 0.006 | 0.087 |
| April 2018 | 868 | 460 | 299 | | 0.058 | 0.031 | 0.020 | |
| **Average** | **828** | **379** | **199** | **882** | **0.055** | **0.025** | **0.013** | **0.059** |

**Table S3.** S species consumption and production rates during the first 3-4 days of incubation with unlabeled nitrate and sulfide. Negative numbers indicate consumption, positive values indicate production. Values are presented with standard error of means. Non-significant rates are indicated by 'ns'. nd = not detectable, na = not analyzed.

| Sampling Date | Depth (m) | $H_2S$ (µmol $L^{-1}$ $d^{-1}$) | | $S^0$ (µmol $L^{-1}$ $d^{-1}$) | | $SO_3^{2-}$ (µmol $L^{-1}$ $d^{-1}$) | | $S_2O_3^{2-}$ (µmol $L^{-1}$ $d^{-1}$) | | $SO_4^{2-}$ (µmol $L^{-1}$ $d^{-1}$) | |
|---|---|---|---|---|---|---|---|---|---|---|---|
| November 2016 | 95 | -5.8 ± 0.5 | -4.4 ± 0.4 | 2.4 ± 0.9 (ns) | nd | na | | na | | na | |
| November 2016 | 105 | -4.7 ± 0.3 | -3.9 ± 0.6 | nd | 0.6 ± 0.3 (ns) | -0.03 ± 0.01 | nd | 0.18 ± 0.07 | 0.19 ± 0.05 | -3.3 ± 0.6 | -3.8 ± 0.3 |
| November 2016 | 155 | -1.9 ± 0.2 | -1.3 ± 0.4 | nd | nd | nd | nd | 0.16 ± 0.08 (ns) | nd | 1.0 ± 0.3 | -0.7 ± 0.2 |
| February 2017 | 105 | -2.7 ± 0.2 | -1-9 ± 0.1 | na | | na | | na | | na | |
| October 2017 | 105 | -6.8 ± 1.0 | -5.6 ± 0.9 | na | | na | | na | | na | |
| April 2018 | 95 | -9 ± 1.8 | -7.4 ± 1.9 | na | | na | | na | | na | |
| **Average** | | **-4.7 ± 0.7** | | | | | | | | | |

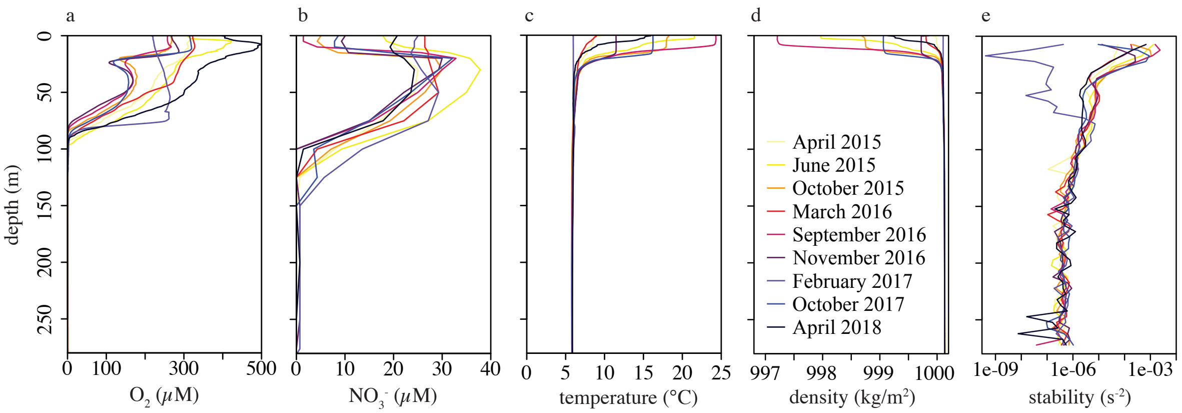


**Figure S1. Water column characteristics of the Lake Lugano North Basin. a)** Dissolved $O_2$ concentrations, **b)** $NO_3^-$ concentrations, **c)** temperature, **d)** density, and **e)** water column stability (Brunt-Väisälä frequency $N^2$) calculated according to Wüest et al. (1992).

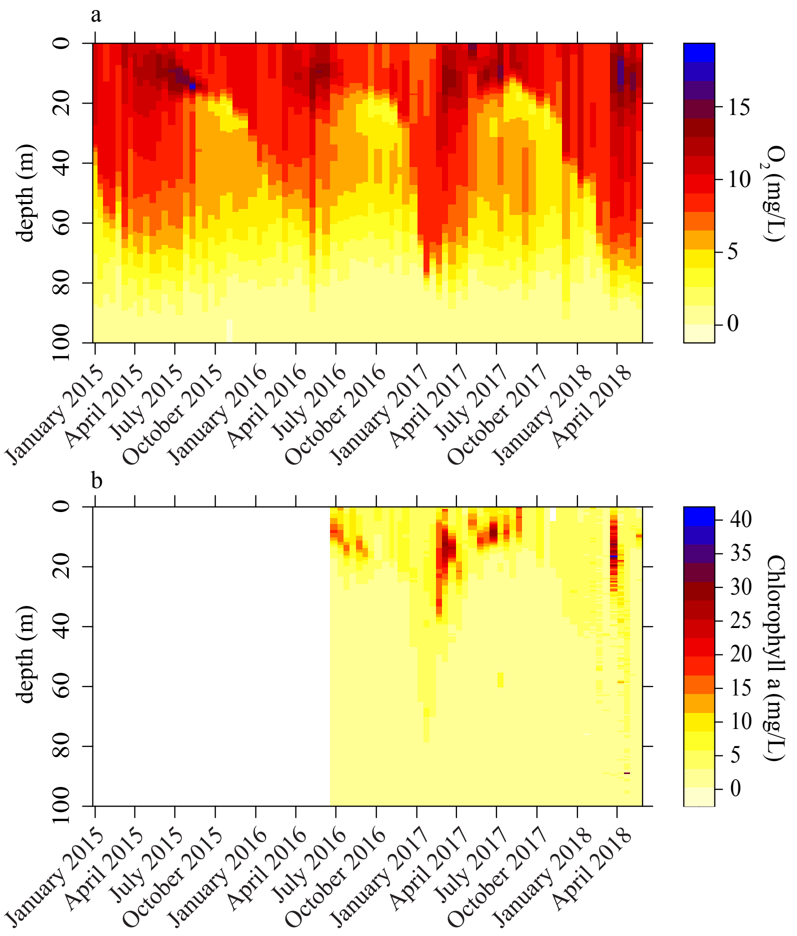


**Figure S2. a)** Dissolved $O_2$ from 2015 to mid-2018 and **b)** chl *a* concentrations from mid-2016 to mid-2018 in the upper 100 m of the water column of the Lake Lugano North Basin.

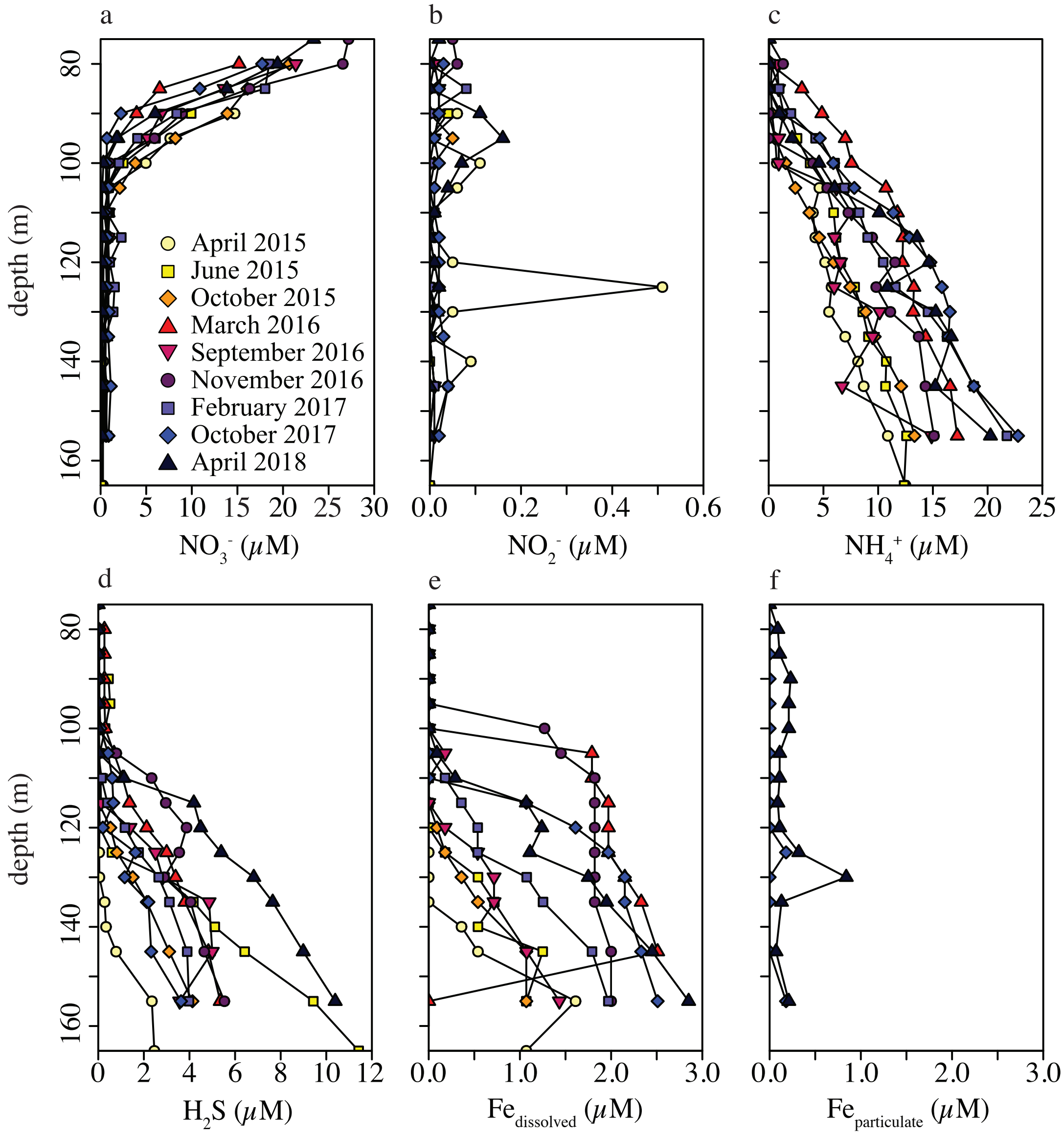


**Figure S3.** Concentration profiles of N species, $H_2S$, and Fe species in the Lake Lugano North Basin. **a)** $NO_3^-$, **b)** $NO_2^-$, **c)** $NH_4^+$, **d)** $H_2S$ (bimane method: April 2015, October 2015, March 2016, September 2016, October 2016; Cline method: June 2015, November 2016, February 2017, April 2018), **e)** $Fe_{dissolved}$, and **f)** $Fe_{particulate}$ (only October 2017 and April 2018).

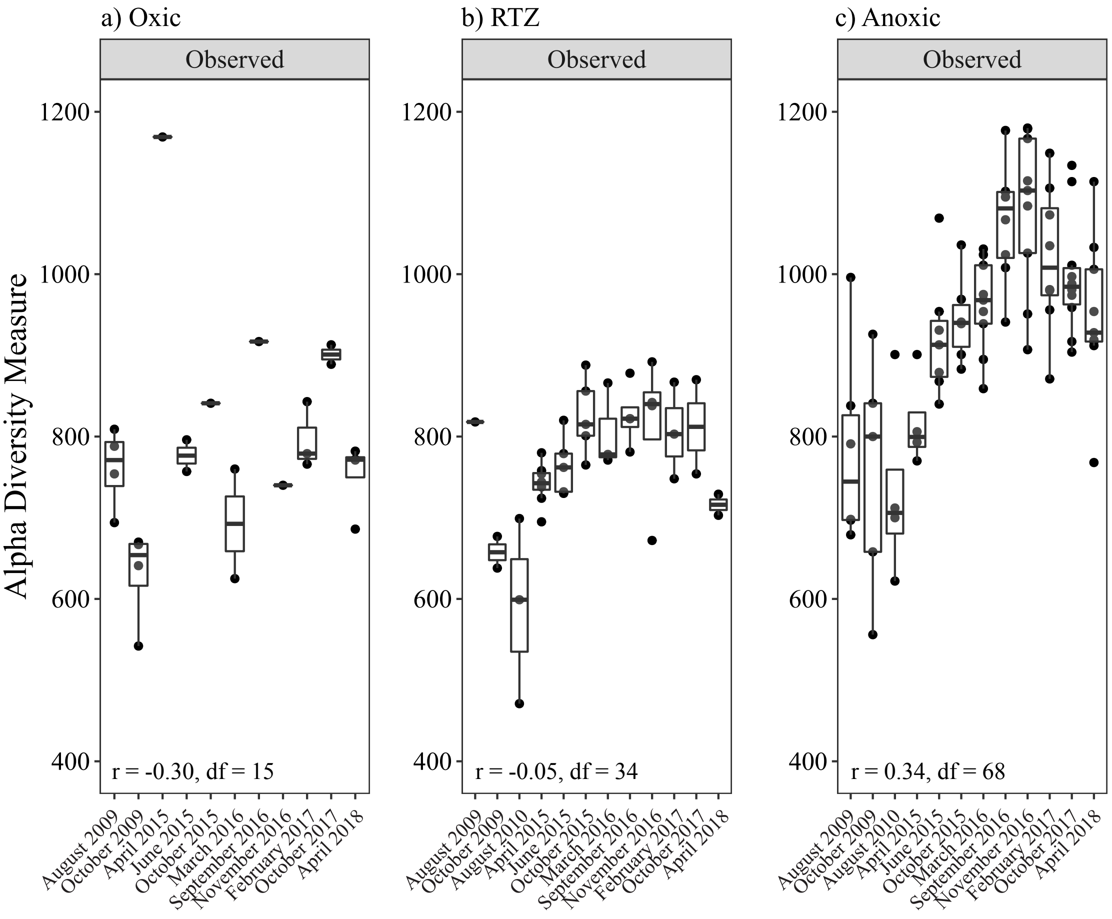


**Figure S4.** Temporal evolution of the observed amplicon sequencing variant (ASV) richness ("Observed") in rarefied datasets from three adjacent redox zones in the water column of in the North Basin of Lake Lugano: **a)** Oxic, **b)** redox transition zone "RTZ", and **c)** anoxic. The Pearson correlation coefficients (r) are given including degrees of freedom (df).

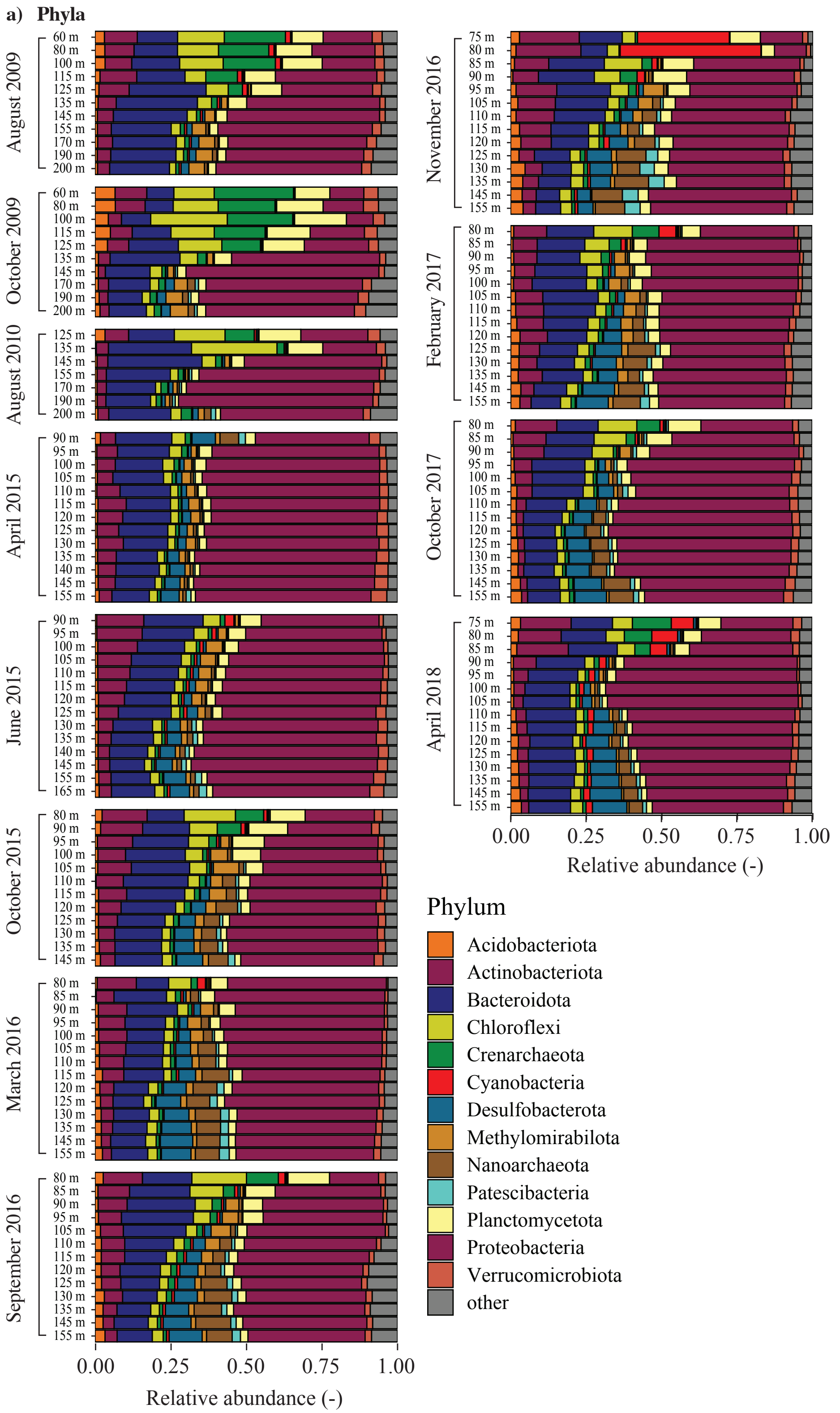


**Figure S5.** Composition of most abundant taxa in the water column of the Lake Lugano North Basin, based on amplicon sequencing variant (ASV) data from 2009 to 2018. Taxa that did not reach a relative abundance of 1% in at least one of the samples are summarized in “other”. **a)** Phyla

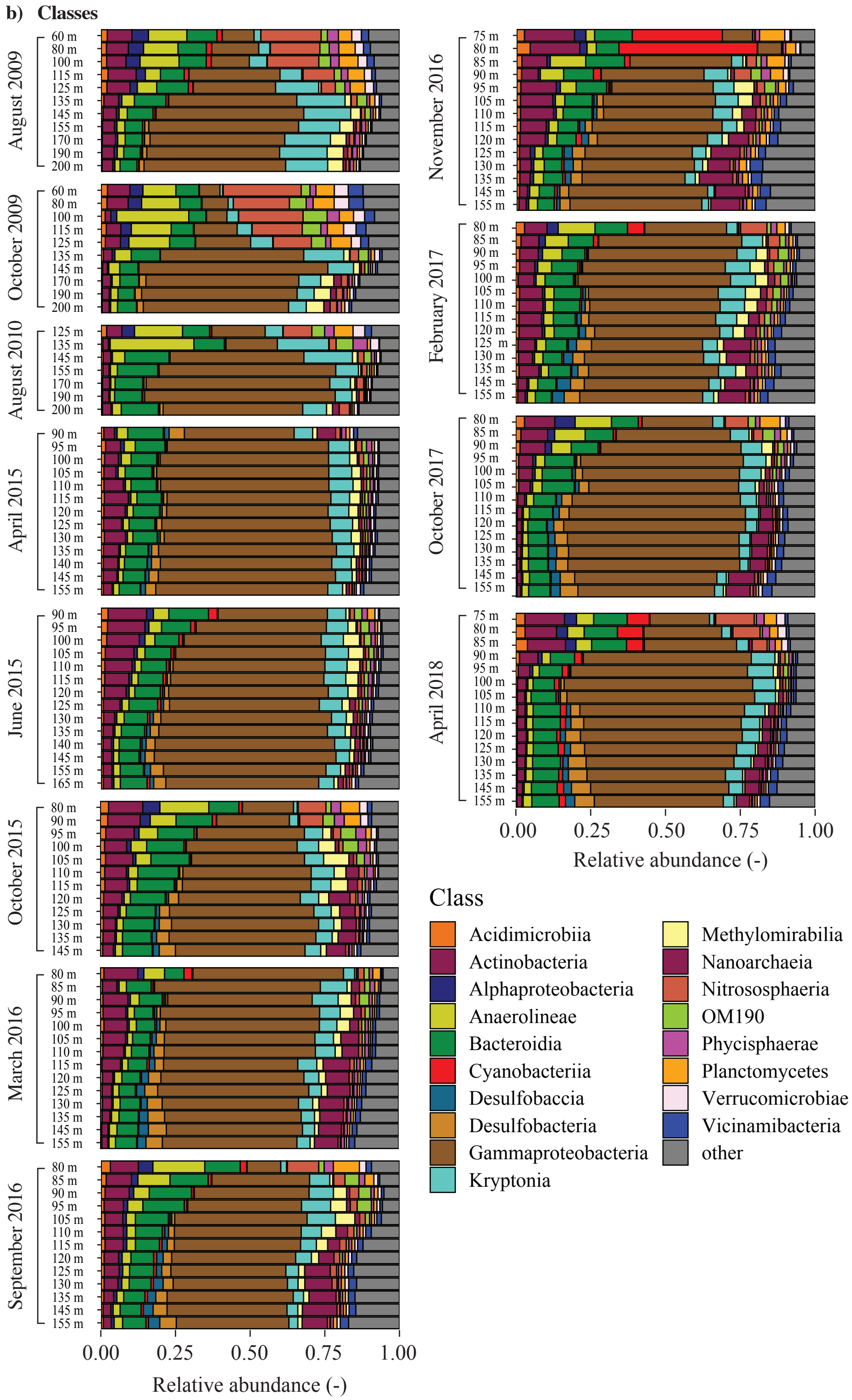


**Figure S5 (continued).** Composition of most abundant taxa in the water column of the Lake Lugano North Basin, based on amplicon sequencing variant (ASV) data from 2009 to 2018. Taxa that did not reach a relative abundance of 1% in at least one of the samples are summarized in "other". **b)** Classes

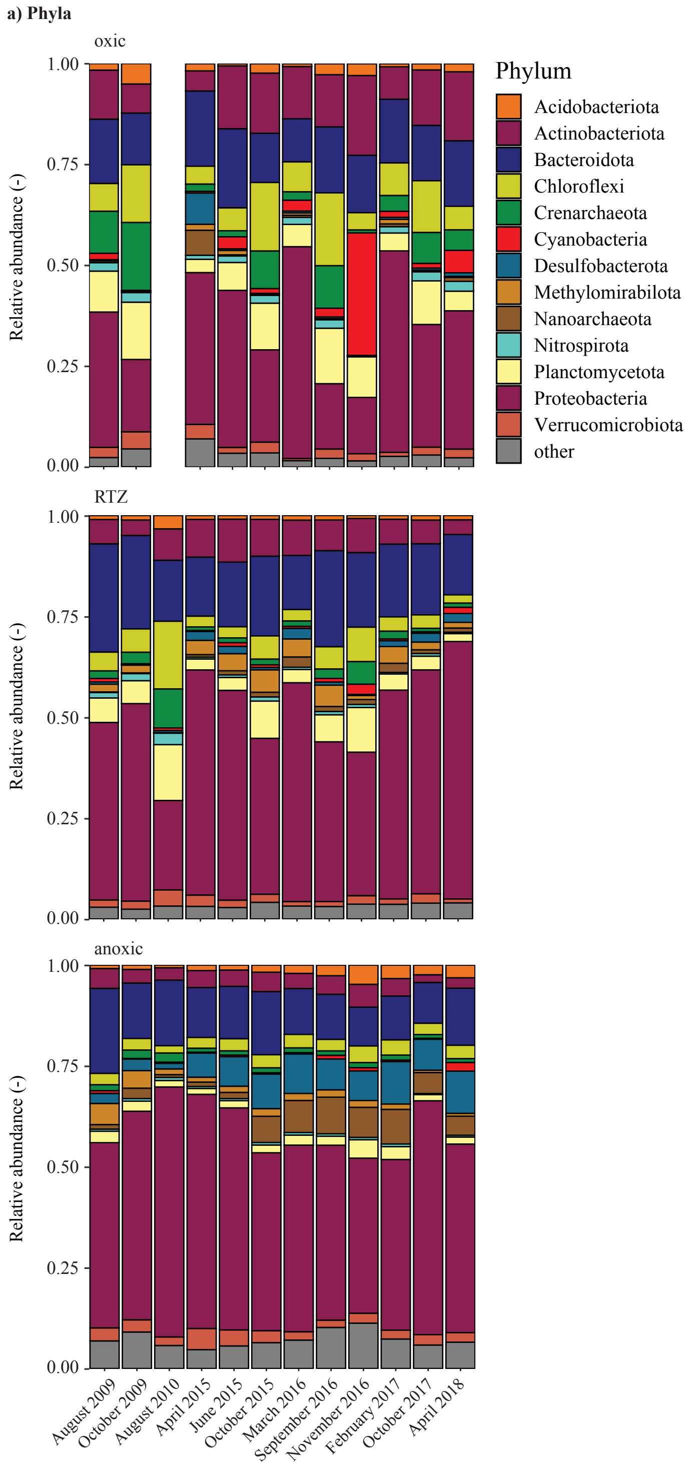


**Figure S6.** Time series of most abundant taxa in the tree different compartments (oxic, redox transition zone "RTZ", anoxic) in the Lake Lugano North Basin, based on amplicon sequencing variant (ASV) data from 2009 to 2018. Taxa that did not reach a relative abundance of 1% in at least one of the samples are summarized in "other". From each time point, one oxic sample, one RTZ sample, and one anoxic sample (see main text) was selected. **a)** Phyla

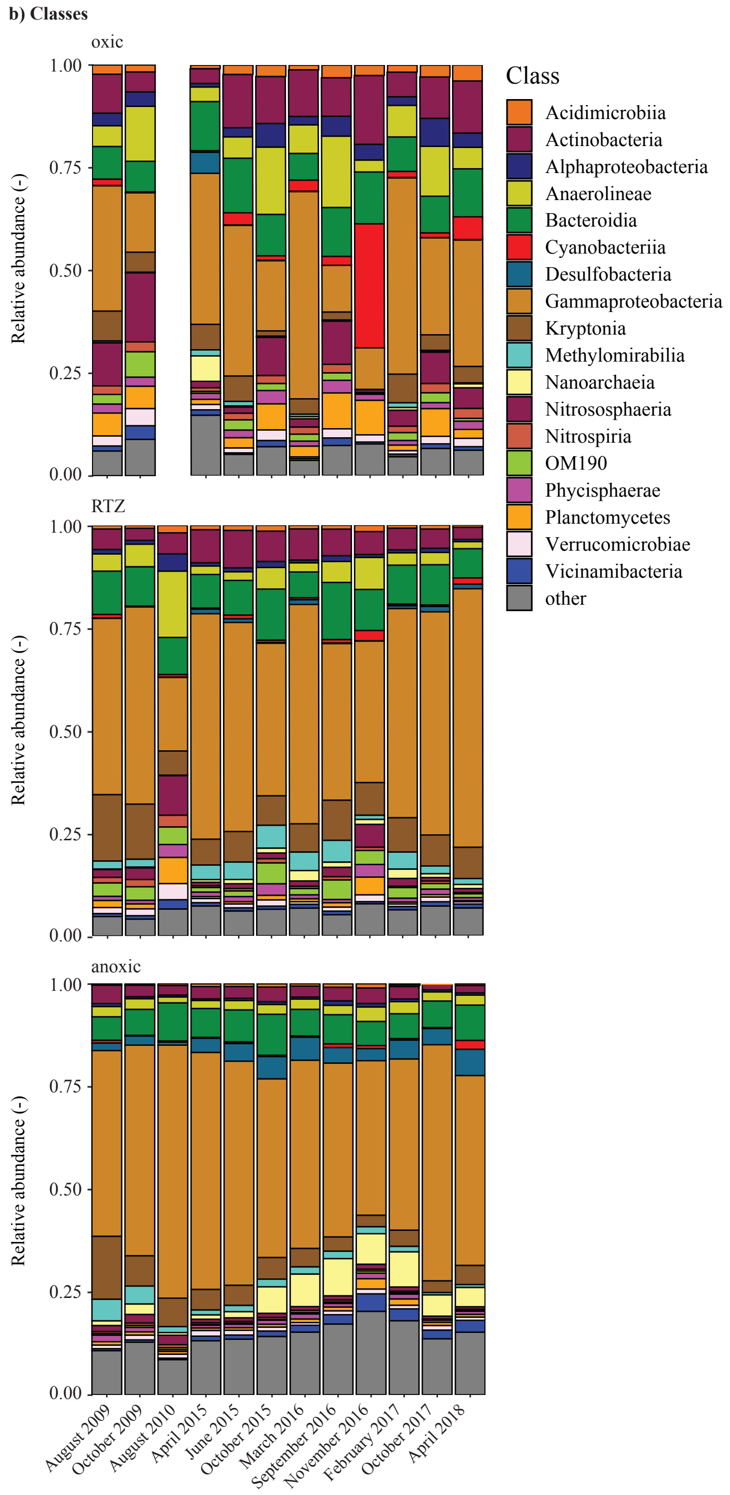


**Figure S6 (continued).** Time series of most abundant taxa in the tree different compartments (oxic, redox transition zone "RTZ", anoxic) in the Lake Lugano North Basin, based on amplicon sequencing variant (ASV) data from 2009 to 2018. Taxa that did not reach a relative abundance of 1% in at least one of the samples are summarized in "other". From each time point, one oxic sample, one RTZ sample, and one anoxic sample (see main text) was selected. **b)** Classes

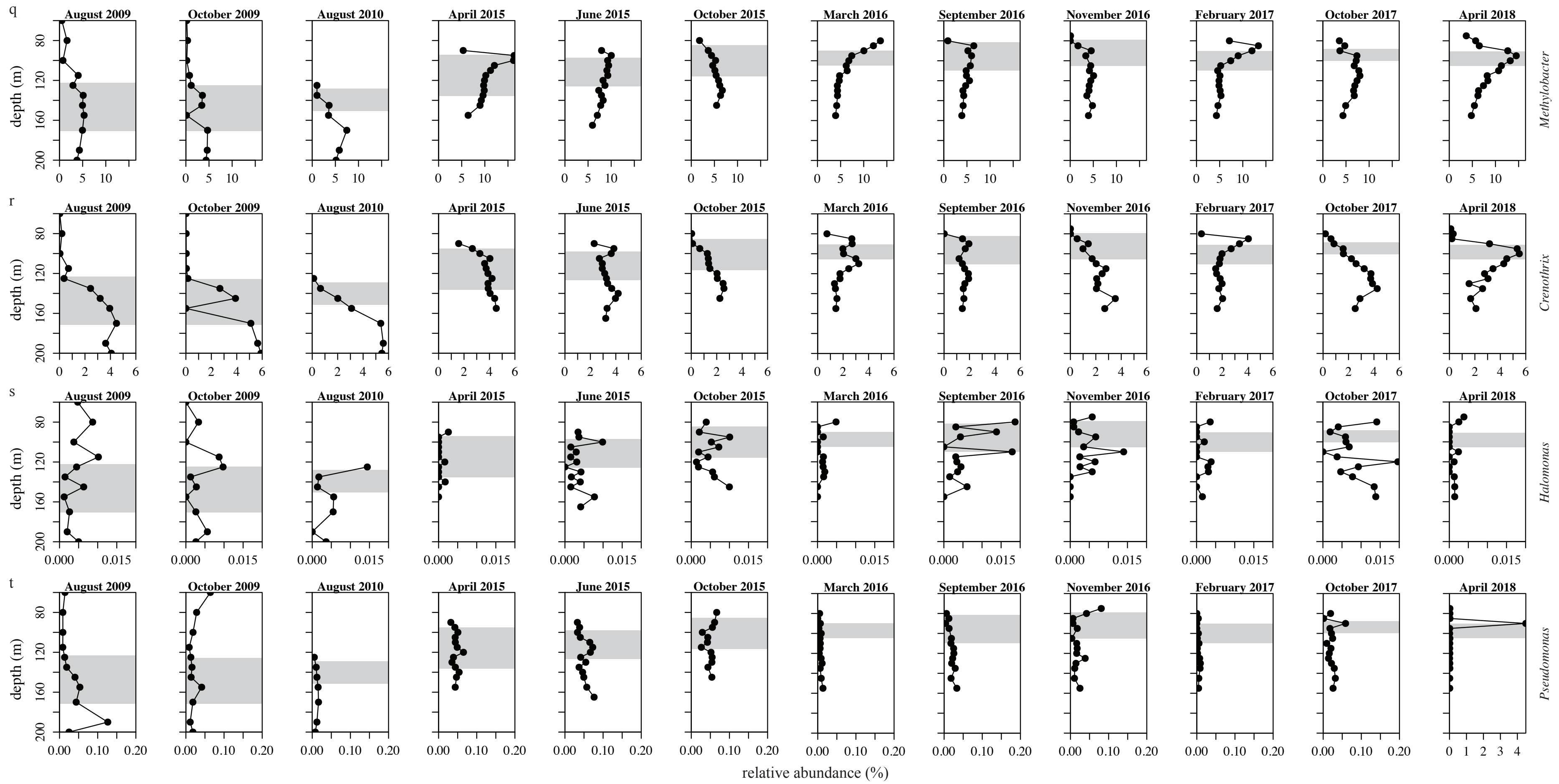


**Figure S7.** Time series of relative-abundance profiles of amplicon sequencing variants (ASVs) of ammonia-oxidizing archaea. **a)** *Ca.* Nitrosopumilus, and ammonia-oxidizing bacteria (AOB), **b)** *Nitrosomonas*, **c)** *Nitrosospira*, and **d)** *Nitrosomonadaceae*. Grey bars represent the redox transition zone (RTZ).

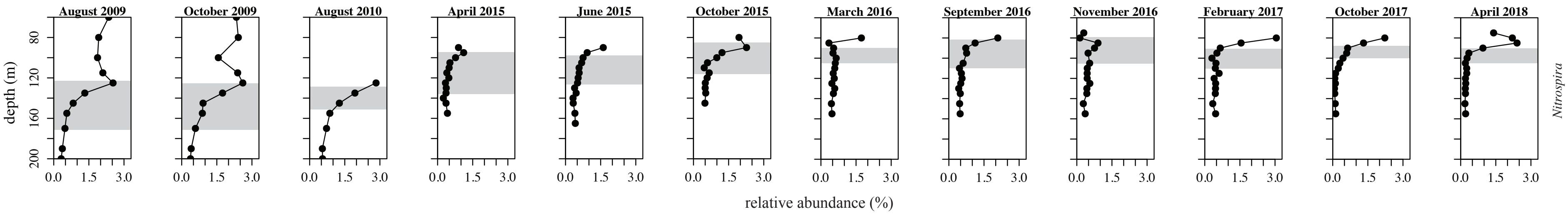


**Figure S8.** Time series of relative-abundance profiles of amplicon sequencing variants (ASVs) of nitrite-oxidizing and complete ammonia oxidation (comammox) performing *Nitrospira* sp. Grey bars represent the redox transition zone (RTZ).

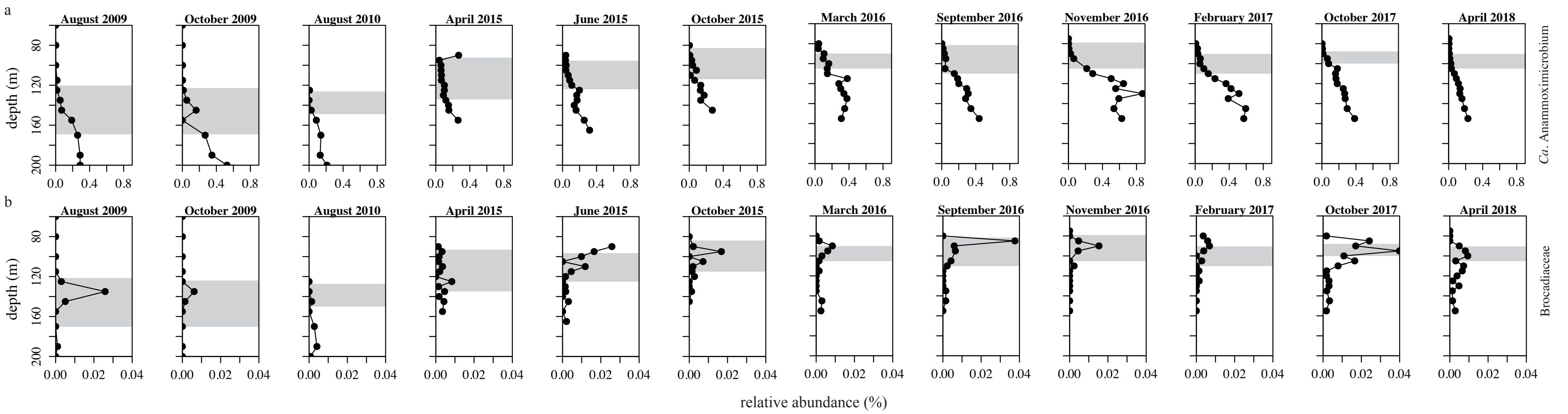


**Figure S9.** Time series of relative-abundance profiles of amplicon sequencing variants (ASVs) of potential anammox bacteria. **a)** *Ca.* Anammoximicrobium and **b)** Brocadiaceae. Grey bars represent the redox transition zone (RTZ).

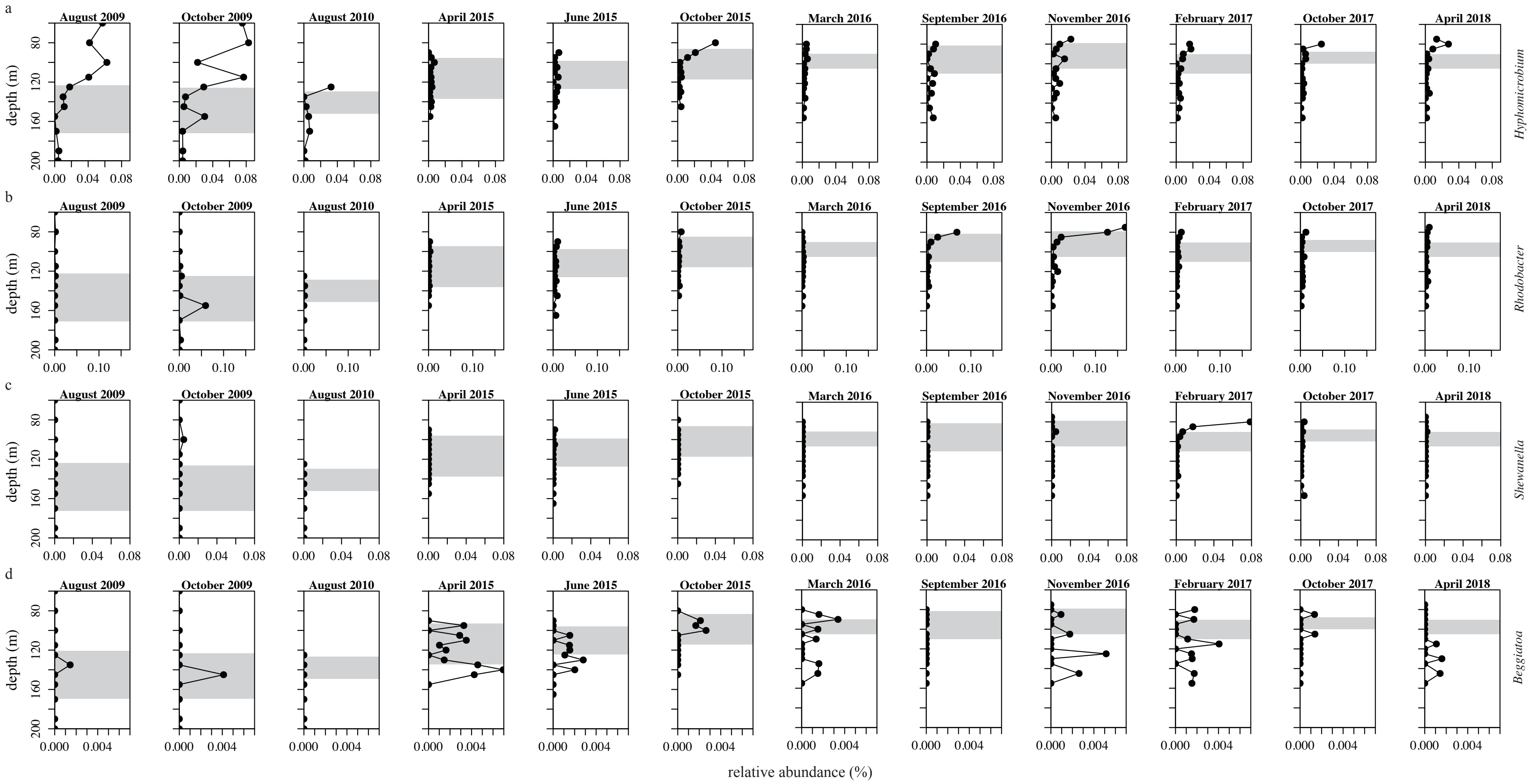

**Figure S10.** Time series of relative-abundance profiles of amplicon sequencing variants (ASVs) of potential N-reducing bacteria. **a)** *Hyphomicrobium*, **b)** *Rhodobacter*, **c)** *Shewanella*, **d)** *Beggiatoa*. Grey bars represent the redox transition zone (RTZ).

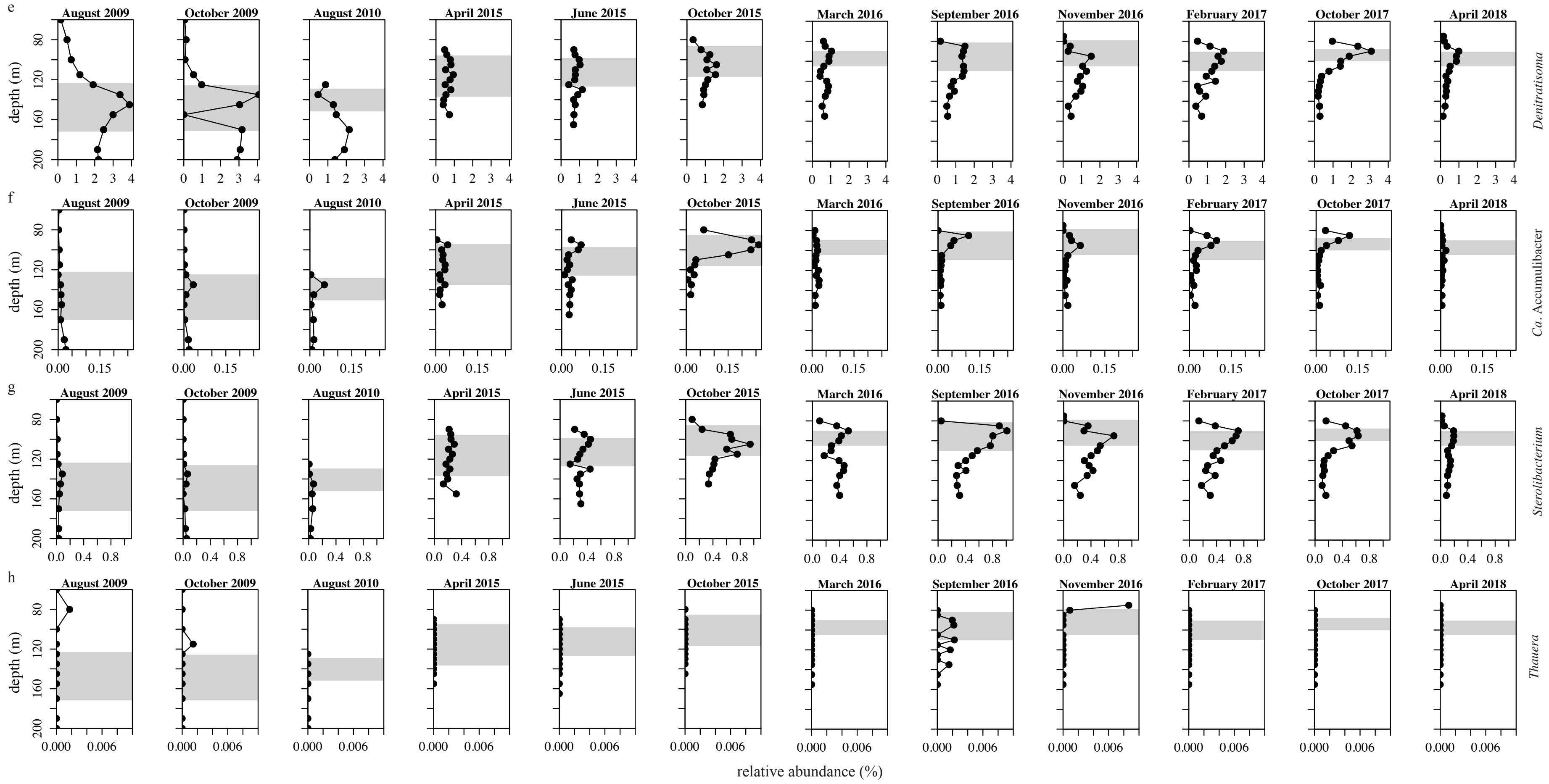


**Figure S10 (continued).** Time series of relative-abundance profiles of amplicon sequencing variants (ASVs) of potential N-reducing bacteria. **e)** *Denitratisoma*, **f)** *Ca.* Accumulibacter, **g)** *Sterolibacterium*, **h)** *Thauera*. Grey bars represent the RTZ.

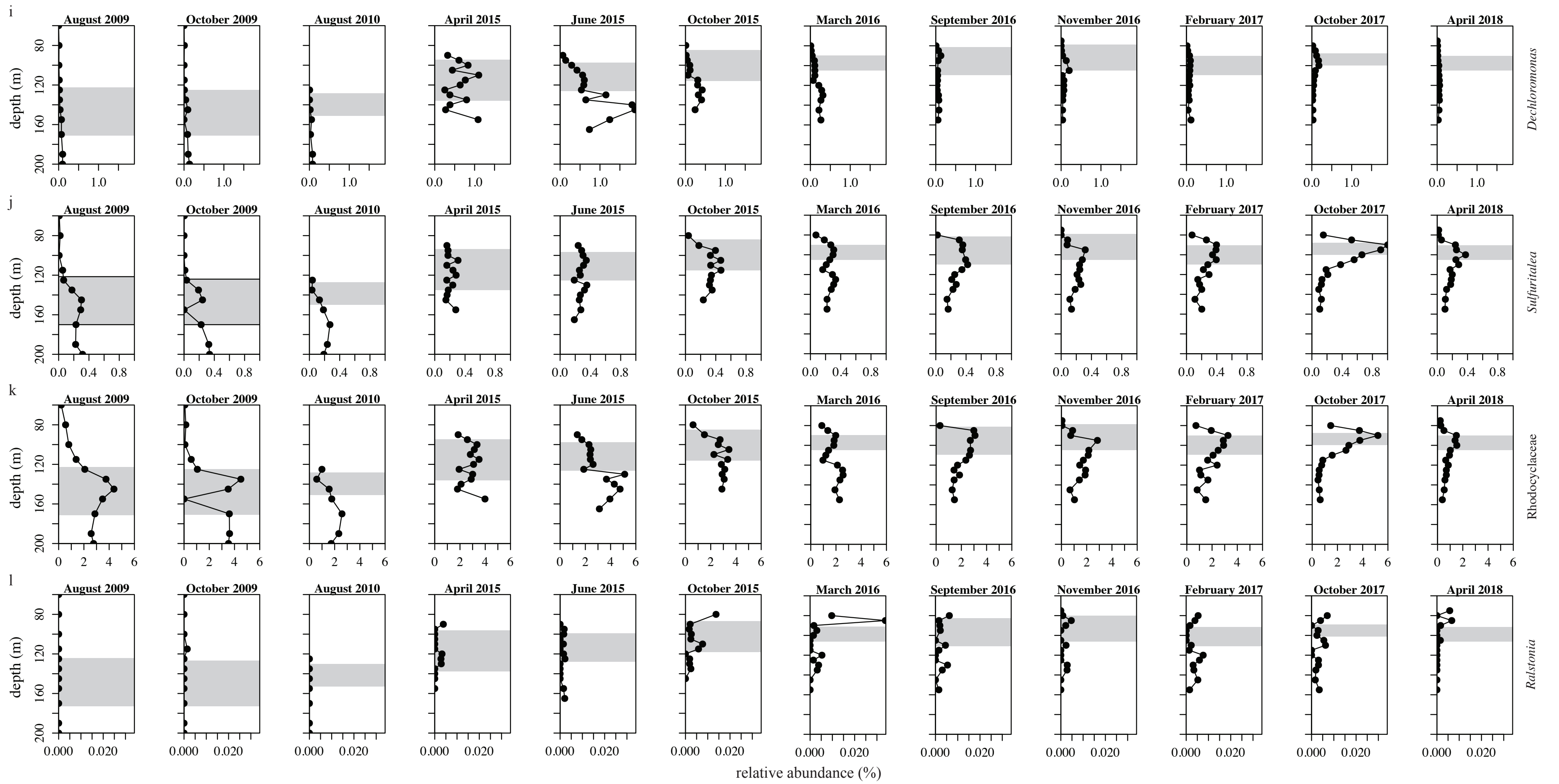


**Figure S10 continued.** Time series of relative-abundance profiles of amplicon sequencing variants (ASVs) of potential N-reducing bacteria. **i)** *Dechloromonas*, **j)** *Sulfuritalea*, **k)** Rhodocyclaceae, **l)** *Ralstonia*. Grey bars represent the RTZ.

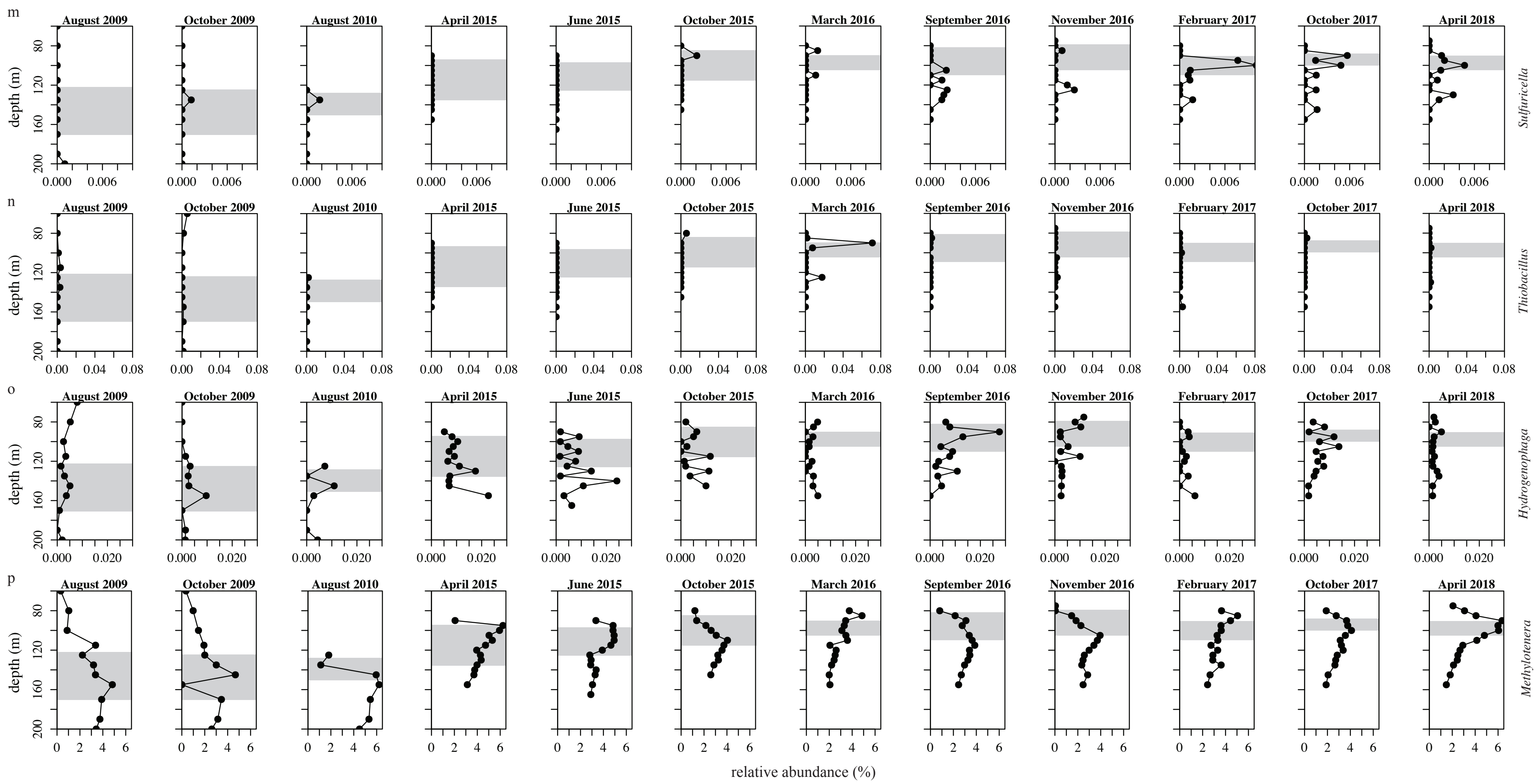


**Figure S10 continued.** Time series of relative-abundance profiles of amplicon sequencing variants (ASVs) of potential N-reducing bacteria. **m)** *Sulfuricella*, **n)** *Thiobacillus*, **o)** *Hydrogenophaga*, **p)** *Methylotenera*. Grey bars represent the RTZ.

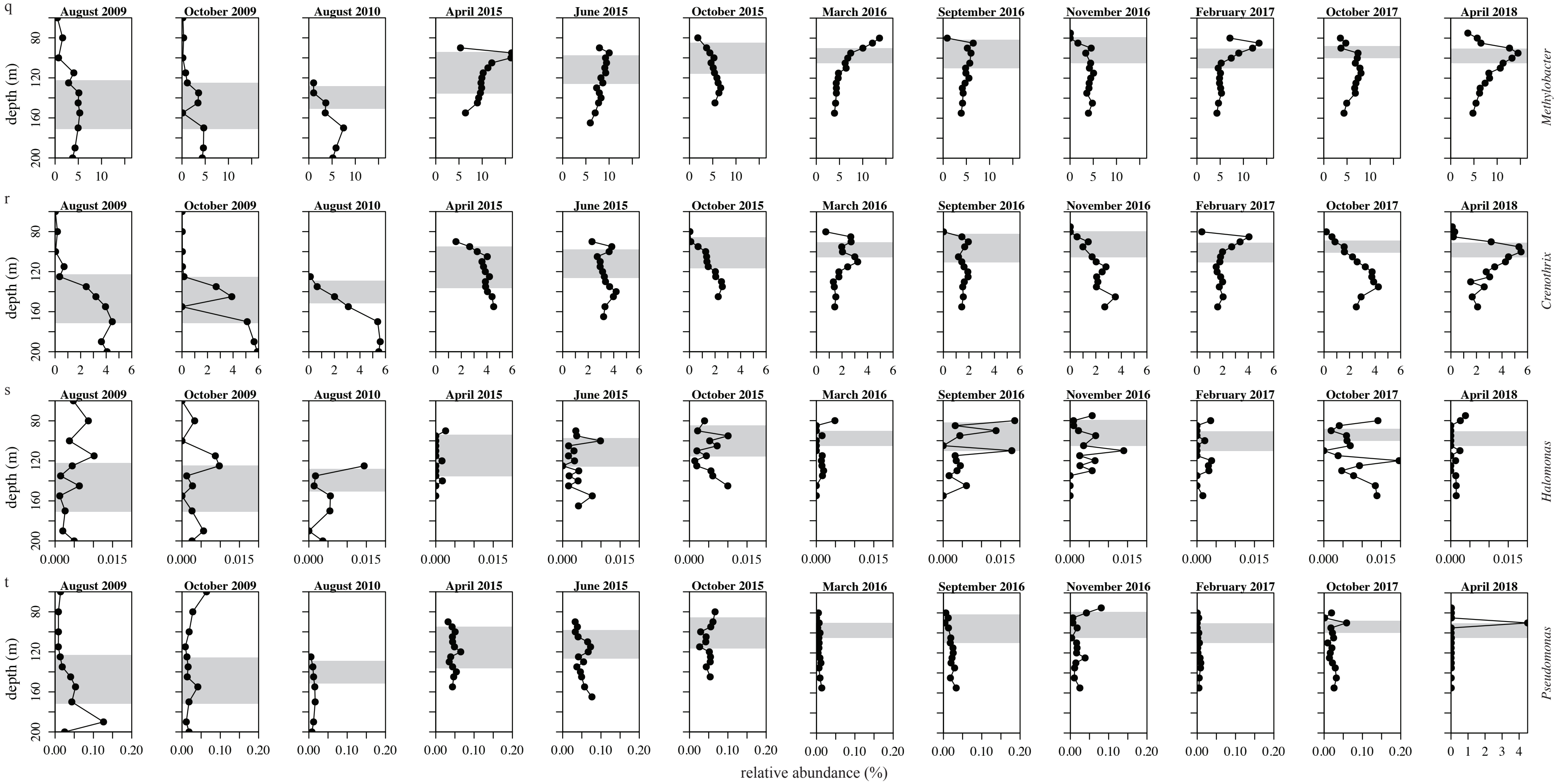


**Figure S10 (continued).** Time series of relative-abundance profiles of amplicon sequencing variants (ASVs) of potential N-reducing bacteria. **q)** *Methylobacter*, **r)** *Crenothrix*, **s)** *Halomonas*, **t)** *Pseudomonas*. Grey bars represent the RTZ.

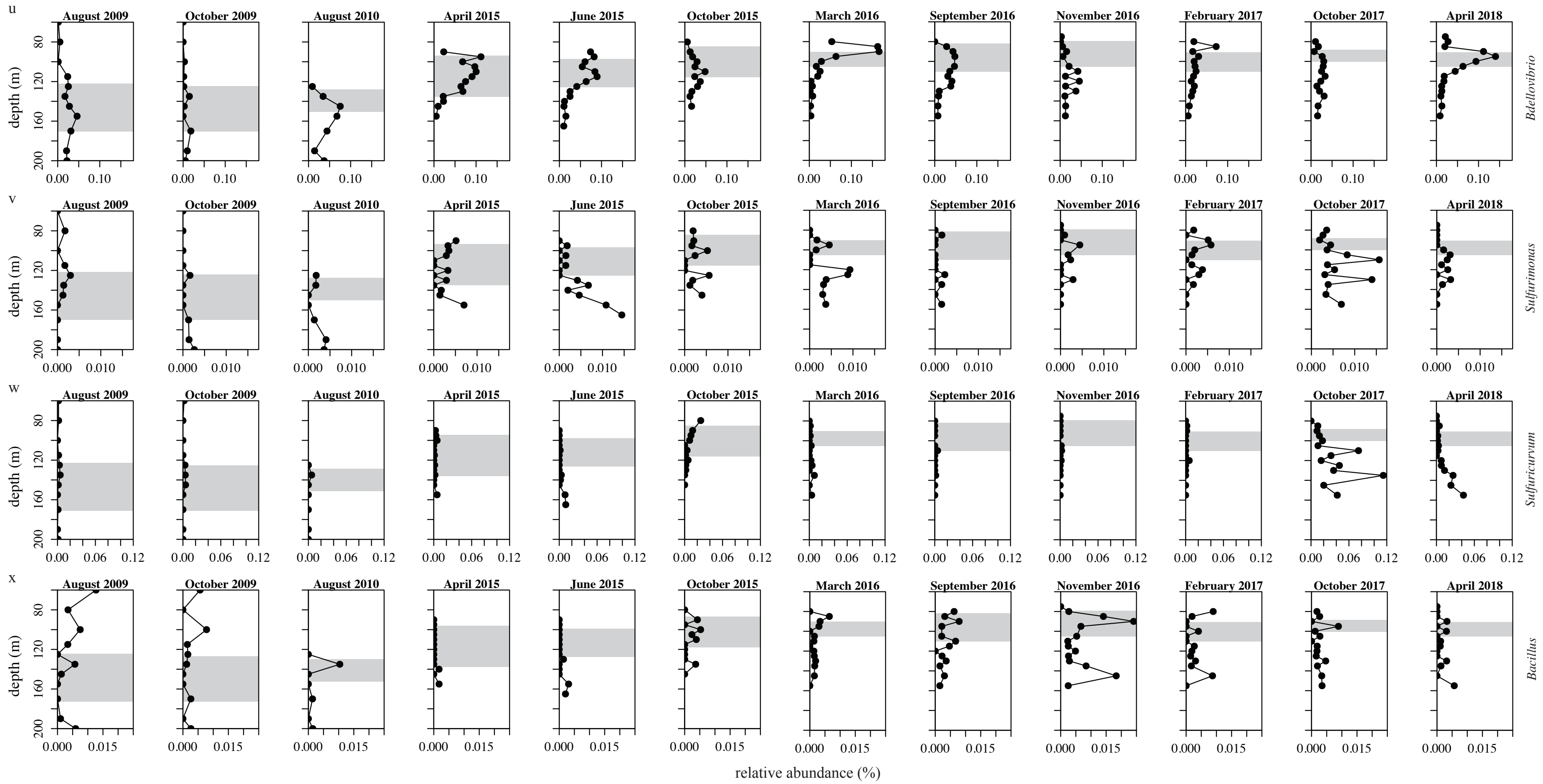


**Figure S10 (continued).** Time series of relative-abundance profiles of amplicon sequencing variants (ASVs) of potential N-reducing bacteria. **u)** Predatory *Bdellovibrio*, **v)** *Sulfurimona*s, **w)** *Sulfuricurvum*, **x)** *Bacillus*. Grey bars represent the RTZ.

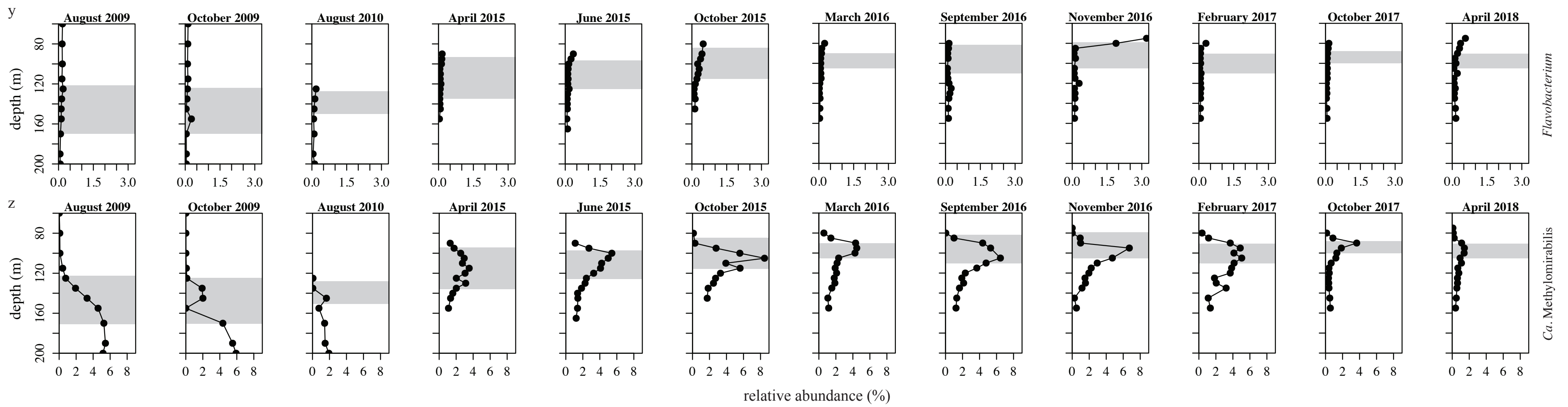


**Figure S10 (continued).** Time series of relative-abundance profiles of amplicon Sequence Variants (ASVs) of potential N-reducing bacteria. **y)** *Flavobacterium*, **z)** *Ca.* Methylomirabilis. Grey bars represent the RTZ.

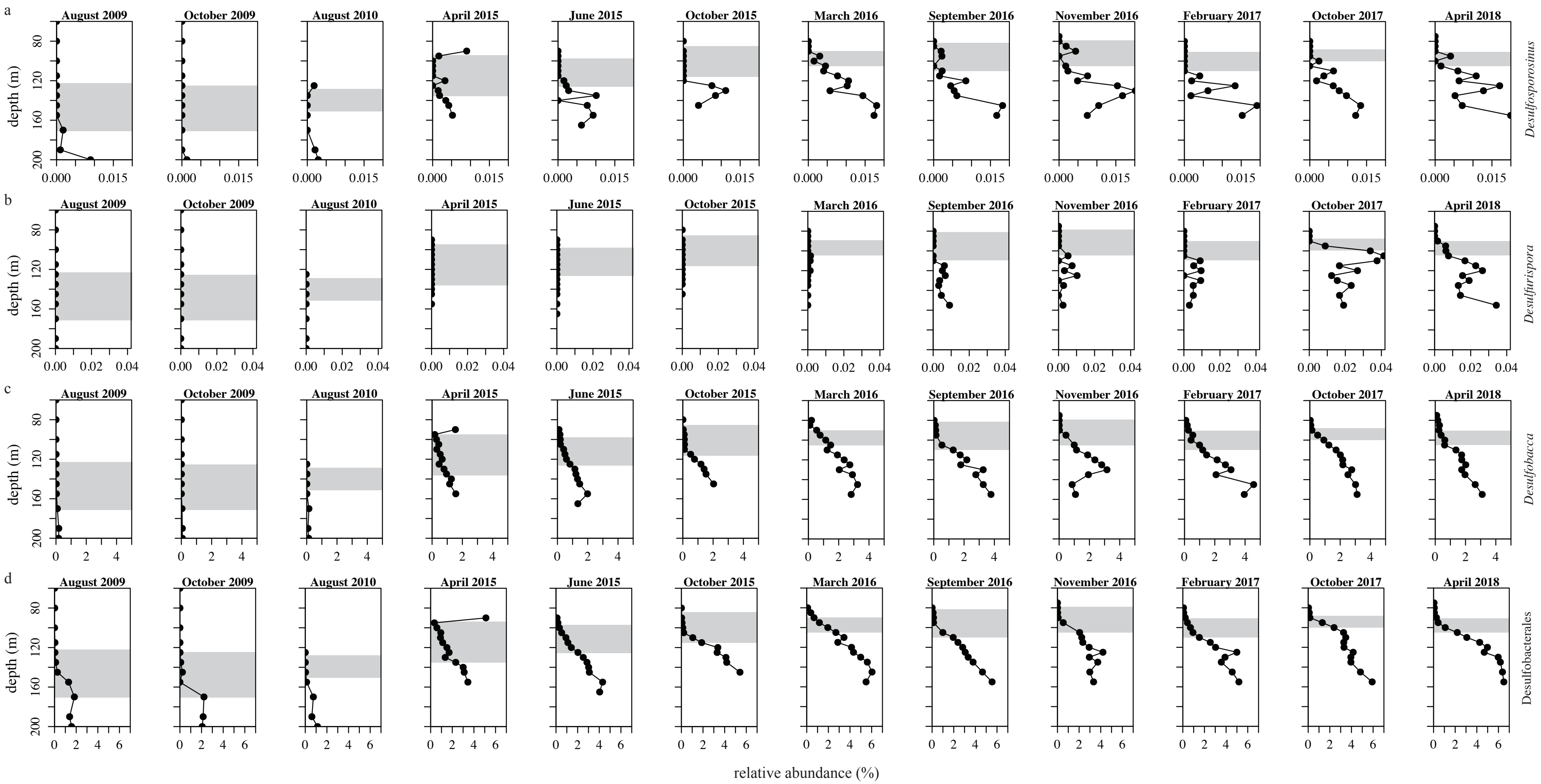


**Figure S11.** Time series of relative-abundance profiles of amplicon sequencing variants (ASVs) of potential S-reducing bacteria. **a)** *Desulfosporosinus*, **b)** *Desulfurispora*, **c)** *Desulfobacca*, **d)** Desulfobacterales. Grey bars represent the redox transition zone (RTZ).

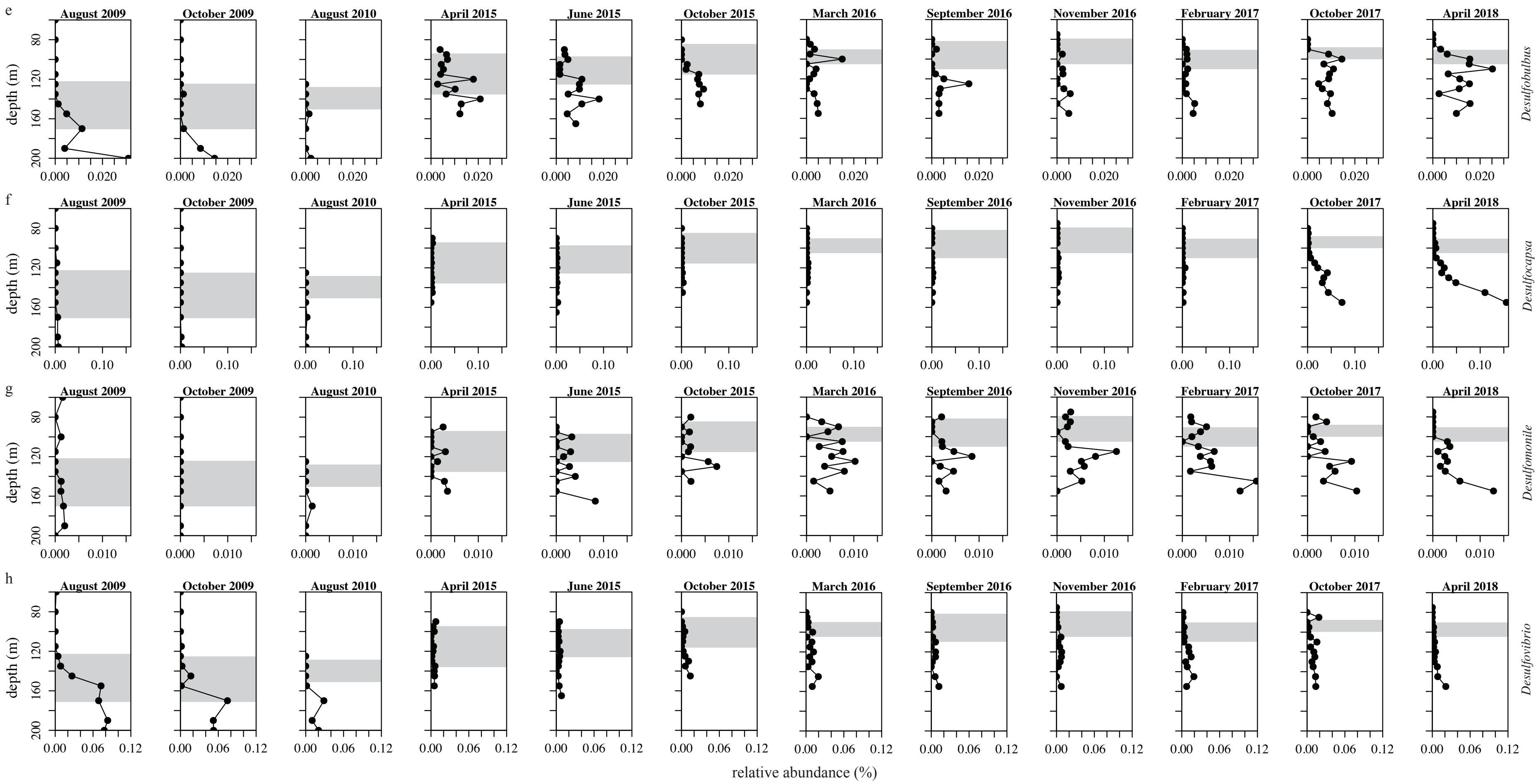


**Figure S11 (continued).** Time series of relative-abundance profiles of Amplicon Sequence Variants (ASVs) of potential S-reducing bacteria. **e)** *Desulfobulbus*, **f)** *Desulfocapsa*, **g)** *Desulfomonile*, **h)** *Desulfovibrio*. Grey bars represent the RTZ.

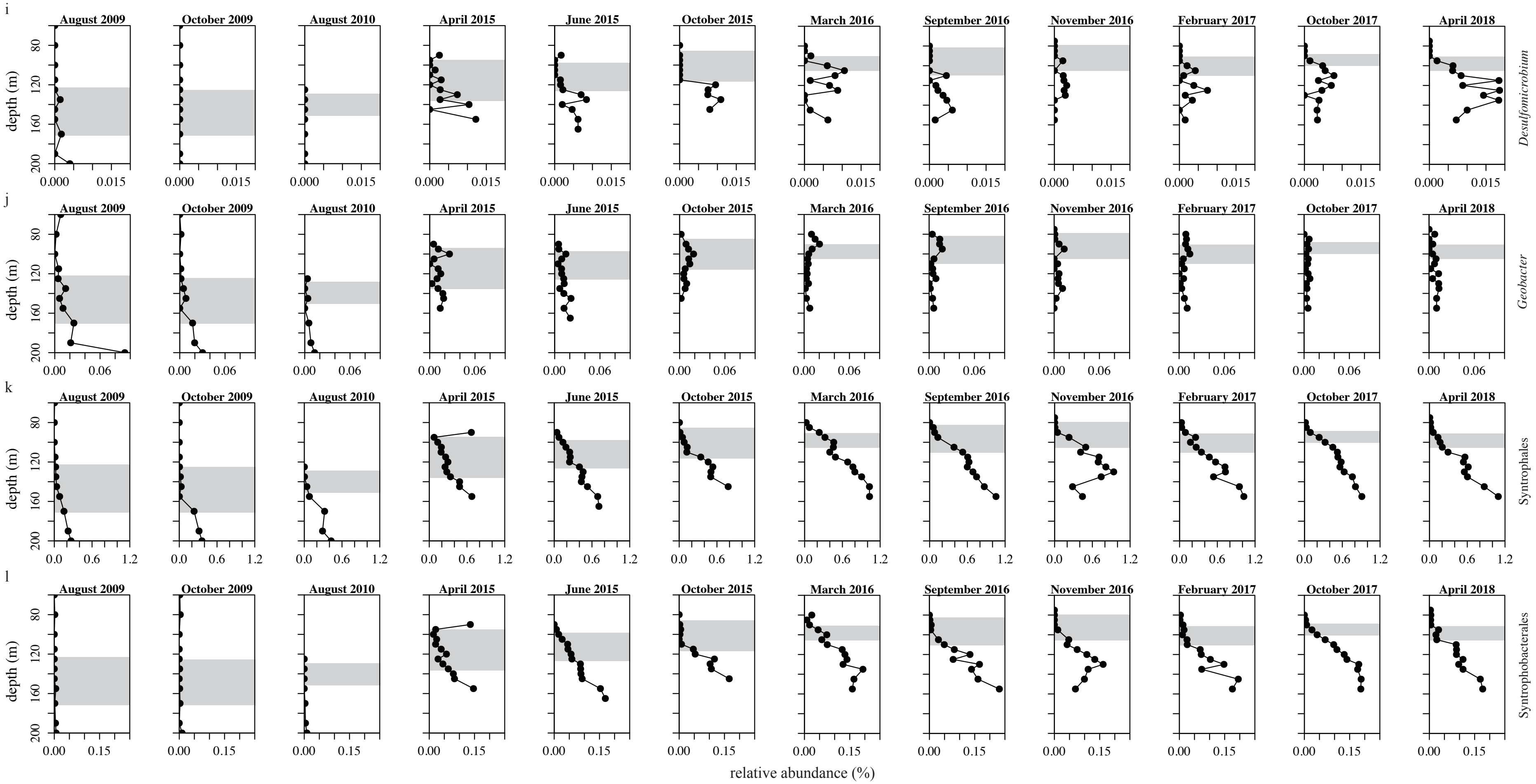


**Figure S11 (continued).** Time series of relative-abundance profiles of Amplicon sequencing variants (ASVs) of potential S-reducing bacteria and syntrophs. **i)** *Desulfomicrobium*, **j)** *Geobacter*, **k)** Syntrophales (secondary fermenters), **l)** Syntrophobacterales (secondary fermenters). Grey bars represent the RTZ.

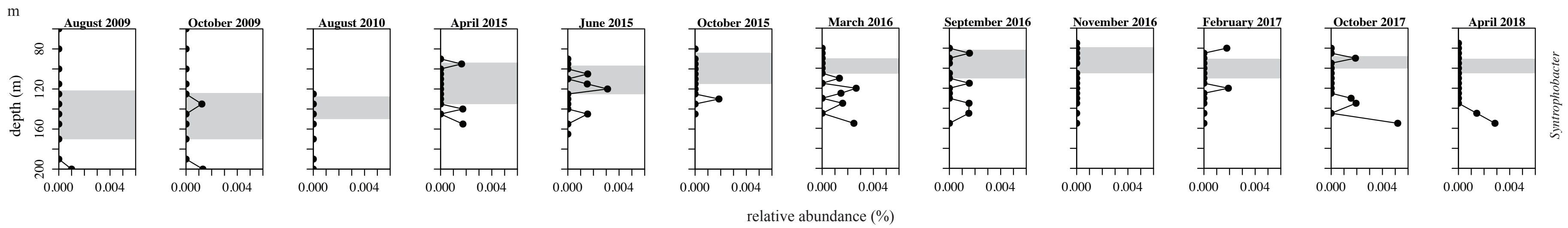


**Figure S11 (continued).** Time series of relative-abundance profiles of amplicon sequencing variants (ASVs) of potential S-reducing bacteria and syntrophs. **m)** *Syntrophobacter* (secondary fermenter). Grey bars represent the RTZ.

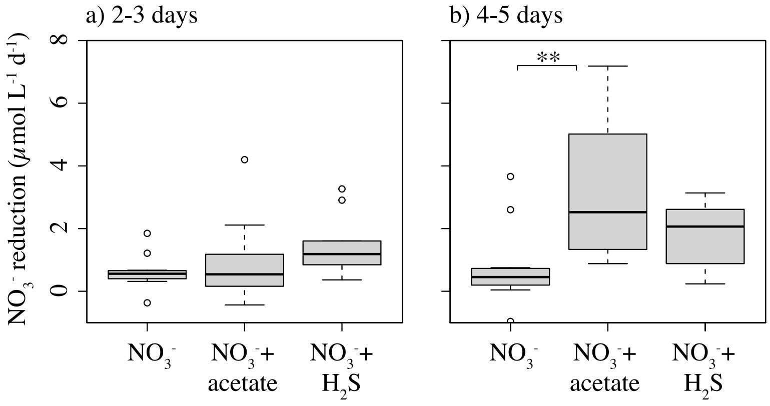


**Figure S12.** $NO_3^-$ reduction rates of incubation experiments with added unlabeled $NO_3^-$; three different treatments are shown ($NO_3^-$ only, $NO_3^-$ and acetate, and $NO_3^-$ and $H_2S$), after **a)** 2-3 days and **b)** after 4-5 days after substrate amendment. Data points originate from incubation experiments performed in November 2016, February 2017, October 2017, and April 2018. Positive values indicate production, negative numbers indicate consumption. Significant differences are marked with * (pairwise t-test; $p < 0.05$ = *, $p < 0.01$ = **, $p < 0.001$ = ***)

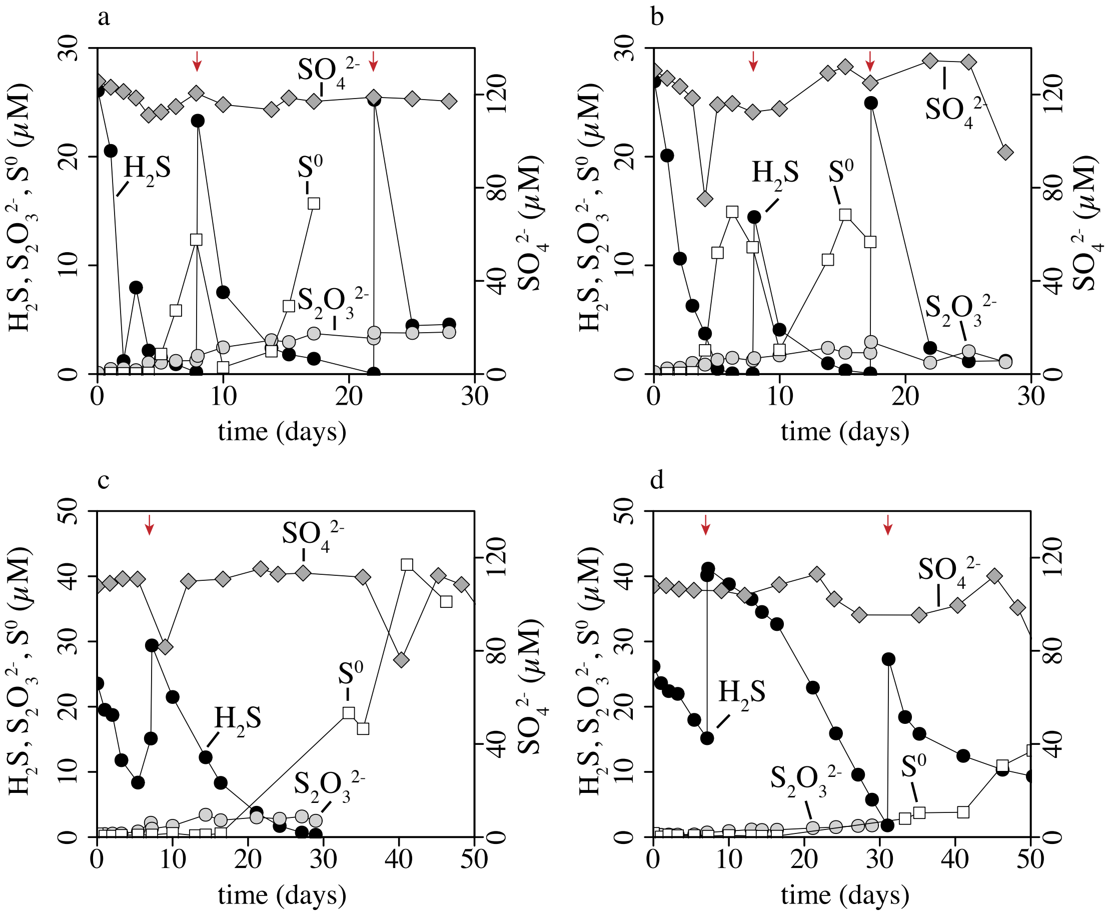


**Figure S13.** Temporal dynamics of $H_2S$, $SO_4^{2-}$, $S^0$, and $S_2O_3^{2-}$ in incubation experiments with unlabeled $NO_3^-$ und $H_2S$ added to anoxic water from 105 m water depth (duplicates in **a** and **b**) and 155 m water depth (duplicates in **c** and **d**) (Lake Lugano North Basin, November 2016). Red arrows indicate time points, where $H_2S$ was resupplied following its intermittent consumption.

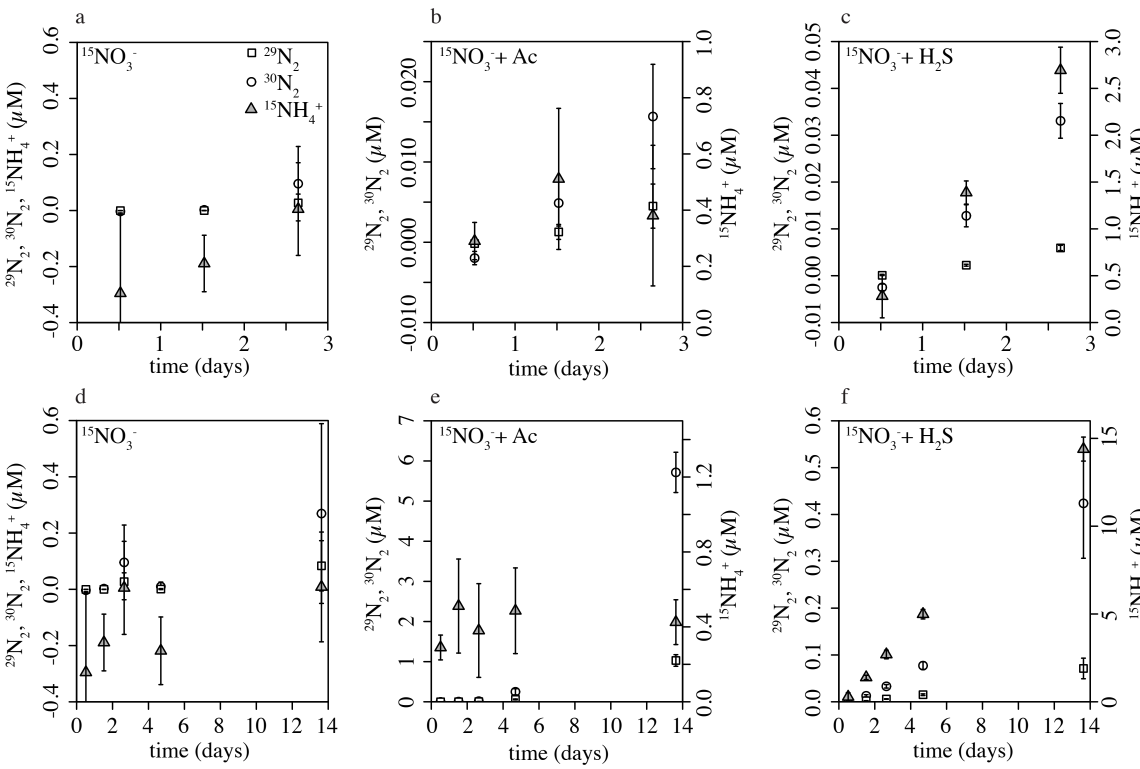


**Figure S14.** Exemplary data of an $^{15}$N-label incubation experiment from April 2018, 95 m. Production of $^{29}N_2$, $^{30}N_2$, and $^{15}N$-$NH_4^+$ after incubation with $^{15}NO_3^-$ **(a, d)**, with $^{15}NO_3^-$ and acetate **(b, e)**, and with $^{15}NO_3^-$ and $H_2S$ **(c, f)**. Panels **a, b, c** show results during 3 days of incubation and panels **d, e, f** show results during 14 days of incubation.

### Supplementary references


Hao, O. J., J. M. Chen, L. Huang, and R. L. Buglass. 1996. Sulfate-reducing bacteria. Crit Rev Environ Sci Technol **26**: 37–41. doi:10.1080/10643389609388489

Haroon, M. F., S. Hu, Y. Shi, M. Imelfort, J. Keller, P. Hugenholtz, and Z. Yuan. 2013. Anaerobic oxidation of methane coupled to nitrate reduction in a novel archaeal lineage. Nature **50**: 2–8. doi:10.1038/nature12375

Jetten, M. S. M., L. Van Niftrik, M. Strous, B. Kartal, J. T. Keltjens, and H. J. M. Op den Camp. 2009. Biochemistry and molecular biology of anammox bacteria. Crit Rev Biochem Mol Biol **44**: 65–84. doi:10.1080/10409230902722783

Kaksonen, A. H., S. Spring, P. Schumann, R. M. Kroppenstedt, and J. A. Puhakka. 2007. *Desulfurispora thermophila* gen. nov., sp. nov., a thermophilic, spore-forming sulfate-reducer isolated from a sulfidogenic fluidized-bed reactor. Int J Syst Evol Microbiol **57**: 1089–1094. doi:10.1099/ijs.0.64593-0

Kits, K. D., M. G. Klotz, and L. Y. Stein. 2015. Methane oxidation coupled to nitrate reduction under hypoxia by the Gammaproteobacterium *Methylomonas denitrificans*, sp. nov. type strain FJG1. Environ Microbiol **17**: 3219–3232. doi:10.1111/1462-2920.12772

Kojima, H., and M. Fukui. 2011. *Sulfuritalea hydrogenivorans* gen. nov., sp. nov., a facultative autotroph isolated from a freshwater lake. Int J Syst Evol Microbiol **61**: 1651–1655. doi:10.1099/ijs.0.024968-0

Kuever, J. 2014. The Family Syntrophobacter, p. 289–299. *In* E. Rosenberg, E.F. DeLong, S. Lory, E. Stackebrandt, and F. Thompson [eds.], The Prokaryotes. Springer.

Martineau, C., F. Mauffrey, and R. Villemur. 2015. Comparative analysis of denitrifying activities of *Hyphomicrobium nitrativorans*, *Hyphomicrobium denitrificans*, and *Hyphomicrobium zavarzinii*. Appl Environ Microbiol **81**: 5003–5014. doi:10.1128/AEM.00848-15

Mustakhimov, I., M. G. Kalyuzhnaya, M. E. Lidstrom, and L. Chistoserdova. 2013. Insights into denitrification in *Methylotenera mobilis* from denitrification pathway and methanol metabolism mutants. J Bacteriol **195**: 2207–2211. doi:10.1128/JB.00069-13

Oswald, K. and others. 2017. *Crenothrix* are major methane consumers in stratified lakes. ISME J **11**: 2124–2140. doi:10.1038/ismej.2017.77

Pandey, C. B., U. Kumar, M. Kaviraj, K. J. Minick, A. K. Mishra, and J. S. Singh. 2020. Science of the Total Environment DNRA: A short-circuit in biological N-cycling to conserve nitrogen in terrestrial ecosystems. Science of the Total Environment **738**: 139710. doi:10.1016/j.scitotenv.2020.139710

Salinero, K. K., K. Keller, W. S. Feil, H. Feil, S. Trong, G. Di Bartolo, and A. Lapidus. 2009. Metabolic analysis of the soil microbe *Dechloromonas aromatica* str. RCB: Indications of a surprisingly complex life-style and cryptic anaerobic pathways for aromatic degradation. BMC Genomics **10**: 1–23. doi:10.1186/1471-2164-10-351

Shao, M. F., T. Zhang, and H. H. P. Fang. 2010. Sulfur-driven autotrophic denitrification: Diversity, biochemistry, and engineering applications. Appl Microbiol Biotechnol **88**: 1027–1042. doi:10.1007/s00253-010-2847-1

Shapleigh, J. P. 2013. Denitrifying Prokaryotes, p. 405–425. *In* E. et Al Rosenberg [ed.], The Prokaryotes – Prokaryotic Physiology and Biochemistry. Springer-Verlag Berlin Heidelberg.

Sparacino-Watkins, C., J. F. Stolz, and P. Basu. 2014. Nitrate and periplasmatic nitrate reductases. Chem Soc Rev **43**: 676–706. doi:10.1039/c3cs60249d

Willems, A. and others. 1989. *Hydrogenophaga*, a new genus of hydrogen-oxidizing bacteria.

Xia, Z., Q. Wang, Z. She, M. Gao, Y. Zhao, L. Guo, and C. Jin. 2019. Nitrogen removal pathway and dynamics of microbial community with the increase of salinity in simultaneous nitrification and denitrification process. Science of the Total Environment **697**: 1–10. doi:10.1016/j.scitotenv.2019.134047

Yang, Y. and others. 2021. The evolution pathway of ammonia-oxidizing archaea shaped by major geological events. Mol Biol Evol **38**: 3637–3648. doi:10.1093/molbev/msab129
